# Flexibility and sensitivity in gene regulation out of equilibrium

**DOI:** 10.1101/2023.04.11.536490

**Authors:** Sara Mahdavi, Gabriel L. Salmon, Patill Daghlian, Hernan G. Garcia, Rob Phillips

## Abstract

Cells adapt to environments and tune gene expression by controlling the concentrations of proteins and their kinetics in regulatory networks. In both eukaryotes and prokaryotes, experiments and theory increasingly attest that these networks can and do consume biochemical energy. How does this dissipation enable cellular behaviors unobtainable in equilibrium? This open question demands quantitative models that transcend thermodynamic equilibrium. Here we study the control of a simple, ubiquitous gene regulatory motif to explore the consequences of departing equilibrium in kinetic cycles. Employing graph theory, we find that dissipation unlocks nonmonotonicity and enhanced sensitivity of gene expression with respect to a transcription factor’s concentration. These features allow a single transcription factor to act as both a repressor and activator at different levels or achieve outputs with multiple concentration regions of locally-enhanced sensitivity. We systematically dissect how energetically-driving individual transitions within regulatory networks, or pairs of transitions, generates more adjustable and sensitive phenotypic responses. Our findings quantify necessary conditions and detectable consequences of energy expenditure. These richer mathematical behaviors—feasibly accessed using biological energy budgets and rates—may empower cells to accomplish sophisticated regulation with simpler architectures than those required at equilibrium.

**Significance Statement:** Growing theoretical and experimental evidence demonstrates that cells can (and do) spend biochemical energy while regulating their genes. Here we explore the impact of departing from equilibrium in simple regulatory cycles, and learn that beyond increasing sensitivity, dissipation can unlock more flexible input-output behaviors that are otherwise forbidden without spending energy. These more complex behaviors could enable cells to perform more sophisticated functions using simpler systems than those needed at equilibrium.

## Introduction

Gene regulation—to which biology owes much of its exquisite sophistication (1)—is replete with network architectures that allow (and credibly depend on) nonequilibrium (2–5). To adapt to environmental cues, cells often dynamically tune concentrations of transcription factors (6) or inducers as their available control variables. This biochemical control adjusts the probabilities of cellular states by regulating rate constants that depend on the transcription factor or effector. The majesty of biological regulation is often woven from the specific shapes of these input (transcription factor concentration) to output (average steady-state gene expression) relationships. As crucial means by which cells adapt their physiology and defy environmental variation, these induction curves also promise to trace design principles that illuminate how spending biochemical energy empowers the very dynamism and fidelity of the living. Stubborn (7, 8)—yet increasingly well-measured (9–11)—energetic budget mismatches and mysteries about what biochemical energy expenditures accomplish place fresh urgency on deciphering how dissipation modifies gene regulation.

How can nonequilibrium relieve fundamental constraints on physiological adaptation, or enhance the flexibility of cellular behavior? To confront this question, here we examine the output behavior of among the simplest closed systems capable of breaking equilibrium using basic reactions pervasive in biology: a cycle of four states. This system can represent the dynamic behaviors of genetic transcription executed by RNA polymerase (RNAP) and regulated by a transcription factor acting as a control variable (Fig. 1A).

**Fig. 1.**
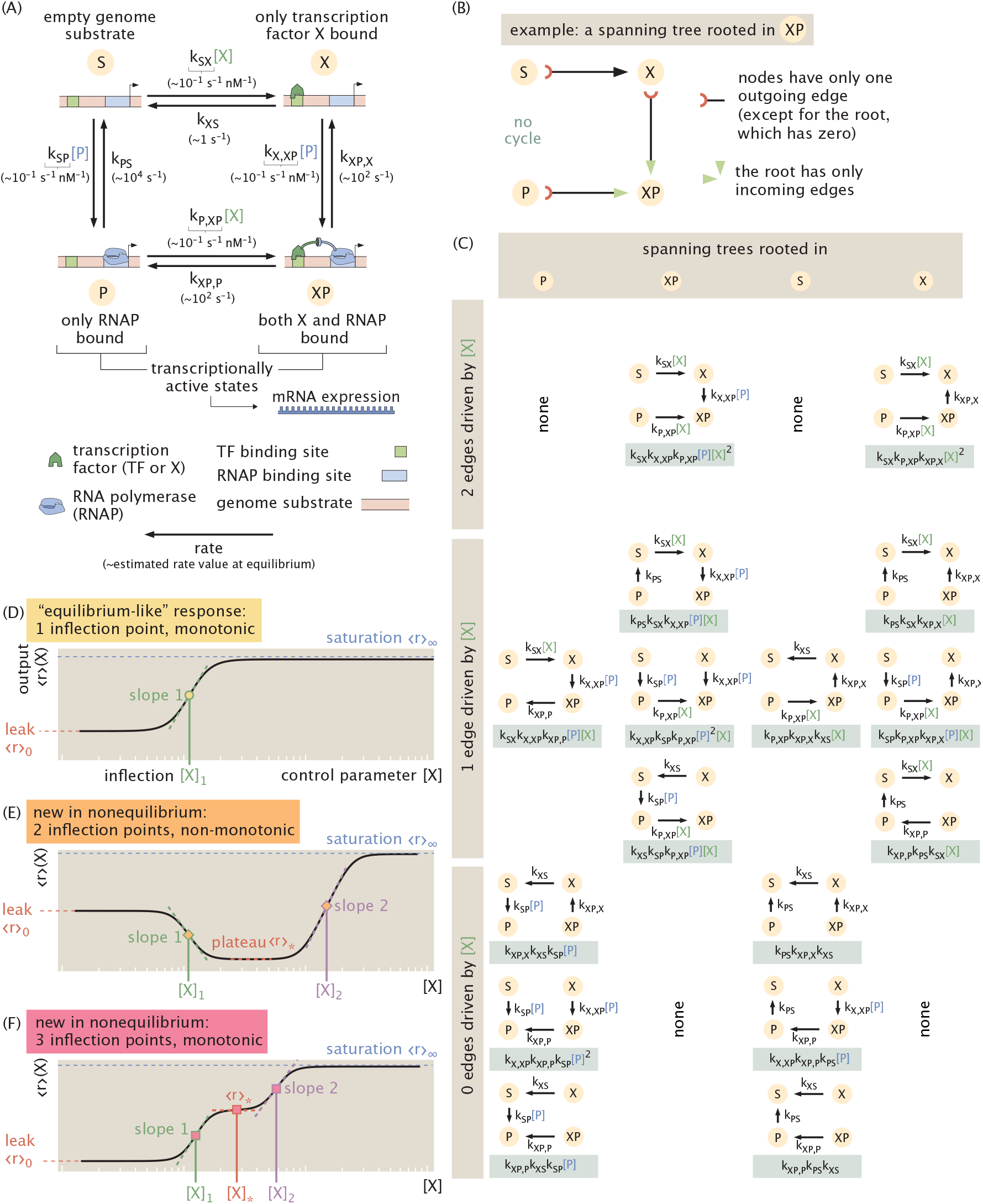
Structure and (non)equilibrium response of a four-state cycle, a fundamental gene-regulatory motif. (A) A square cycle of four-states emerges when up to two molecules (such as a transcription factor *X* and polymerase *P*) can bind to a common substrate (say a genome). Output observables ⟨*r*⟩ are linear combinations of the state probabilities; for instance, mRNA production scales with the probabilities of transcriptionally active states where polymerase is bound to the genome (states *P* and *XP*). These outputs vary with the control parameter [*X*], here schematized as the concentration of a transcription factor. (B) An example of a spanning tree (rooted in state XP) like those that define steady-state probabilities via the Matrix Tree Theorem. (C) All 16 directed, rooted spanning trees of the four-state cycle in (A): trees are grouped by the root state (in columns) and by how many participating edges depend on the control parameter *X* (in rows). As guaranteed by the Matrix Tree Theorem, the steady-state probability of any state—in or out of equilibrium—is given by the sum of the weights of these spanning trees, introducing up to a quadratic dependence in *X* in any output, as represented by Eq. 1. (D-F) Three universal output behaviors (*regulatory shape phenotypes*) can result from this architecture. A monotonic “equilibrium-like” sigmoidal output (D) manifests a Hill-like or MWC-like response, behavior familiar from equilibrium thermodynamic models. However, exclusively out of equilibrium, new multiply-inflected regulatory shape phenotypes become possible. Under drive, outputs can (E) vary non-monotonically and reach two inflection points with the control parameter; or show three inflection points and vary monotonically (F). These richer phenotypes show a wider set of properties that characterize each curve: these include the “leak” value of the observable when the control variable is absent (⟨ *r* ⟩ _0_ = ⟨ *r* ⟩ ([*X*] = 0), in orange; the saturation asymptotic limit as the control variable is maximally present (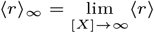 ; in light blue); the observable’s values at intermediate plateau regions (⟨*r*⟩ ^*^; in red); and slopes 1 and 2 at inflection points [*X*]_1_ and [*X*]_2_ when they are defined (in green and purple, respectively).

Given their simplicity, equivalents of the system in Fig. 1A have enjoyed earlier study in guises such as enzymatic control (12); remodeling of nucleosomes (5); and other settings in transcription (13, 14). In this work, we use tools from graph theory (15, 16) to explore the full space of transcriptional steady-state outputs available for this system under different energetic drives, compared to equilibrium control. We find that all equilibrium responses must be monotonic (with one inflection point) as a function of control variables, such as the concentration of transcription factor, measured in a conventional logarithmic scale. In contrast, we discover that nonequilibrium models can exhibit three types of output: an “equilibrium-like,” monotonic response with one inflection point, potentially displaced from equilibrium; a new —but still-monotonic—shape with three inflection points; and a new, surprising non-monotonic shape with two inflection points, where, for instance, increasing a control variable can change its effect from repression to activation. Combining analytical and numerical analysis, we globally bound the maximal sensitivities of transcriptional responses. Demonstrating that these mathematical behaviors are feasible to access within biological energy expenditures around typical rates, we systematically analyze the impact of breaking detailed balance along each transition rate. This analysis establishes design principles for optimizing sensitivity and unlocking dramatic behaviors that are especially prone to implicate nonequilibrium in measurements.

These broader, multiply-inflected transcriptional responses unlocked by nonequilibrium could be harnessed to achieve useful physiological functions. Our findings illustrate surprising regularity visible from graph theoretic tools, and explicate how even primordial biological networks operating out of equilibrium can rival the regulatory sophistication of (plausibly) larger, slower networks at equilibrium.

## Results

### A model of a pervasive gene regulatory motif

At steady-state, a system is in *equilibrium* (or, equivalently, at *detailed balance*) if, for all pairs of states (*i, j*), the probability flux *k*_*ij*_ *p*_*i*_ state *j* equals the flux *k*_*ji*_*p*_*j*_ into state *i*, where *p*_*i*_ is the probability of state *i* and *k*_*ij*_ is the rate of transitions from state *i* to *j*. Otherwise, the system is out of equilibrium and requires energetic dissipation to sustain the system’s steady-state. For systems closed to external material inputs, nonequilibrium steady-states can only be achieved with systems that contain at least one cycle; linear or branched architectures at steady-state must be at equilibrium (see Supporting Information (SI), §1B: *Closed steady-state systems are either equilibrium or cyclic* and (17, 18)). A single cycle is thus the simplest closed setting where the intriguing new consequences of nonequilibrium become possible.

A cycle of four states emerges naturally from up to two molecules binding or unbinding to a substrate. When the substrate is a promoter site on the genome *S*, one molecule is RNA polymerase *P*, and the second molecule is a transcription factor protein *X* that can enhance or impede polymerase binding to the genome, the resulting cycle captures transcriptional regulation. Specifically, the four states represent the empty site of the genome substrate (“S”); the genome substrate bound to the transcription factor only (“X”); to the polymerase only (“P”); or to both (“XP”). Figure 1A illustrates this central, motivating setting. (Note that the transcription factor and polymerase concentrations [*X*] and [*P*] do not affect whether the system is in or out of equilibrium, and can be tuned while separately maintaining any extent of disequilibrium—see SI, *§*1C: *The cycle condition relates a ratio of rate constants to (non)equilibrium*.)

This square cycle of states pervades gene regulation. In one of the widest experimental surveys of prokaryotic regulatory motifs yet available—mapping over one hundred new regulatory interactions in *E. coli*—motifs regulated by a single transcription factor, which can often manifest a four-state cycle, were found to be the most common regulated architectures (19), joining similar reports from aggregated databases (20). These cyclic architectures contrast the more commonly studied motif of simple repression that cannot break detailed balance (see SI, §1B: *Closed steady-state systems are either equilibrium or cyclic*) (1, 6, 19–21). The four-state cycle finds widespread examples or structural-equivalents in eukaryotic gene regulation as well (5, 13, 22, 23). Eukaryotic gene expression is a setting where explicit ATP-consumption is especially plausible (3, 4) yet poorly understood (2, 8, 13).

Kinetic measurements often justify the assumption that transcription factors bind and unbind with genomes quickly relative to transcription by polymerase. This separation of timescales makes macroscopic gene expression proportional to the steady-state probability of finding the system in transcriptionally-active microstates. (We precisely validate this assumption for our setting using plausible transcriptional rates in the SI, §2C: *Biologically, timescales are plausibly separated enough that transcription is well represented by small Markov chains*.)

We note that the average gene production rate ⟨*r*⟩ _mRNA_, proportional to gene expression, is a typical and crucial output of interest. This response grows with the net probability that the polymerase is bound, ⟨*r* _mRNA_ ⟩= *r*(*p*_*P*_ + *p*_*XP*_), where *r* is the transcription rate once the polymerase is bound, *p*_*p*_ is the probability of the state *P* where just the polymerase is bound, and *p*_*XP*_ is the probability of the state *XP* where both polymerase and transcription factor are bound.

However, other outputs (that depend on other states) may also be biologically or experimentally significant. For instance, the localization of the transcription factors themselves to the genome (to recruit other co-factors or epigenetic modifications) can shape biological function independent of the polymerase, e.g. invoking the probability *p*_*X*_. We accommodate the breadth of these possible outputs by studying how any (nonnegative) linear combination 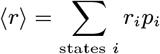 of state probabilities varies with the transcription factor concentration *X* as a control variable, where *r*_*i*_ gives the potency of the *i*th state. These different outputs and problem settings are captured by adopting particular {*r*_*i*_}, but as we will now see, all are subject to universal behavior.

### Nonequilibrium steady-state output responses

To explore how these input-output responses operate away from equilibrium, we cannot depart from the equilibrium statistical mechanical models, which use the thermodynamic energies of each state to calculate their probabilities, that suffice for acyclic architectures (such as simple repression) (1, 6, 24–26). Instead, we embrace a fully kinetic description (also known as a chemical master equation or continuous-time Markov chain) based on transitions between states. A large increase in complexity and the number of parameters typically accompanies this generalization. Fortunately, these dynamics admit a beautiful and powerful correspondence to graph theory that helps tame this complexity (15). Our guide is the Matrix Tree Theorem, which gives a simple diagrammatic procedure on a network’s structure to find stationary probabilities (see Methods and SI, §2D: *Deriving the universal form: The Matrix Tree Theorem on the square graph yields a ratio of quadratic polynomials*). In brief, the Matrix Tree Theorem asserts that at steady-state, the probability of any state is proportional to the sum of products of rate constants over all spanning trees rooted in that state. Here, a *spanning tree* is a (directed) subset of edges on the graph of states that collectively visits every state exactly once, privileging a *root* state, which has no outgoing edges. Figure 1B illustrates these requirements with an example of a rooted spanning tree in our four-state graph. Counting all sixteen rooted spanning trees of the four-state transcriptional system (Figure 1C) and deploying the

Tree Theorem explains how probabilities must vary with the transcription factor control parameter [*X*]. Depending on the root (separated by column in Figure 1C), each spanning tree carries two edges that depend on [*X*] (top row of Fig. 1C); one edge (middle row, Fig. 1C); or no [*X*] −dependent edges (bottom row, Fig. 1C). This structure yields statistical weights with up to quadratic scaling with [*X*]. Hence we find that the form of any output function ⟨*r*⟩, in or out of equilibrium, is a ratio of quadratic polynomials in [*X*],

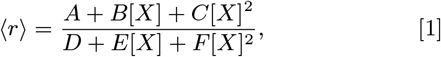

where the coefficients *A, B, C, D, E* and *F* are sums of subsets of (weighted) directed spanning trees carrying various [*X*]-dependencies (see SI, §2D: *Deriving the universal form: The Matrix Tree Theorem on the square graph yields a ratio of quadratic polynomials*). The denominator, the sum of all rooted spanning trees and hence also a quadratic polynomial, serves as a normalizing factor that converts statistical weights to probabilities and represents a nonequilibrium partition function.

Note that while we derived the output form Eq. 1 using the particular choice of [*X*]-dependent arrows appropriate for this transcriptional setting, the same formalism can treat many other control parameters that appear quite (structurally or biologically) distinct from these details, such as a concentration of another internal molecule (for instance polymerase, [*P*]) or an external molecule (for instance explicit drive by [*AT P*]). The SI, §2H: *Driving different arrows in the square graph can still yield a ratio of quadratic polynomials* gives some further examples of different placements of controlled edges that still produce a network output with the functional form of Eq. 1, and therefore remain precisely addressable by the analysis of this paper. Other outputs will require a fresh application of the Matrix Tree Theorem and new analysis but benefit from the same framework.

### Equilibrium output curves are constrained and always sigmoidal

Eq. 1 describes all induction curves, in or out of equilibrium, produced by this four-state transcriptional system. When detailed balance does hold, this equation becomes equivalent to thermodynamic statistical-mechanical models (as it must). We explain algebraic correspondences to thermodynamic models, like those communing with earlier transcriptional experiments (6, 26), in the SI, §G.3, *Validating consilience between kinetic and thermodynamic viewpoints*. Importantly, we find that the equilibrium condition demotes any observable output to the simpler form of a ratio of *linear* polynomials in [*X*], namely

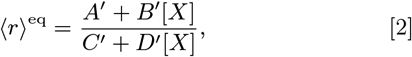

for constants {*A*^′^, *B*^′^, *C*^′^, *D*^′^ } set wholly by thermodynamic parameters (see the SI, §G.1: *Demotion of responses to a (monotonic) ratio of linear polynomials at equilibrium*). Not coincidentally, this functional form formally reproduces or evokes the Hill induction, Michaelis-Menten, Langmuir-binding, Monod-Wyman-Changeux, or two-state Fermi function forms from the equilibrium statistical mechanics of binding commonly used to model and fit induction curves in natural (6, 27) or synthetic (28) settings. This equilibrium curve is paradigmatic of our biochemical intuition—sigmoidally saturating, with one point of inflection, with respect to transcription factor concentration [*X*] in a conventional logarithmic scale (see Fig. 1A and the SI, §2E: *Discussion on observable conventions: the logarithmic control variable*).

### New regulatory shape phenotypes unlocked by nonequilibrium

How much more complex is the regulation realizable by nonequilibrium outputs ⟨*r*⟩ (Eq. 1), compared to that of their equilibrium special case, ⟨*r*⟩ ^eq^ (Eq. 2)? To reach the qualitative essence of this question, we first investigate the possible *shapes* of the output curve. Specifically, we monitor the output’s changes in concavity with respect to the control parameter. We postpone comment on the characteristic positions and scales of output curves—any shifts in their horizontal position (*viz*. any characteristic concentration scales) or vertical expanses (*e*.*g*. maximally-induced responses)—until shortly.

Neglecting scales and shifts allows us to collapse the general, six-parameter output curve of Eq. 1 to a normalized function of just two emergent shape parameters,

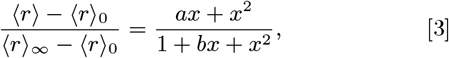

Here, the emergent shape parameters *a* and *b* are complicated functions of the coefficients in Eq. 1 (and hence of underlying rate constants), and *x* is the governing concentration [*X*] measured in terms of a characteristic concentration scale (all defined in the SI, §2F: *Collapse of eight parameters into two emergent fundamental shape parameters* (*a, b*)). The values ⟨*r*⟩_0_ ≡ ⟨*r*⟩ ([*X*] = 0) and 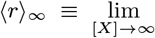 and *saturation* (maximally-induced) responses; we return to these values in the following subsections. This representation preserves the concavity of the response function, allowing us to explore shapes and quantitative features in a two-dimensional space more efficiently and comprehensively than possible in the space of the eight rates.^*^

Harnessing this collapsed representation, we discover that all output curves assume just three different universal shapes (see Methods & SI, §2I: *Any averaged observable ⟨r⟩ has zero, one, two, or three inflection points, with varying monotonicity*).^†^ First, the output can be sigmoidal and monotonic, with a single inflection point, with respect to the control parameter (on a log scale), recalling the shape of the equilibrium response (Fig. 1D). Uniquely out of equilibrium, however, two additional multiply-inflected response shapes become possible. Under energy expenditure, outputs can become nonmonotonic and show two inflection points (Fig. 1E), or remain monotonic with three inflection points (Fig. 1F), with respect to the log of the control parameter. Responses with three inflections are always shaped as depicted in Fig. 1F: maximally steep at the first and third inflection points, but minimally steep at the second inflection point.

Clearly, these nonequilibrium curves are marked departures from simple equilibrium-like sigmoids, but betray a remarkable parsimony and regularity, given that they describe all departures from equilibrium for any rate parameter values. These three regulatory behaviors can pose different physiological implications for an organism; admit distinct quantitative constraints on sensitivity (as we will soon see); and require different conditions on underlying rate constants to be reached. In view of their categorical differences, we refer to these possible shapes as *regulatory (shape) phenotypes*.^‡^

### Quantitative traits of response functions

Beyond their shape phenotypes, regulatory output curves affect the destiny of organisms through their quantitative traits. Further, engineering responses with desirable properties—e.g. high gain, low background, tight affinity, and high sensitivity with respect to an inducer—is a critical and intensely-pursued design goal of synthetic biology (28, 30); such traits can also themselves reveal the presence of nonequilibrium, as with the presence of ultrasensitivity (31).

These properties include the *leakiness* ⟨*r*⟩_0_ ≡ ⟨*r*⟩([*X*] = 0) and *saturation* 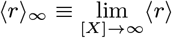 defined earlier; and the *dynamic range* (difference between the leakiness and the saturation, |⟨*r*⟩_∞_− ⟨*r*⟩ _0_ |). In addition, the response’s maximum *sensitivity* with respect to the input (often characterized by a suitable *logarithmic sensitivity*, sharpness, or effective Hill coefficient)—and the level(s) of input where this maximal sharpness occurs, namely the location(s) of the inflection point(s)—are crucial determinants of regulatory adaptability. For equilibrium-like binding curves, just one input level (the single inflection point, localizing maximal sensitivity) suffices to define the horizontal position of the curve. This inflection point is often linked with the input needed to induce a response about halfway between leakiness and saturation, denoted the *EC50*. However, the new complexity of nonequilibrium outputs introduces additional characteristic concentration scales (at each point of inflection) and their associated locally-extremal sensitivities.

Does spending energy enable finer control over these quantitative traits, beyond growing their number? In fact, as we now discuss, only some traits are given extra adjustability by spending energy.

### Leakiness, saturation, and EC50 are tunable at equilibrium

Without the transcription factor, the system cannot be found in any microstate that involves it, collapsing four states into just the two {*S, P*} states. This pair of states forms an acyclic graph, so these steady-state probabilities must show detailed balance (i.e. are set purely thermodynamically). Thus, leakiness ⟨*r*⟩ _0_, determined exclusively by *S* and *P* states, can be adjusted freely while maintaining detailed balance. Analogously, when the transcription factor concentration is saturating ([*X*] →∞), the system is never found in the two microstates without the transcription factor, again admitting an orthogonal description of a balance between two states, now {*X, XP* }. Hence, saturation ⟨*r*⟩_∞_ is also freely adjustable at equilibrium. These leakiness and saturation values are independently adjustable by two separate energy parameters—the binding energies of the polymerase to the genome when the transcription factor is absent or present, respectively. At equilibrium, once the leakiness and saturation are fixed by energy parameters, the response’s maximal sensitivity (slope at the inflection point) is predetermined and no longer tunable, as revealed by its algebraic dependencies (see SI §G.2). In contrast, while the location of the governing inflection point depends on these two energy parameters, it can also be tuned—remaining at equilibrium—using another energy parameter (the binding energy between the transcription factor and genome). (See SI, §G.2:*Leakiness, saturation, and EC50 are tunable at equilibrium* for details.)

### Nonequilibrium control of sensitivity obeys shape-dependent global bounds

Out of equilibrium, the sensitivity of responses enjoys greater adjustability. Specifically, the diversity of input-output curves accessible under drive motivate us to assess sensitivity by a suitably normalized slope *s*([*X*]), defined by

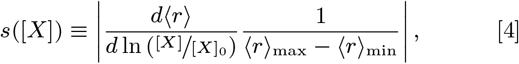

where 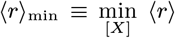 and 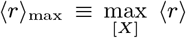 are the extremal values of the observable over all [*X*], and [*X*]_0_ is an arbitrary characteristic concentration scale ensuring dimensional consistency. For monotonic curves, the maximum ⟨*r*⟩ _max_ and minimum ⟨*r*⟩ _min_ responses are necessarily the uninduced leakiness ⟨*r*⟩ _0_ and the maximally-induced saturation ⟨*r*⟩_*∞*_ (or vice-versa), whereas for nonmonotonic responses with two inflections, the maximal and minimal responses can occur at intermediate finite values of [*X*].

This normalized sensitivity *s*([*X*]) is directly related to familiar measures such as the logarithmic sensitivity and the effective Hill coefficient, but more naturally describes sensitivities of nonmonotonic phenotypes using finite values (see SI, §J: *New bounds on nonequilibrium sensitivity*).

By combining wide numerical sampling, symbolic inequality solving, and analytical arguments (see SI, §J: *New bounds on nonequilibrium sensitivity*), we investigated the maximal normalized sensitivity *s*([*X*]) any response curve can exhibit for the four-state system across its three possible shape phenotypes. We discovered that sensitivity is tightly bounded above *and* below by precise finite limits; these limits vary by phenotype. Figure 2 summarizes these bounds, visualized by how normalized and centered response curves 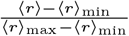 behave around inflection points of maximal slope. Equilibrium response curves always show a normalized sensitivity of exactly one-fourth. Out of equilibrium, singly-inflected response curves can increase this maximal sensitivity up to one-half, or *decrease* maximal sensitivity below the equilibrium value to a numerical value of about 0.158. (We lack a coherent explanation for this curious numerical lower bound, but verified it by precise symbolic inequality solving; see SI, §J). Driven curves with two inflection points all have maximal sensitivity of *at least* the equilibrium level of one-fourth, but up to one-half. Driven curves with three inflection points all show maximal sensitivity of *at most* the equilibrium level of one-fourth, and at least a sensitivity of one-eighth.

**Fig. 2.**
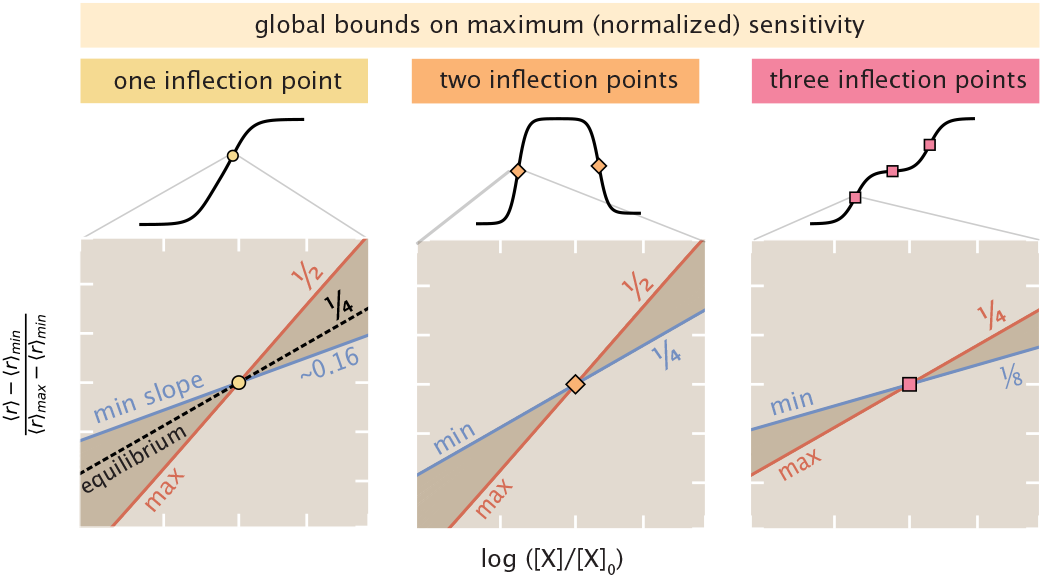
Global bounds, in or out of equilibrium, restrict maximal (normalized) response sensitivity (with respect to input concentrations [*X*] on a log scale). Plotted are normalized responses 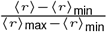 near points of inflection that maximize slope, separated by shape phenotype. When the output has one inflection point (left), the maximal sensitivity is bounded between a minimum of 0.158 (blue line) and a maximum of 1*/*2 (red line) for any set of rate values or any dissipation; this subsumes the equilibrium case, whose normalized sensitivity is fixed at 1*/*4 (black dotted line). When the output has two inflections (middle), the maximal sensitivity is bounded between 1*/*4 and 1*/*2. When the output has three inflections (right), the maximal sensitivity is bounded between 1*/*8 and 1*/*4.

Cast in terms of the *raw* maximal sharpness *d ⟨r⟩ /d* ln (^[*X*]^*/*_[*X*]0_) of each response curve, these bounds report that raw maximal sharpness is always between one eighth and one half of the distance between the maximum and minimum responses per *e*≈ 2.7-fold increase in the concentration [*X*]. We stress that these bounds on sensitivity, in terms of the observed ⟨*r*⟩ _min_ and ⟨*r*⟩ _max_, are tighter quantitative constraints than bounds merely in terms of the maximal or minimal potency values 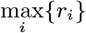 or 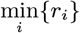 that any microstate of the system can show, as can be connected to recent, related upper bounds (29). This follows since in general the extrema of the *average* observable response curve over all [*X*] are usually more restricted than the most extreme potencies over microstates (namely, 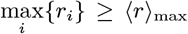 and 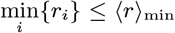). (See SI, §J.4: *General upper bound on a related, differently-normalized slope*.)

These findings emphasize that network architecture and dissipation are not the only hard global constraints that bound sensitivity. The global shape of the response curve further categorically constrains the possible sensitivity. This relationship is potentially biologically relevant: for instance, it is impossible for an organism regulated by the square-graph transcriptional motif to achieve both a triply-inflected output curve and a normalized sensitivity greater than that at equilibrium. This represents a tradeoff between the shape complexity of a response and its maximal sensitivity.

### Breaking detailed balance along each edge

Our foregoing analysis has been mathematically general. That is, the constrained shapes and bounds on sensitivity hold for any response following Eq. 1, over all rate constant values and energetic dissipations. These constraints also apply even—as previously noted—if the response is produced by a different underlying graph architecture than the particular transcriptional motif shown in Fig. 1A, as long as the graph still yields spanning trees that depend up to quadratically on the control variable. Just because multiply-inflected or adjustable response curves are mathematically possible, however, does not establish that they are biologically plausible. To assess whether these behaviors can be accessed using physiologically-plausible amounts of energy expenditure or typical biological rates, we now specialize to the plausible particulars of transcription as in Fig. 1A. In the remainder of this paper, we quantify the extent of dissipation sustaining a nonequilibrium steady-state by focusing on the free energy Δ*μ* coupled to the system, with units of *k*_*B*_*T* or Joule; we refer to this quantity as the *nonequilibrium driving force* or simply as the *(net) drive* (see SI, §1D: *Discussion of various ways of quantifying dissipation* for discussion of different quantitative aspects of dissipation). In addition, we now adopt the transcriptional potencies *r*_*P*_ = *r*_*XP*_ = 1 and *r*_*S*_ = *r*_*X*_ = 0. This choice makes our response observable ⟨*r*⟩ _mRNA_ the probability that polymerase is bound to the genome.

Typical empirical binding energies, diffusion-limited rates, and single-molecule kinetic measurements yield order-of-magnitude estimates for the eight rates governing transcription at equilibrium (see SI, §B:*Order of magnitude estimated rate constants for prokaryotic transcription* and Fig. 1A). First, we choose a set of default rates consistent with these orders-of-magnitude (given in the lower right stem plot of Fig 3C). Next, we investigate how breaking detailed balance by spending energy to increase or decrease a single rate constant at a time—while keeping the seven other rates fixed at biological default values—modulates the transcriptional response curve. Hydrolyzing an ATP molecule makes available ≈20 *k*_*B*_*T* of energy (BNID 101701, (32); (33)) that can be used as a chemical potential gradient to drive transitions (for instance, by powering an enzymatically-assisted pathway (34)). This amount of free energy is also the scale observed to power active processes like biomolecular motors (35). Accordingly, to conservatively emulate a biological energy budget, we allot a maximum of just two ATP hydrolyses’ worth of free energy, | Δ*μ*| ≤40 *k*_*B*_*T*, to break detailed balance. This budget for drive allows a given individual rate to be scaled by up to a factor exp[Δ*μ/k*_*B*_*T*] = exp[*±*40].

**Fig. 3.**
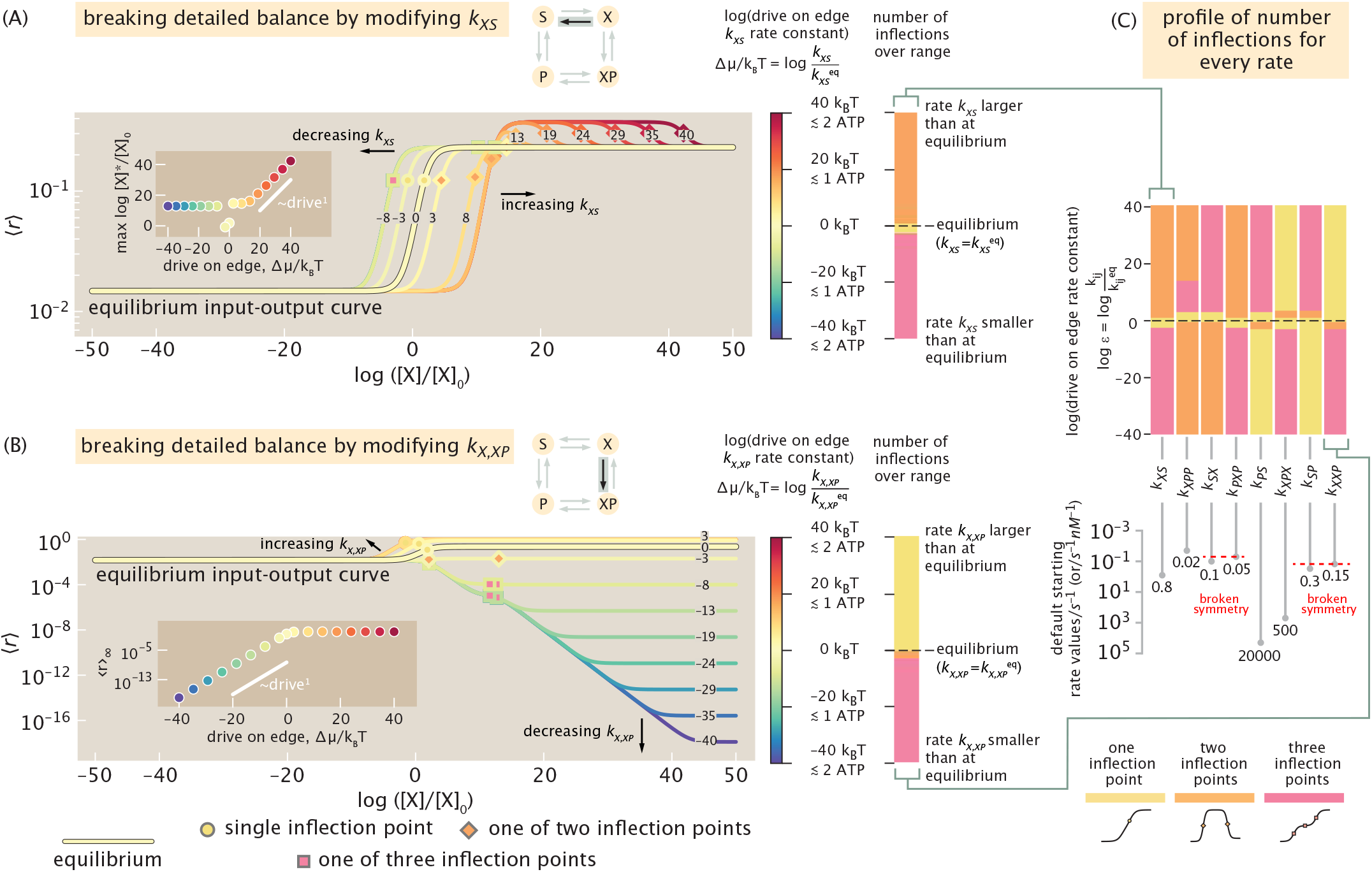
Systematically breaking detailed balance edge-by-edge. (A) Example of how spending energy to modify a single rate (here, *k*_*XS*_)—while the seven other rates remain fixed—changes the response curve away from default equilibrium behavior (pale yellow curve labeled “0” net drive and outlined in black). Responses from rate values larger than (or smaller than) at equilibrium are shown in increasingly red (or blue) colors, respectively; curves are also labeled with the numerical values of the net drive that generated them in *k*_*B*_*T* units (positive for an increase; negative for a decrease). Each curve’s resulting inflection points are marked by yellow, orange, or pink markers, denoting one to three inflection points (respectively), and summarized in the associated one-dimensional (shape phenotypic) phase-diagram with the same colors on the right. Inset: the position of the final inflection point max ln [*X*]^*^*/*[*X*]_0_ versus net drive (power law exponent is ∼ 1); eccentric points near zero drive result from the shifts in shape phenotype in that vicinity. (B) Another representative behavior is displayed when *k*_*X,XP*_ is instead the rate varied. Inset: the saturation *r* _∞_ versus net drive (power law exponent is ∼ 1). (C) Summary of how all eight rates respond to energy expenditure to realize different regulatory shape phenotypes. Below, stem plots give precise values of each default rate constant at equilibrium. (These rates acknowledge initial “broken symmetries” among the rates that violate the conditions Eq. 5 by default, facilitating more ready access to nonmonotonicity. The SI Appendix, §2K, documents the impact of departing from different default starting rates that instead satisfy Eq.5.) (Here, the reference concentration scale setting the horizontal offset of the concentration axis is [*X*]_0_ ≡ 1 nM.)

Applied edge-by-edge, this procedure reveals that biologically-feasible energy expenditures dramatically modify the response curve and easily attain all three regulatory shape phenotypes. Illustrating this regulatory plasticity, Fig. 3A shows how breaking detailed balance by scaling a rate up (increasingly red curves) or down (increasingly green-blue curves) can shift response curves to the left or right on the horizontal log[*X*] axis (effectively tuning what EC50 formerly represented at equilibrium), and also smoothly change the number of inflection points. Yet even for the same net nonequilibrium driving force, the consequences of breaking detailed balance depend significantly on the edge it is broken along. Fig. 3B shows another representative behavior by modifying a different edge, where the major effect of departing equilibrium is to modulate the leakiness, saturation, or intermediate scales of the response. Despite the diversity of this regulation, quantitatively-regular control behavior emerges as well: inset plots emphasize that phenotypic properties such as the position, max {log[*X*]^*^}, of the final inflection point and the saturation, ⟨*r*⟩ _∞_, scale as power laws with the net drive over some regimes.

This broad regulatory flexibility is sustained over all eight rate constants, whose comprehensive response behaviors under drive are analyzed in the SI, §2K: *Systematic census of effects of pushing on one and two edges*. Fig. 3C summarizes how driving each rate attains different shape phenotypes (number of inflections). Notably, any rate can be driven to access any of the three response shape phenotypes at some small, biologically-feasible dissipation. Yet the minimum nonequilibrium driving force values needed to unlock a given phenotype—and the fraction of rate space manifesting said phenotype—varies markedly across the rates. For instance, the two-inflection-point nonequilibrium response shape (orange) is only reached for a fairly narrow, fine-tuned region of drive for the rates *k*_*PS*_, *k*_*XP,X*_, *k*_*SP*_, and *k*_*X,XP*_, but is the most common shape phenotype over finite net drives for the rates *k*_*XS*_, *k*_*XP,P*_, *k*_*SX*_, and *k*_*P,XP*_. Such variable consequences of injecting energy along different rate transitions reflect the privileged roles that states *XP* and *P* play in the graph, given that their probability is the transcriptionally-potent response we monitor. The contrasting impacts of modifying each edge are also sensitive to the default rates that define the system’s biological equilibrium starting point, a revealing dependence that we will return to shortly in the final Results section.

### Breaking detailed balance two edges at a time

Adjusting one edge at a time, as we have just investigated, is but one of many ways a network could invest energy to control its input-output function. Indeed, the classical scheme of kinetic proofreading recognized that many steps could each be driven independently (36), as has later been repeatedly observed in the multistep ways that T-cell or MAPK activation implement kinetic proofreading (37–40) or in mechanochemical operation of myosin motors (41). How do such distributed investments of energy afford expanded control of response functions? To understand this question, we now appraise how breaking detailed balance along up to two edges at a time expands how different response behaviors may be accessed. With two independent drives (one for each edge’s departure from its default biological value), the formerly-one-dimensional phase diagrams of Fig. 3 become slices of two-dimensional phase diagrams that map where response shapes are reached (see Fig. 4A-B; and also the census of how all twenty-eight rate pairs behave found in the SI, §2K).

**Fig. 4.**
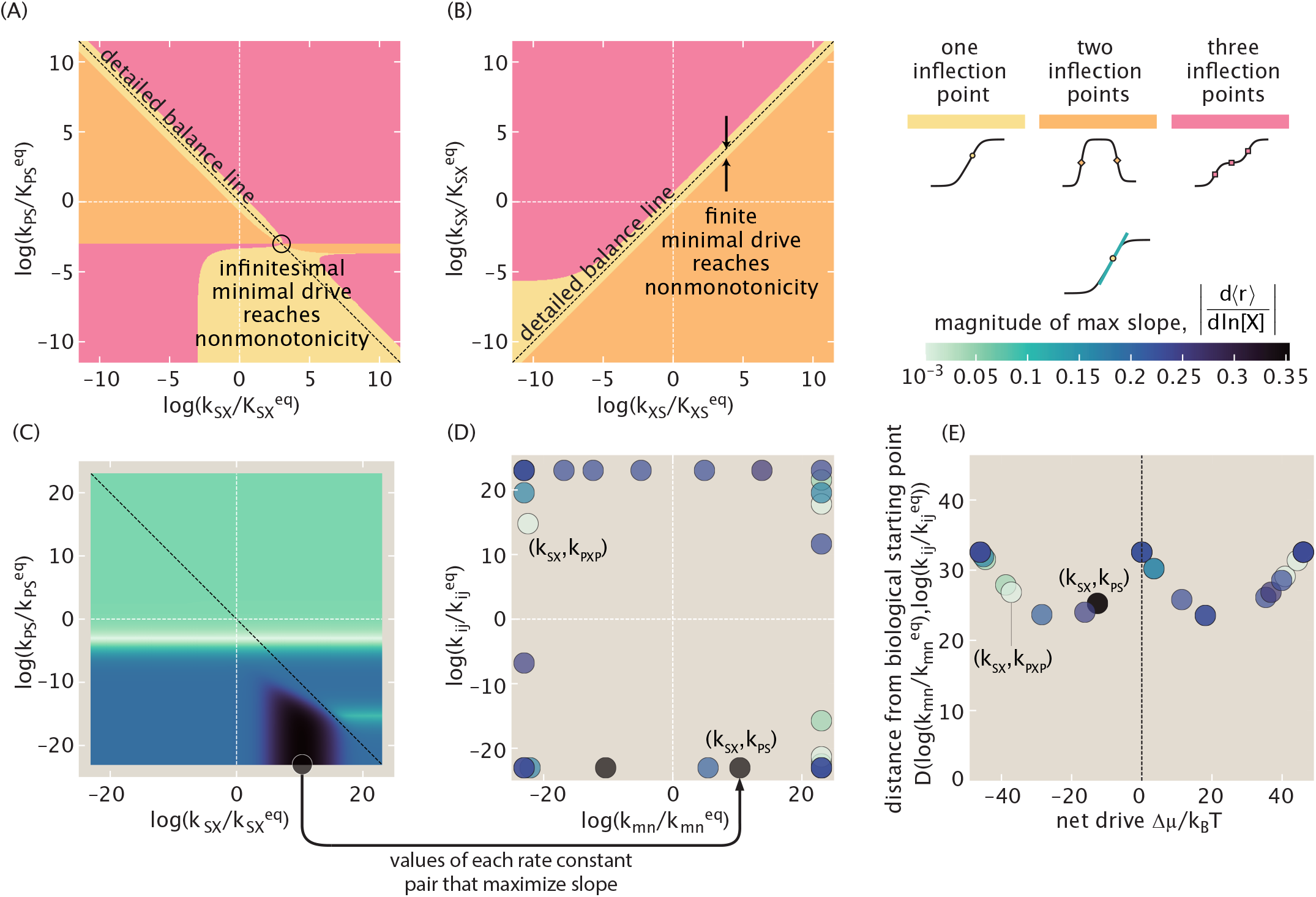
Breaking detailed balance along two edges unlocks higher sensitivity and multiply-inflected outputs with smaller drive than required for breaking detailed balance along single edges. (A) Adjusting the rate pair (*k*_*SX*_, *k*_*P S*_)—while fixing the other six rates at their default biological values at equilibrium (of Figure 1A and Figure 3C’s stem plot)—varies the number of inflection points (light yellow: one inflection, orange: two inflections, pink: three inflections), in a 2D analog of Figure 3. Specifically, this rate pair illustrates a case where nonmonotonic two-inflection curves can be reached with only an infinitesimal net drive. (B) In contrast, when tuning (*k*_*XS*_, *k*_*SX*_), a finite minimum drive is needed to access nonmonotonicity; numerical sampling reveals that this total drive is the same as required while only tuning one edge at a time. (C) Maxima of raw slope *d ⟨r⟩ /d* ln [*X*]*/*[*X*]_0_ over the same modulations (axes) of the rate pair (*k*_*SX*_, *k*_*P S*_) shown in (A), with slope-maximizing rates within the permissible rate space indicated with a circle. [*X*]_0_ ≡ 1 nM is a reference concentration. (D) Overlaying the same positions of maximal slope for all twenty-eight rate pairs emphasizes that optimal slopes are found at the boundary of the permissible rate space. Marker colors reflect the maximal slope achieved for each rate pair. Panel (E) summarizes the behavior of panel (D) by representing each optimal rate pair value with two important natural parameters: the net drive Δ*μ/k*_*B*_*T* (either the log ratio or log product of each rate’s difference from their equilibrium starting values, depending on the relative (counter)clockwise orientation of the rates in a pair); and the net total distance the optimal values are found from their starting values in rate space, 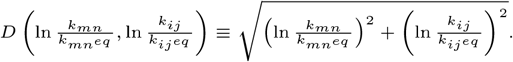.

Geometrically more complex than their one-edge equivalents in Fig. 3, these two-edge phase diagrams expose new ways to transition between the shape phenotypes. One measure of this new facility is the energetic cost needed to reach nonmonotonic (two inflection-point) response curves. Starting from biological equilibrium, what is the minimum net drive Δ*μ*_0_ required for the response to become nonmonotonic, when energy can be injected along just one edge at a time (Fig 3) or up to two edges at a time (Fig. 4A & 4B)? Regarding this question, we find that the 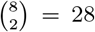 possible pairs of edges can be divided into two types. A few—like the edge pair (*k*_*XS*_, *k*_*SX*_) illustrated in Fig. 4B—require the same finite total dissipation to reach nonmonotonicity as needed if only pushing on either individual edge. However, the majority of rate pairs—such as the edge pair (*k*_*SX*_, *k*_*PS*_)—offer a dissipative bargain: by controlling both rates it is possible to find a point in rate space where only an infinitesimal departure from detailed balance activates nonmonotonicity (as circled in 4A). These inifinitesimal minimal drives contrast the finite drives always required while modifying single edges (Fig. 3C). This new economy is enjoyed by the 22 rate pairs that include at least one of the four special rates *k*_*X,XP*_, *k*_*SP*_, *k*_*XP,X*_, or ; their membership will be a clue for identifying critical conditions on nonmotonicity we deduce in the next (and final) Results section.

The richer behaviors achievable by breaking detailed balance along two rates (instead of just one) become even more pronounced from the lens of sensitivity. The heatmap of Fig. 4C depicts the maximal unnormalized sharpness *d* ⟨*r* ⟩ /*d* ln[*X*] reached by modifying the rate pair (*k*_*SX*_, *k*_*PS*_) (the same rates mapped phenotypically in the phase space of Fig. 4A). If only one rate constant at a time were allowed to be driven, only the slices of sharpness along the white dotted *x* = 0 and *y* = 0 vertical and horizontal lines would be accessible, at most realizing a maximal unnormalized sharpness of ;S 0.15 with respect to the concentration [*X*] on a log scale. However, once both edges can be modified, it becomes possible to access the maximal slope region on the lower right, yielding a greater maximum sensitivity of about 0.35. Repeating this procedure for all 28 rate pairs, as shown in Fig. 4D, we find that the points in rate space that maximize slope all require *both* rate constants in each pair to be modified from their default equilibrium values (lying away from the *x* = 0 and *y* = 0 vertical and horizontal lines). To maximize sensitivity, all rate pairs show one (but usually not both) rate constant that has been driven to the maximal extent allowed by the nonequilibrium driving force budget (localizing optimal points to the borders—but not necessarily corners—in Fig. 4D). The net drive Δ*μ* ensuing from both rate’s departure from their equilibrium values is often distinct from those independent departures. Fig. 4E recasts the same slope-maximizing points in Fig. 4D in terms of these two separate properties (the net drive Δ*μ*, and the average geometric distance, *D*, each edge moved from its biological starting point.) Different rate pairs show dramatically different optimal maximum sensitivities at varying cost: choosing to break detailed balance along the (*k*_*SX*_, *k*_*PS*_) can achieve a maximal slope of about 0.35 (probability units per *e*-fold change in [X]) at a net drive of only Δ*μ*≈ 10 *k*_*B*_*T* (dark grey marker), but choosing less wisely the rate pair (*k*_*SX*_, *k*_*PXP*_) at best attains a slope of about 0.054 (probability units per *e*-fold change in [X]), even while spending a net energy Δ*μ* 35 *k*_*B*_*T* almost four times as large. Collectively, these findings highlight how prudently distributing dissipation over the transitions in a network can achieve more precise and dramatic responses.

### Generic rate conditions forbid access to nonmonotonic responses

Why, as we have seen, are nonmonotonic responses accessed with different ease while driving some rates—or still more economically, rate pairs—rather than others? How do the default equilibrium rates from which biology departs affect the tunability of responses? Confronting these questions leads us to glean general kinetic conditions that enable or forbid nonmonotonicity. We reformulate the criterion for nonmonotonicity to explicitly invoke net drive and rate constants (see SI, §2L:*Crucial imbalances in rate-constants are required for nonmonotonic responses*). Using these analytical arguments, we determine that nonmonotonicity is forbidden for any net drive when transition rates satisfy the following, surprisingly loose, conditions:

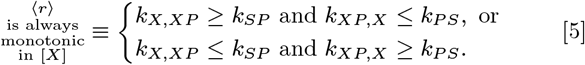

That is, if the presence of the transcription factor on the genome increases or decreases the polymerase’s binding rate in a sense opposite to its effect on the unbinding rate (or leaves either unchanged), the response must depend on the transcription factor monotonically. Only when the transcription factor plays a functionally “ambiguous,” dualistic role—coherently changing both the polymerase’s binding and unbinding rates (that themselves have opposite effects on the response)—may the response become nonmonotonic under a sufficient net drive. Since access to nonmonotonicity is governed by kinetic conditions in Eq. (5)—but thermodynamic parameters instead set whether a response is globally activating or repressing (SI §)—the qualitative origin of nonmonotonicity stems from when kinetic and thermodynamic aspects in the system oppose each other.

This condition of Eq. 5 helps explain why some rates and rate pairs reach regulatory shape phenotypes so differently under drive, and how default starting rate constants matter. A comprehensive census of responses while driving one edge at a time when default rates *satisfy* Eq. 5 is provided in the SI Appendix.

Instructively, Eq. 5 demands that when the transcription factor does *not* change the polymerase’s (un)binding rates— namely, either *k*_*X,XP*_ = *k*_*SP*_ or *k*_*XP,X*_ = *k*_*PS*_ —the response must be monotonic. By default, under the often reasonable classical assumption that the binding rate of polymerase is purely diffusion-limited (1), the transcription factor indeed may not affect the polymerase’s binding rate, thus forcing the response to be monotonic.^§^ This type of biophysical constraint may contribute to why monotonic transcriptional responses are most canonically pictured as monotonic. However, while plausible, this biophysical scenario is hardly inescapable or universal. In fact, even for architectures as “simple” as *lac* repression, there is gathering empirical evidence that proteins associate with DNA binding sites under more intricate regulation than merely diffusion (42). Transcription factors that mediate steric access to the genome (dissipatively or not), such as via DNA looping (43), may also be especially prone to contravene this condition.

## Discussion

In this work, we dissected how spending energy transforms the control of gene expression in a minimal and common transcriptional motif. Harnessing a kinetic description and diagrammatic procedure from graph theory, we found that any transcriptional outputs follow a universal form with respect to a control parameter like a transcription factor’s concentration. We discovered these responses may only adopt three shapes, including an equilibrium-like (monotonic, sigmoidal) response. Uniquely out of equilibrium, however, two unexpected and noncanonical output behaviors become possible: a doubly-inflected, nonmonotonic response; and a triply-inflected, monotonic response. Underneath wide parametric complexity, we established tight global bounds on transcriptional response’s maximal sensitivity and learned these can vary and tradeoff with response shape. Next, we systematically mapped how biologically-feasible amounts of energy along single rates or rate pairs control responses. These findings established that the noncanonical responses are easily accessed around rates plausible for transcription, especially when dissipation can be distributed more widely over a network. Last, we uncovered global and transparent kinetic conditions that forbid (or enable) novel nonmonotonic responses.

The flexible regulation unlocked by nonequilibrium could be widely biological salient. Responses that can show three inflection points—instead of just one at equilibrium—could effectively accomplish the role of two classical (singly-inflected) input-output functions. Since an inflection can mark a local region of enhanced output sensitivity, and effectively implement a threshold, this functionality could allow cells to achieve distinct cellular fates, such as in Wolpert’s classical French Flag model (44). By contrast to our small architecture, canonical pictures of multiple thresholded responses usually require multiple genes—often at least one specific gene per threshold (45). One imporant example is the celebrated Dorsal protein in *Drosophila*, where two critical thresholds have been proposed to accomplish *twist* gene activation and *decapentaplegic* gene repression to help establish distinct parts of dorsal patterns in embryonic development (46, Fig. 2.26, p. 64). We propose that triply-inflected responses from a single gene could accomplish some of this same functionality with a smaller architecture.

Nonmonotonic response functions with two inflection points could empower cells to accomplish more sophisticated signal processing, such as band-pass or band-gap filtering of chemical inputs, and/or generate temporal pulses of chemical outputs. Similar implications have been been explored by Alon & coworkers, *inter alios*, who established how nonmonotonic outputs can be produced by chaining together incoherent feed-forward loops (47–50). To achieve more complex outputs, these networks use transcriptional interactions among multiple genes at equilibrium—e.g. from two to six (or more) genes in such examples. Hence these networks operate with comparatively larger sizes and timescales than mere binding-unbinding reactions on a single gene’s regulatory network like the square graph we study in this report. We suggest these comparisons contribute new material to a maturing discourse about when and how biology uses thermodynamic or kinetic control mechanisms (34, 41).

Even responses that remain “equilibrium-like” with a single inflection benefit from energy expenditure, since our bounds establish they may be up to two times more sensitive than at equilibrium, and enjoy new kinetic (instead of merely thermodynamic) ways of controlling the location of the governing inflection point (EC50).

While only mild net drives transpire to unlock useful regulatory shapes and traits, our analysis emphasizes other mechanistic factors that govern how easily these behaviors can be accessed, or measured as signatures of nonequilibrium in natural or synthetic settings.

First, the biological network’s architecture determines whether these new macroscopic behaviors can be attained at all. Although prokaryotic gene regulation has regularly shown a compelling coherence between quantitative measurements and equilibrium statistical mechanical models (including demanding studies from our own laboratories over the past two decades (6, 19, 24, 51, 52) and beyond (43)), many of the most fiercely interrogated systems (e.g. the *lac* repressor) are indeed exactly those with acyclic network topologies that make nonequilibrium steady-states impossible (without open fluxes) and guarantee detailed balance. This reflects a possible overrepresentation of biological settings where detailed balance may be expected *a priori* to apply on mere structural grounds. On the other hand, the means to spend energy biochemically clearly exist, even in bacteria through two-component regulatory systems (53) and other active settings like nucleosome remodeling in eukaryotes (5). Hence our findings invite a renewed and vigorous reappraisal of whether signatures of nonequilibrium are in fact lurking in architectures that are more prone to accommodate it, such as the four-state “simple activation” motif we discussed here. Moreover, the measurements (or synthetic biological perturbations) needed to map the nonequilibrium landscape of transcriptional responses must differ from the convenient binding site modifications (e.g. parallel promoter libraries (19, 54)) previously used to test equilibrium models, since manipulating binding energies inherently preserves detailed balance. Developing fresh experimental approaches to augment or attenuate a single transition between microstates (or set of transitions) *in situ* to break detailed balance is a crucial direction of future empirical work, whose value is advocated for by our results. To manipulate and probe tractable models of transcription, these methods might include optogenetic control (55, 56), or suitable adjustments of governing enzyme concentrations or activities.

Second, where energy is invested crucially dictates which regulatory behaviors are available. We found that investing energy along more than one rate at once was capable of achieving more dramatic response curves more economically. This finding may help explain the many observations in biological systems where energy is independently injected along multiple steps (36–41). However, since each independently-regulated injection of energy may also be accompanied by architectural costs, not all examples of biological regulation may contain the distributed dissipation machinery required to make novel nonequilibrium response signatures conspicuous.

Third, the structures of responses while breaking detailed balance edge-by-edge, and our general kinetic criteria that forbid nonmonotonicity, highlight that certain critical imbalances between rate constants are needed to produce the most conspicuously non-sigmoidal shape phenotypes available out of equilibrium. On basic biophysical grounds, some natural systems may—or may not—exhibit the required rate imbalances to make novel responses as easy to activate (see SI, §L.2: *Conditions that suffice to forbid nonmonotonicity*).

Indeed, the rate imbalances required to produce nonmonotonicity we found are non-obvious. These kinetic criteria have significant implications for organizing parameter explorations. For instance, we show in the SI, §2M: *Implications of critical symmetry conditions for widespread numerical screens* that an exciting study just published (13) exploring the informational consequences of nonequilibrium in a four-state model (that is mappable to our setting) imposes simplifying assumptions on rate constants that in fact preclude the possibility of non-monotonic responses, according to our monotonicity criterion. We expect that our approach and kinetic criteria will help future works include and capture the regulatory consequences of these rich behaviors. We anticipate this flexibility may be especially germane for environments that present nonuniform input statistics.

The contrast between the nonequilibrium steady-states possible to support using this “simple activation” architecture, and the difficulty of sustaining nonequilibrium steady-states in a simple repression architecture that lacks a cycle, also possibly provides a new design principle to understand the timeless question of why both activators and repressors are employed as distinct architectures when they can produce the same mean gene expression. Intriguing rationalizations based on ecological demand have been offered for why these architectures are used differently in *E. coli*, such as the classical proposal by Savageau (57–59). We speculate that another, quite distinct, feature—the very possibility of using nonequilibrium to steer input-output response curves so flexibly—may also contribute to why organisms might use a simple-activation (or other cycle-containing) architecture over acyclic architectures, all other features being equal. Whether this nonequilibrium controllability significantly shapes the natural incidence of regulatory architectures can only be assessed using quantitative measurements of input-output behaviors from a much broader set of architectures than the relatively narrow (e.g. Lac repressor, Bicoid, CI in bacteriophage-*λ* switch) subjects of existing analyses.

Our work provides explicit maps of parameter spaces that can guide the naturalist looking for whether this expanded regulation occurs naturally in some manifestations of transcription. This information is also a guide to the synthetic biologist who endeavors to engineer such responses in genetic circuits and exploit the advantages of producing complex regulation using a small driven network, instead of a comparatively larger, more slowly tuned network of multiple genes at equilibrium.

Beyond advocating for experimental progress, our findings invite many theoretical extensions. How dissipation affects the intricate tradeoffs between sensitivity, specificity, speed, and stochasticity in (steady-state or transient) gene regulation is a large, open, physiologically-relevant question amenable to further graph-theoretic dissection. In addition, we hope for deeper analytical rationalization of our bounds on sensitivity; our upper bounds surely share similar foundations with looser, more architecturally general, bounds recently and insightfully established by Owen & Horowitz (29), though our additional lower bounds and different mathematical quantities suggest separate theoretical ingredients.

Overall, we foresee that graph-theoretic treatments like we have deployed here—and as have been first so powerfully established and refined by other foundational investigators (16)—will produce further dividends when addressing still more sophisticated networks. Logically (but not psychologically) equivalent to tedious, purely algebraic analysis of steady-state probabilities, these perspectives promise to be engines of discovery amid the complexity of nonequilibrium biology, just as diagrammatic analyses such as Feynman diagrams continue to catalyze progress in field theory and particle physics (60, 61).

## Materials and Methods

### Nonequilibrium steady-state probabilities via the Matrix Tree Theorem

Consider a continuous-time Markov chain with *N* states, whose transition rates *k*_*ij*_ between states *i* and *j* are stored in the *j, i*th element of the transition matrix **L**, and so the probabilities **p**(*t*) = [*p*_1_, …, *p*_*N*_]^*T*^ of finding the system in these states evolve according to

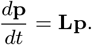

(With this convention of **p** as a column vector, the columns of the matrix **L** sum to zero and the diagonal entries are accordingly 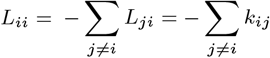 Note that (**Lp**)_*i*_ is the net probability flux entering the node *i*. Identifying our Markov system as a weighted graph, a *spanning tree* over the states is a set of *N* − 1 edges that visits every state exactly once. A spanning tree 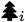 *rooted* in a state *i* contains no outgoing edges from state *i* (and exactly one outgoing edge for every other state *j* ≠ *i*). (These notions are summarized in the example of Fig. 1B.) The **Matrix Tree Theorem** (MTT) (also known as the Markov Chain Tree Theorem) states that at steady state 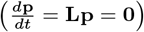, the statistical weight of the *i*th state is the sum of products of rate constants over spanning trees rooted in node *i*

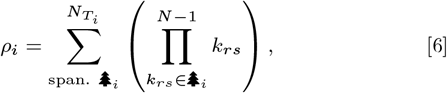

where 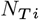 is the number of spanning trees rooted in *i* (16, 21). This weight *ρ*_*i*_ is the relative odds of finding the system in state *i* as a fraction of all the statistical weights 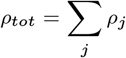, namely *ρ*_*i*_ */ρ*_*tot*_. Applying the MTT to the regulatory motif of Fig. 1A indicates that any steady-state probabilistic observable depends on the transcription factor control parameter [*X*] according to Eq. 1 (see SI).

### Emergent shape parameters & shape phenotypes

The collapsed shape representation of Eq. 3 allows us to solve for the number of positive solutions to *d*⟨*r*⟩ /*d* ln ([*X*]*/*[*X*]_0_), yields the numbers of possible inflection points (via, for instance, Descartes’ rule of signs or explicit inequality solving) and hence shapes (see SI). Numerical and symbolic analysis of the space formed by these two emergent shape parameters (*a, b*) (Eq. 3 and SI appendix) helps establish our global bounds on sensitivity. Ultimately, this collapsed representation is also a crucial theoretical stepladder to find the generic conditions forbidding nonmonotonicity given in Eqs. 5 (see SI).

### Single edge and edge pair perturbations

We estimated default biological rates for transcription at equilibrium by synthesizing reported binding affinities, association rates, and diffusion constants. We solved the condition for an inflection point symbolically and numerically (see SI).

## Data & Availability

All symbolic and numerical code used for this study’s analyses and presented figures will be available open-source. See https://github.com/glsalmon1/graphnoneq.

## ACKNOWLEDGMENTS

We gratefully acknowledge feedback from and formative discussions with Forte Shinko; Nicholas Lammers; Avi Flamholz; Jordan Horowitz; Jeremy Owen; and Jané Kondev. G.L.S. thanks the US NSF GRFP under Grant DGE-1745301 and Caltech’s Center for Environmental Microbial Interactions for support. R.P. is supported by the Rosen Center at Caltech and the NIH 1R35 GM118043 (MIRA). H.G.G. is supported by NIH R01 Award (R01GM139913), and the Koret-UC Berkeley-Tel Aviv University Initiative in Computational Biology and Bioinformatics. H.G.G. is also a Chan Zuckerberg Biohub–San Francisco Investigator.

## Footnotes

* The two-parameter simplicity of Eq. 3 is one possible nonequilibrium sophistication of the (usually one-parameter) data collapses used to unify simpler, equilibrium, two-state physiological responses (27) and regulation (6) in bacteria.

† Throughout our analysis and discussion in this paper, we monitor the shape, number of inflection points, and sensitivity of transcriptional outputs with respect to the control parameter of the concentration of transcription factor, on a *logarithmic* scale. We use this logarithmic convention in alignment with common practice in biochemical and transcriptional studies (6, 28, 29).

‡ We use the phrase “regulatory (shape) phenotype,” referring to the overall shape of a response curve, to distinguish our meaning from the usage of Reference (2), who instead referred to specific *quantitative traits* within curves of a single mathematical shape (such as sensitivity or noise) as “regulatory phenotypes.”

§ By contrast, by the assumption that the transcription factor has the typical biophysical effect of changing the affinity between the polymerase and genome, the polymerase’s off-rate from the genome *is* affected by the transcripton factor’s presence, and *k*_*XP,X*_ *I*= *k*_*P S*_. So usually it is not an equality between polymerase’s off-rates that prevents a response from being nonmonotonic.

## Notes

### Competing Interest Statement

The authors have declared no competing interest.

## References

1. M Morrison, M Razo-Mejia, R Phillips, Reconciling kinetic and thermodynamic models of bacterial transcription. PLoS computational biology 17, e1008572 (2021).

2. R Grah, B Zoller, G Tkačik, Nonequilibrium models of optimal enhancer function. Proc. Natl. Acad. Sci. 117, 31614–31622 (2020).

3. F Wong, J Gunawardena, Gene regulation in and out of equilibrium. Annu. review biophysics 49, 199–226 (2020).

4. A Coulon, CC Chow, RH Singer, DR Larson, Eukaryotic transcriptional dynamics: from single molecules to cell populations. Nat. reviews genetics 14, 572–584 (2013).

5. R Shelansky, H Boeger, Nucleosomal proofreading of activator–promoter interactions. Proc. Natl. Acad. Sci. 117, 2456–2461 (2020).

6. M Razo-Mejia, et al., Tuning transcriptional regulation through signaling: a predictive theory of allosteric induction. Cell Syst. 6, 456–469 (2018).

7. IS Farmer, CW Jones, The energetics of escherichia coli during aerobic growth in continuous culture. Eur. journal biochemistry 67, 115–122 (1976).

8. B Zoller, T Gregor, G Tkačik, Eukaryotic gene regulation at equilibrium, or non? arXiv preprint 2110.06214 (2021).

9. J Rodenfels, KM Neugebauer, J Howard, Heat oscillations driven by the embryonic cell cycle reveal the energetic costs of signaling. Dev. cell 48, 646–658 (2019).

10. Q Yu, D Zhang, Y Tu, Inverse power law scaling of energy dissipation rate in nonequilibrium reaction networks. Phys. review letters 126, 080601 (2021).

11. X Yang, et al., Physical bioenergetics: Energy fluxes, budgets, and constraints in cells. Proc. Natl. Acad. Sci. 118, e2026786118 (2021).

12. H Ge, M Qian, H Qian, Stochastic theory of nonequilibrium steady states. part ii: Applications in chemical biophysics. Phys. Reports 510, 87–118 (2012).

13. NC Lammers, AI Flamholz, HG Garcia, Competing constraints shape the nonequilibrium limits of cellular decision-making. Proc. Natl. Acad. Sci. 120, e2211203120 (2023).

14. KM Nam, R Martinez-Corral, J Gunawardena, The linear framework: using graph theory to reveal the algebra and thermodynamics of biomolecular systems. Interface Focus. 12, 20220013 (2022).

15. J Gunawardena, A linear framework for time-scale separation in nonlinear biochemical systems. PloS one 7, e36321 (2012).

16. I Mirzaev, J Gunawardena, Laplacian dynamics on general graphs. Bull. mathematical biology 75, 2118–2149 (2013).

17. TL Hill, Free energy transduction and biochemical cycle kinetics. (Courier Corporation), (2013).

18. H Qian, Open-system nonequilibrium steady state: statistical thermodynamics, fluctuations, and chemical oscillations (2006).

19. WT Ireland, et al., Deciphering the regulatory genome of escherichia coli, one hundred promoters at a time. Elife 9, e55308 (2020).

20. M Rydenfelt, HG Garcia, RS Cox III, R Phillips, The influence of promoter architectures and regulatory motifs on gene expression in escherichia coli. PLoS one 9, e114347 (2014).

21. R Phillips, HG Garcia, Physical Genomics: From E. coli to Elephants. (Princeton University Press), (2023).

22. R Shelansky, et al., A telltale sign of irreversibility in transcriptional regulation. bioRxiv pp. 2022–06 (2022).

23. LA Mirny, Nucleosome-mediated cooperativity between transcription factors. Proc. Natl. Acad. Sci. 107, 22534–22539 (2010).

24. HG Garcia, R Phillips, Quantitative dissection of the simple repression input–output function. Proc. Natl. Acad. Sci. 108, 12173–12178 (2011).

25. HG Garcia, J Kondev, N Orme, JA Theriot, R Phillips, Thermodynamics of biological processes in Methods in enzymology. (Elsevier) Vol. 492, pp. 27–59 (2011).

26. L Bintu, et al., Transcriptional regulation by the numbers: models. Curr. opinion genetics & development 15, 116–124 (2005).

27. LR Swem, DL Swem, NS Wingreen, BL Bassler, Deducing receptor signaling parameters from in vivo analysis: Luxn/ai-1 quorum sensing in vibrio harveyi. Cell 134, 461–473 (2008).

28. AJ Meyer, TH Segall-Shapiro, E Glassey, J Zhang, CA Voigt, Escherichia coli “marionette” strains with 12 highly optimized small-molecule sensors. Nat. chemical biology 15, 196–204 (2019).

29. JA Owen, JM Horowitz, Size limits the sensitivity of kinetic schemes. Nat. Commun. 14, 1280 (2023).

30. JA Brophy, CA Voigt, Principles of genetic circuit design. Nat. methods 11, 508–520 (2014).

31. H Qian, Phosphorylation energy hypothesis: open chemical systems and their biological functions. Annu. Rev. Phys. Chem. 58, 113–142 (2007).

32. R Milo, P Jorgensen, U Moran, G Weber, M Springer, Bionumbers—the database of key numbers in molecular and cell biology. Nucleic acids research 38, D750–D753 (2010).

33. QH Tran, G Unden, Changes in the proton potential and the cellular energetics of escherichia coli during growth by aerobic and anaerobic respiration or by fermentation. Eur. journal biochemistry 251, 538–543 (1998).

34. JM Horowitz, K Zhou, JL England, Minimum energetic cost to maintain a target nonequilibrium state. Phys. Rev. E 95, 042102 (2017).

35. MM Lin, A general efficiency relation for molecular machines. arXiv preprint 2210.04380 (2022).

36. JJ Hopfield, Kinetic proofreading: a new mechanism for reducing errors in biosynthetic processes requiring high specificity. Proc. Natl. Acad. Sci. 71, 4135–4139 (1974).

37. DM Britain, JP Town, OD Weiner, Progressive enhancement of kinetic proofreading in t cell antigen discrimination from receptor activation to dag generation. Elife 11, e75263 (2022).

38. PS Swain, ED Siggia, The role of proofreading in signal transduction specificity. Biophys. journal 82, 2928–2933 (2002).

39. TW McKeithan, Kinetic proofreading in t-cell receptor signal transduction. Proc. national academy sciences 92, 5042–5046 (1995).

40. W Cui, P Mehta, Identifying feasible operating regimes for early t-cell recognition: The speed, energy, accuracy trade-off in kinetic proofreading and adaptive sorting. PloS one 13, e0202331 (2018).

41. JD Mallory, AB Kolomeisky, OA Igoshin, Kinetic control of stationary flux ratios for a wide range of biochemical processes. Proc. Natl. Acad. Sci. 117, 8884–8889 (2020).

42. E Marklund, et al., Sequence specificity in dna binding is mainly governed by association. Science 375, 442–445 (2022).

43. T Kuhlman, Z Zhang, MH Saier Jr, T Hwa, Combinatorial transcriptional control of the lactose operon of escherichia coli. Proc. Natl. Acad. Sci. 104, 6043–6048 (2007).

44. L Wolpert, Positional information and the spatial pattern of cellular differentiation. J. theoretical biology 25, 1–47 (1969).

45. S Papageorgiou, Y Almirantis, Gradient model describes the spatial-temporal expression pattern of hoxa genes in the developing vertebrate limb. Dev. dynamics 207, 461–469 (1996).

46. L Wolpert, C Tickle, AM Arias, Principles of development. (Oxford University Press, USA), 6 edition, (2015).

47. S Kaplan, A Bren, E Dekel, U Alon, The incoherent feed-forward loop can generate nonmonotonic input functions for genes. Mol. systems biology 4, 203 (2008).

48. S Ishihara, K Fujimoto, T Shibata, Cross talking of network motifs in gene regulation that generates temporal pulses and spatial stripes. Genes to cells 10, 1025–1038 (2005).

49. S Basu, Y Gerchman, CH Collins, FH Arnold, R Weiss, A synthetic multicellular system for programmed pattern formation. Nature 434, 1130–1134 (2005).

50. R Entus, B Aufderheide, HM Sauro, Design and implementation of three incoherent feedforward motif based biological concentration sensors. Syst. synthetic biology 1, 119–128 (2007).

51. DL Jones, RC Brewster, R Phillips, Promoter architecture dictates cell-to-cell variability in gene expression. Science 346, 1533–1536 (2014).

52. RC Brewster, et al., The transcription factor titration effect dictates level of gene expression. Cell 156, 1312–1323 (2014).

53. AY Mitrophanov, EA Groisman, Signal integration in bacterial two-component regulatory systems. Genes & development 22, 2601–2611 (2008).

54. TC Yu, et al., Multiplexed characterization of rationally designed promoter architectures deconstructs combinatorial logic for iptg-inducible systems. Nat. communications 12, 325 (2021).

55. W Zhang, et al., Optogenetic control with a photocleavable protein, phocl. Nat. Methods 14, 391–394 (2017).

56. H Liu, G Gomez, S Lin, S Lin, C Lin, Optogenetic control of transcription in zebrafish. PloS one 7, e50738 (2012). Mahdavi & Salmon et al.

57. MA Savageau, Design of molecular control mechanisms and the demand for gene expression. Proc. Natl. Acad. Sci. 74, 5647–5651 (1977).

58. MA Savageau, Genetic regulatory mechanisms and the ecological niche of escherichia coli. Proc. Natl. Acad. Sci. 71, 2453–2455 (1974).

59. U Alon, An introduction to systems biology: design principles of biological circuits. (CRC press), 2nd edition, (2019) See especially section 7.6”demand rules for gene regulation can minimize errors,” page 129.

60. D Kaiser, Drawing theories apart in Drawing Theories Apart. (University of Chicago Press), (2009).

61. M Veltman, Diagrammatica: the path to Feynman diagrams. (Cambridge University Press) No. 4, (1994).

