## Supplementary Appendix for "Flexibility and sensitivity in gene regulation out of equilibrium"

#### This PDF file includes:

Figs. S1 to S15
Table S1
SI References

### 1. Linear Markovian dynamics, $\frac{d\mathbf{p}}{dt} = \mathbf{L}\mathbf{p}$, and cycles, are common

**A. Mathematically, to first order, many dynamics are continuous time Markov chains.** Also referred to as *kinetic schemes* (1) or viewed as representations of *chemical master equations* (2), continuous time Markov chains capture (approximately or exactly) how many systems change in time. When a single (possibly effective) typical timescale $\tau_{ij}$ (or rate $k_{ij} = 1/\tau_{ij}$) is used to describe a transition between every pair of states $i$ and $j$, the description amounts to a continuous time Markov chain. Or, if the $i$th component $p_i(t)$ of the system's state probability evolves in time according to some function $f(\mathbf{p}(t))$ that depends on only the current state, we propose that a Taylor expansion to first order in $\mathbf{p}$ around a (hypothetical) empty system's state $\mathbf{0}$ also yields such a description,

$$\frac{dp_i}{dt} = f_i(\mathbf{p}(t)) \qquad [1]$$

$$= \nabla \mathbf{f}_i^\top (\mathbf{p}(t) - \mathbf{0}) \; + \; (\mathbf{p}(t) - \mathbf{0})^\top \left(\frac{\partial^2 f_i}{\partial \mathbf{p}^2}\right) (\mathbf{p}(t) - \mathbf{0}) \; + \; \cdots \qquad [2]$$

$$\approx \nabla \mathbf{f}_i^\top \mathbf{p}(t) = \sum_j \frac{\partial f_i}{\partial p_j} p_j(t) = \sum_j \frac{\partial \frac{dp_i}{dt}}{\partial p_j} p_j(t); \qquad [3]$$

we can store these equations in a matrix form, defining $L_{ij} \equiv \frac{\partial \frac{dp_i}{dt}}{\partial p_j}$ to give

$$\frac{d\mathbf{p}}{dt} = \mathbf{L}\mathbf{p}. \tag{4}$$

Armed with the fact that total probability is conserved, $\sum_i p_i = 1$, one can further immediately conclude that

$$\frac{d}{dt}\left(\sum_i p_i\right) = \sum_i \frac{dp_i}{dt} = 0 \tag{5}$$

$$= \vec{1}^\top (\mathbf{L}\mathbf{p}) = 0, \tag{6}$$

and since this must hold for arbitrary $\mathbf{p}$, we see that $\vec{1}^\top \mathbf{L} = \vec{0}^\top$, namely the rows of $\mathbf{L}$ sum to zero.[*] So the diagonal entries of $\mathbf{L}$ can be expressed as $L_{ii} = -\sum_{j \neq i} L_{ji}$.

**B. Closed steady-state systems are either equilibrium or cyclic.** Why can we conclude that a graph without cycles cannot show nonequilibrium steady-states (and so must be in detailed balance at steady-state)? Since this question is about graph structures and generic steady-states, we turn to the Matrix Tree Theorem, discussed more fully in the main text and illustrated in this supplement's §D, which emphasizes insights come from the nature of spanning trees.

First, recall that detailed balance occurs when for any pair of states $(i, j)$, the steady-state probabilities satisfy,

$$p_i k_{ij} = p_j k_{ji} \tag{7}$$

$$\rightarrow \rho_i k_{ij} = \rho_j k_{ji}, \tag{8}$$

where we have divided by the common normalizing factor $\rho_{\text{tot}}$ in the second expression such that $\rho_i = p_i/\rho_{\text{tot}}$.

Next, consider how spanning trees in a graph are structured and their algebraic consequences. For any steady state, whether in or out of equilibrium, the statistical weight of a state $i$ is the sum of spanning trees rooted in $i$,

$$\rho_i = \sum_{\text{span. trees } m} \underbrace{\prod_{k_{rs} \in \text{ tree } m} k_{rs}}, \tag{9}$$

$$= \sum_{\text{span. trees } m} T_i^{(m)}, \tag{10}$$

where we have included the algebraic reminder that some $m$th spanning tree $T_i^{(m)}$ rooted in node $i$ is a product of suitable rate constants $k_{rs}$ such that every node is visited exactly once and there is no outgoing edge from the root $i$.

How are the spanning trees rooted in a node $i$ related to those rooted in a connected node $j$? By structural requirement these trees are quite similar. Indeed, *if* a tree $T_i$ rooted in $i$ contains the edge $k_{ji}$, then we can always convert it to a valid spanning tree rooted in $j$ instead by "flipping" that edge to contain $k_{ij}$ instead, building the newly rooted tree $T_j = \frac{k_{ij}}{k_{ji}} T_i$. (This re-rooting works because the rest of the edges in the original $i$-rooted tree $T_i$ have not been altered so still have out-degree exactly one; all the nodes in the graph are still visited by the tree; and now $j$ has out-degree zero, as required of a valid spanning tree rooted in $j$.) If all the spanning trees rooted in $i$ contain the edge $k_{ji}$, then this re-rooting operation works to build all the trees rooted in $j$, giving

$$\rho_j = \sum_m T_j^{(m)} = \sum_m \frac{k_{ij}}{k_{ji}} T_i^{(m)} \tag{11}$$

$$= \frac{k_{ij}}{k_{ji}} \sum_m T_i^{(m)} \tag{12}$$

$$= \frac{k_{ij}}{k_{ji}} \rho_i, \tag{13}$$

which is exactly the requirement of detailed balance between $i$ and $j$.

However, while every spanning tree of an acyclic graph (where $i$ and $j$ are connected) *will* contain the edge $k_{ij}$ or $k_{ji}$ (since there is one path in the graph allowing them to be connected), this is no longer true for graphs containing a cycle: other paths can connect $i$ and $j$ that do not directly contain the $(i, j)$ edges and thus build valid spanning trees. In that case, we cannot always write $T_j = \frac{k_{ij}}{k_{ji}} T_i$ and so cannot factor out $\frac{k_{ij}}{k_{ji}}$ from the weights $\rho_i$ and $\rho_j$. This means that only such cyclic graphs can violate detailed balance at steady-state.

[*] In more (indicial) words, $\sum_i \frac{dp_i}{dt} = \sum_i \left(\sum_j L_{ji} p_j\right) = \sum_j p_j \left(\sum_i L_{ji}\right) = \sum_j p_j \left(L_{jj} + \sum_{i \neq j} L_{ji}\right)$. Since this must hold true for any value of $p_j$, we see that $L_{jj} + \sum_{i \neq j} L_{ji} = 0$ for all states $j$, confirming the form of the diagonal entries of the matrix.

***B.1. Example of an acyclic system: the simple repression motif.*** This connection between structure and the impossibility of violating detailed balance is illustrated in the simple repression motif. Here, repressors are assumed to sterically exclude the polymerase's binding (3, 4); this condition permits just three states in a linear graph that lacks a cycle. Specifically, call "$S$" the empty genome substrate state; $R$ the repressor-bound genome state; and $P$ the polymerase-bound genome state. These states form the linear graph,

$$R \underset{k_{SR}[R]}{\overset{k_{RS}}{\rightleftharpoons}} S \underset{k_{PS}}{\overset{k_{SP}[P]}{\rightleftharpoons}} P. \quad [14]$$

Since there is only one rooted spanning tree per root state, the Matrix Tree Theorem says that the steady-state statistical weights of the states are

$$\begin{pmatrix} \rho_R \\ \rho_S \\ \rho_P \end{pmatrix} = \begin{pmatrix} k_{SR}[R]k_{PS} \\ k_{RS}k_{PS} \\ k_{RS}k_{SP}[P] \end{pmatrix}. \quad [15]$$

Thus, the *ratios* between these statistical weights must be $\frac{\rho_R}{\rho_S} = \frac{k_{SR}[R]k_{PS}}{k_{RS}k_{PS}} = \frac{k_{SR}[R]}{k_{RS}}$, and $\frac{\rho_P}{\rho_S} = \frac{k_{RS}k_{SP}[P]}{k_{RS}k_{PS}} = \frac{k_{SP}[P]}{k_{PS}}$.

Now we explicitly verify that given this special case of an acyclic architecture, these statistical weights are unchanged by imposing the further requirement of detailed balance. The condition of detailed balance is equivalent to stating that the input and output fluxes between any pair of nodes must equal,

$$\begin{cases} \rho_S k_{SR}[R] & = \rho_R k_{RS} \\ \rho_S k_{SP}[P] & = \rho_P k_{PS}. \end{cases} \quad [16]$$

We see at once that indeed, this statement of detailed balance is fully equivalent to the relative statistical weights we found by the Matrix Tree Theorem. (We need only consider $N - 1 = 2$ ratios in this case, by the normalization of total probability.) So as expected, the stationary probabilities found by the Matrix Tree Theorem further satisfy detailed balance, for this linear (acyclic) simple repression motif.

**C. The cycle condition relates a ratio of rate constants to (non)equilibrium.** In a graph composed of a single cycle of states, the net drive maintaining a nonequilibrium steady-state is related to the ratio of products of rate constants taken in opposing directions around the cycle (5). Here we pedagogically discuss this connection by showing that when this ratio is one, and the system is at steady-state, then the system must be at detailed balance, and vice versa.

Consider such a cyclic weighted graph composed of $N$ nodes and $2N$ edges (encoding the bidirectional transitions); enumerate the states from 1 to $N$, and the corresponding edge weights as the rates $k_{i,i+1}$ and $k_{i+1,i}$ between neighboring nodes $(i, i+1)$. (In what follows, given the cyclic structure of the graph, we adopt the notational convention that indices are to be taken modulo $N$.) For notational convenience, define the product of rate constants in the clockwise (+; increasing index $i$ direction) as

$$\gamma_+ \equiv \prod_{i=1}^{N} k_{i,i+1},$$

and the analogous product in the counter-clockwise direction as

$$\gamma_- \equiv \prod_{i=1}^{N} k_{i+1,i}.$$

Our goal is to show that when both their ratio $\gamma$ is unity,

$$\gamma \equiv \frac{\gamma_+}{\gamma_-} = \frac{\prod_{i=1}^{N} k_{i,i+1}}{\prod_{i=1}^{N} k_{i+1,i}} = 1 \quad [17]$$

and the system is at steady-state—namely that the net influxes and outfluxes balance for each node in graph,

$$0 = J_{i,i+1} - J_{i+1,i} + J_{i-1,i} - J_{i,i-1}, \forall i \in [\![1\ ;\ N]\!], \quad [18]$$

detailed balance is automatically satisfied, and vice versa. The detailed balance condition is that

$$J_{i,i+1} = k_{i,i+1}\rho_i = k_{i+1,i}\rho_{i+1} = J_{i+1,i}, \forall i \in [\![1\ ;\ N]\!]. \quad [19]$$

First, we verify the logical direction Detailed Balance, Eq. [19] $\Rightarrow$ (Steady State, Eq. [18] AND $\gamma = 1$, Eq. [17]). Rewriting the detailed balance condition Eq. [19] readily confirms this desired logical direction; specifically, we see,

$$\gamma = \frac{\prod_{i=1}^{N} k_{i,i+1}}{\prod_{i=1}^{N} k_{i+1,i}} = \frac{\Pi_{i=1}^{N} J_{i,i+1}}{\Pi_{i=1}^{N} J_{i+1,i}} = 1.$$

Next we verify the opposite logical direction, that Eq. [19] $\Leftarrow$ (Steady State, Eq. [18] AND $\gamma = 1$, Eq. [17]). Starting from the cycle condition of $\gamma = 1$ allows us to rewrite the influx through a given node $m$ as $J_{m+1,m} = \dfrac{\prod_{j=1}^{N} J_{j,j+1}}{\prod_{j=1,j\neq m}^{N} J_{j+1,j}}$. The outflux through a node $p$ is analogously $J_{p,p+1} = \dfrac{\prod_{j=1,j\neq p}^{N} J_{j,j+1}}{\prod_{j=1}^{N} J_{j+1,j}}$. Using these expressions to replace each of the four flux terms that appear in the steady-state condition Eq. [18], for all nodes $m \in \{i, i-1\}$ and $p \in \{i, i-1\}$, gives

$$0 = \frac{J_{12}...J_{i-1,i}J_{i+2,i+3}...J_{N1}}{J_{21}...J_{i,i-1}J_{i+1,i+2}...J_{1N}} \left[ \left( \frac{J_{i+1,i+2}}{J_{i+2,i+1}} - \frac{J_{i,i+1}}{J_{i,i+1}} \right) \frac{J_{i-1,i}}{J_{i,i-1}} - \frac{J_{i+1,i+2}}{J_{i+2,i+1}} \left( \frac{J_{i,i+1}}{J_{i+1,i}} - \frac{J_{i-1,i}}{J_{i,i-1}} \right) \right]. \qquad [20]$$

This expression simplifies to imply that the ratio of influxes to outfluxes must be the same across all pairs of edges, $\frac{J_{i-1,i}}{J_{i,i-1}} = \frac{J_{i+1,i+2}}{J_{i+2,i+1}} = H, \forall i \in [\![2\ ;\ N]\!]$, for some value $H$. Last, substituting the condition Eq. [17] implies that $H = 1$, and therefore implies Eq. [19], completing the desired correspondence.

**D. Discussion of various ways of quantifying dissipation.** The field of nonequilibrium thermodynamics quantifies nonequilibrium using different mathematical quantities. The nonequilibrium driving force, also referred to as the net (chemical) drive, is one key quantity. For a single cycle, the net drive $\Delta\mu$ is the net difference in chemical potential, namely free energy, imposed by one progression around the cycle along the nonequilibrium steady-state flux (5), (6, Ch. 13). For a single cycle, this net drive is related to the cycle parameter $\gamma$ we have just discussed in the previous subsection via

$$\Delta\mu = k_B T \ln\gamma. \qquad [21]$$

The units of this nonequilibrium driving force are energy ($k_B T$); in view of its centrality in describing nonequilibrium steady-states, this net drive is the quantity we use to analyze nonequilibrium in this paper.

Another related, central quantity that governs nonequilibrium behavior is the dissipation rate, or entropy production rate, which for a single cycle (at steady-state) is

$$\dot{W} \equiv \Delta J \Delta\mu = (J_{i,i+1} - J_{i+1,i})\ k_B T\ \ln\gamma, \qquad [22]$$

where $\Delta J$ is the nonequilibrium steady-state's net flux difference along any of the cycle graph's edges. This entropy production rate has units of work (energy per time).

Interestingly, note that Eq. [22] makes clear that even if a cycle requires a finite net drive $\Delta\mu \neq 0$ to maintain a nonequilibrium probability distribution over states, if the system is made to operate slowly enough—by reducing the magnitudes of all rates (hence fluxes $J$) simultaneously (while retaining their relative imbalances, e.g. in the same $\gamma$ and hence the same $\Delta\mu$)—the entropy production rate can be made arbitrarily small, $\dot{W} \to 0$. (Since our chief focus is on the statically controlled, steady-state behavior of regulatory systems, we do not analyze the entropy production rate in this paper, in favor of the net drive $\Delta\mu$.)

### 2. Insights into the square graph

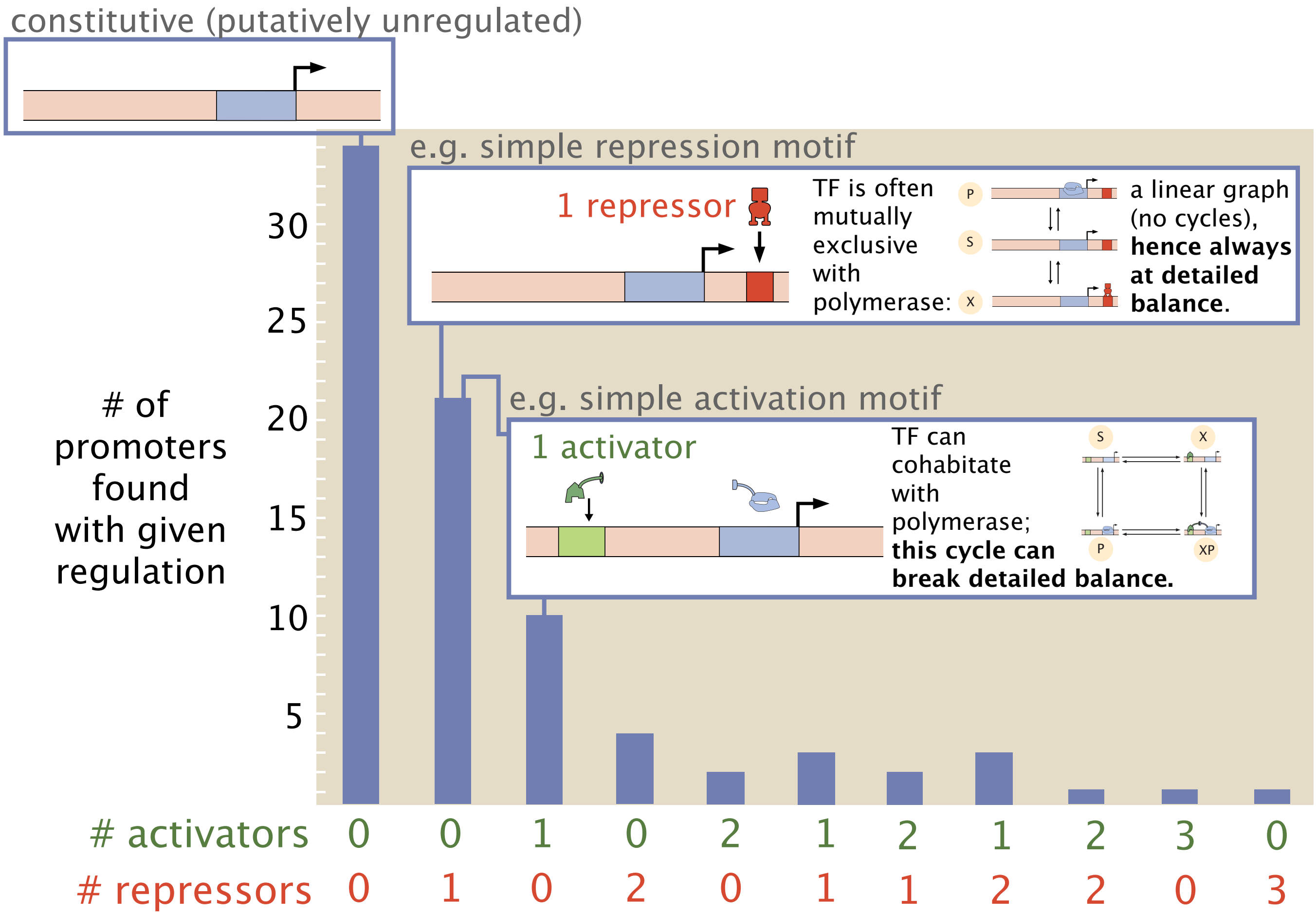

**Fig. S1.** An experimental histogram of empirically-observed gene regulatory motifs in *E. coli* (7) reveals that many promoter sites are regulated by a single repressor or activator. A single repressor can often implement the simple repression motif, where the repressor excludes the polymerase from binding, allowing just three states in a linear graph. (Raw histogram data are courtesy of Reference (7).) The reason that the "simple activation motif" is schematized as linked to both the (0 activator, 1 repressor) and (1 repressor, 0 activator) histogram bar is that while steric exclusion commonly occurs for repressors, often making single repressors well described by a linear graph of three states, some repressors do not completely exclude the polymerase, permitting a cycle motif too.

**A. The simple four-state cycle motif pervades prokaryotic and eukaryotic gene regulation.** Reference (7) is among the widest experimental censuses discovering regulatory interactions in *E. coli* in the recent literature. This study found that transcriptional architectures with one activator or repressor are the most commonly observed regulated transcriptional architectures. This pervasiveness of operons regulated by individual transcription factors is a finding confirmed by wider censuses based on aggregated studies in RegulonDB (8).

Thanks to common steric overlaps between the repressor binding site and polymerase binding site (8), a repressor is often—though not necessarily—mutually exclusive with the polymerase (see Fig. S1). In this case, the repressor implements a simple repression motif, a graph which lacks a cycle (3). However, when the repressor does not sterically exclude the polymerase, a cycle of four states emerges. The same cycle of four states emerges with activators, whose binding sites rarely directly overlap with the polymerase binding site (8); this produces a "simple activation motif." These observations affirm that a single cycle of four states is a common motif in prokaryotic gene regulation. Equivalents of such a cycle also occur regularly across eukaryotic gene regulation (9).

**B. Order of magnitude estimated rate constants for prokaryotic transcription.** Here, to allow us to assess how accessible interesting regulatory shape phenotypes are in the vicinity of biological rate parameters, we estimate default equilibrium rates typical of transcriptional systems in prokaryotes.

First, we remark that the correspondences between thermodynamic and kinetic viewpoints discussed in §G.3—specifically,

Eq. [60]—provides the following parameter correspondences useful for our estimates:

$$\begin{cases} K_1 = \frac{k_{PS}}{k_{SP}} = [P]\frac{N}{P}e^{\beta\Delta\epsilon_{pd}} \\ K_2 = \frac{k_{XP,P}}{k_{PX,P}} = [X]\frac{N}{X}e^{\beta(\Delta\epsilon_{xd}+\epsilon_{xp})} \\ K_3 = \frac{k_{XP,X}}{k_{X,XP}} = [P]\frac{N}{P}e^{\beta(\Delta\epsilon_{pd}+\epsilon_{xp})} \\ K_4 = \frac{k_{XS}}{k_{SX}} = [X]\frac{N}{X}e^{\beta\Delta\epsilon_{xd}} \end{cases}, \qquad [23]$$

where we have defined four equilibrium constants $K_1, \ldots, K_4$ (with units of concentration); $[X]$ is the concentration of transcription factor (say, in nanomolar) but $X$ is the absolute copy number (and analogously for the polymerase with $[P]$ and $P$); $N$ is the number of nonspecific binding sites on the genome; $\Delta\epsilon_{yd}$ energy difference between a state where molecule $Y$ is bound specifically to the genome versus nonspecifically; and $\epsilon_{xp}$ is the interaction energy between the transcription factor $X$ and the polymerase $P$. We note that $[X] = \frac{X}{N_A V_{cell}}$ and $[P] = \frac{P}{N_A V_{cell}}$, where $N_A$ is Avogadro's number and $V_{cell}$ is the volume of the cell, usually taken here to be that characteristic of *E. coli*, $V_{cell} \approx 1\ \mu m^3$. Noting that since one nanomolar is conveniently $1\ nM \approx \frac{1}{N_A V_{cell}}$, a natural unit for the rate constants depending on the concentration of transcription factor or polymerase is $s^{-1}nM^{-1}$.

Armed with these conventions, we now estimate the order of magnitude of governing rate constants from available measurements and empirical data.

- First, we consider plausible binding e.g. **on-rates** of polymerase or transcription factors to the genome.
  - Taking the Lac repressor as evocative of transcription factors, three empirical measurements give plausible on-rate values, and illustrate some empirical variation:
    - Ref. (10) (BNID 106392; (11)) reports a $k_{on} \approx 2.8 \times 10^7\ s^{-1}\ M^{-1} = \boxed{2.8 \times 10^{-2}\ s^{-1}\ nM^{-1}}$ for the Lac repressor.
    - Ref. (12) (BNID 104607; (11)) reports an appreciably larger association rate of $k_{on} \approx 7 \times 10^9\ s^{-1}\ M^{-1} = \boxed{7\ s^{-1}\ nM^{-1}}$.
    - In their SI, reference (13) report that they took measurements from a paper by Hammar *et al.* (14), who in their Fig. 2 report (from single molecule, *in vivo* measurements) that in *E. coli*, it takes the Lac repressor an average time of about $\tau_{on} \approx 30\ s$ to bind to O1 or Osym operator sites. The later reference (13) report without citation that the copy number of Lac repressors in this older paper's setting was in fact about 4 copies per cell ($\approx$ 4 nM). This implies an association rate of about $k_{on} \approx \frac{1}{\tau_{on}\ c} \approx \frac{1}{30\ s\ \times 4\ nM} \sim \boxed{10^{-2}\ s^{-1}\ nM^{-1}}$.

    An intermediate average of these various empirical data suggest a few tenths of a nanomolar per second is a reasonable scale for the basal association rate.
  - We compare the empirical measurements above with an order-of-magnitude theoretical estimate presuming diffusion-limited binding. RNAP's binding site is approximately $20 - 34$ bp long; each base-pair is separated by 3.4Å (15); so the characteristic scale $a$ we could expect of this binding site is about $a \approx 9$ nm. The diffusion constant of polymerase is $D_{poly} \approx 0.4\mu m^2/s$ (16), while the (effective, *in vivo*) diffusion coefficient for LacI is $D_{LacI} \sim 0.4\mu m^2$ (BNID 102038; (11); this effective diffusion constant for LacI plausibly reflects both 3D diffusion between nonspecific binding events and 1D genome-associated diffusion (17)). Reference (18) reports that the apparent (3D) diffusion coefficient of RNA polymerase II in the nucleus is (1-5) $\mu m^2/s$, similar to other transcription factors. (Altogether, these values indicate taking a diffusion constant of about $D \sim 1\ \mu m^2/s$ is reasonable.) A diffusion limited on-rate calculation then predicts that

$$k_{on} = 4\pi D a \sim 12(1\mu m^2/s)(9 \times 10^{-3}\ \mu m) \sim 0.11\ /s \underbrace{\mu m^3}_{(1/0.602)nM^{-1}} = 0.17/s/nM \sim \boxed{10^{-1}/s/nM}.$$

- Next we appraise characteristic energy scales among transcription factors, polymerase, and the specific sites on the genome:
  - According to Ref. (19) (BNID 103594; (11)), the polymerase binds more favorably to the Lac specific binding site than nonspecific sites on the genome by an energy difference of about $\Delta\epsilon_{pd} \approx -2.9\ k_BT$ so $\boxed{\beta\Delta\epsilon_{pd} \sim -3}$.
  - Ref. (20) reports that the Lac repressor preferentially binds to the specific operator binding sites with energies of ranging from $\Delta\epsilon_{xd} \approx -15.3\ k_BT$ (for the O1 site) to $\Delta\epsilon_{xd} \approx -9.7\ k_BT$ (for the O3 site). So we take as representative $\boxed{\beta\Delta\epsilon_{xd} \sim -13}$.
  - Ref. (19, Fig. 2) (BNID 103591; (11)) reports that the CRP activator interacts with RNAP with an interaction energy of approximately $\boxed{\beta\epsilon_{xp} \sim -4}$.

Since transcription factors plausibly stick to the genome by a factor $K_1/K_4 \approx \exp\left(-\beta\Delta\epsilon_{xd} + \beta\Delta\epsilon_{pd}\right) \sim \exp(13-3) = \exp(10) \sim 2 * 10^4$ stronger compared to the polymerase's interaction with the genome (19), we remark that any few-fold difference in the on-rate of polymerase to the genome (compared to the on-rate of the transcription factor to the genome) is not likely to be hugely significant in estimating $k_{off} = K_D k_{on}$. Therefore we will take the on-rates of polymerase and transcription factor to be essentially the same (diffusion-limited) value:

$$\boxed{k_{on} \sim 0.1/s/nM}.$$

- Considering *E. coli*, the number of nonspecific binding sites is about $N \approx 5 \times 10^6$ (19, 20) and the polymerase copy number is about $P \approx 10^3$ copies per cell (20). This suggests $[P] \approx 10^3\ nM$ and we estimate $k_{SP}[P] \equiv k_{X,XP}[P] \approx \left(0.1\ s^{-1}\ nM^{-1}\right)\left(10^3\ nM\right) \approx \boxed{10^2\ s^{-1}}$.
- While it is precisely how variation in the concentration $[X]$ tunes transcription that we are interested in, it is still instructive to report typical ranges for these transcription factor concentrations. As summarized in (15) (namely http://book.bionumbers.org/what-are-the-copy-numbers-of-transcription-factors/), cellular censuses show that repressing transcription factors typically have between $10 - 10^3$ copies per cell and activating transcription factors typically have between $1 - 10^2$ copies. This implies $[X] \sim$ few $\times 10^2$ nM. So ignoring the very variation in $[X]$ we're interested in, point estimates for $k_{SX}[X] \equiv k_{P,XP}[X]$ are $\approx (0.1\ s^{-1}\ nM^{-1})(\text{few} \times 10^2\ nM) \approx \boxed{\text{few} \times 10\ s^{-1}}$.

Altogether, these estimates enter to simplify Eq. 60 and imply an approximate, default set of all rates. We summarize these order of magnitude values in Figure 1A of the main text and the table S1 below. (In the later analyses examining the consequences of drive along individual edges or pairs of edges, we choose and analyze more precise sets of default rate values consistent with these orders of magnitude; see Figures S10 and K.1.)

| rate | meaning | calculation | order of magnitude estimate |
|---|---|---|---|
| $k_{XS}$ | unbinding of TF from empty genome | $k_{SX}e^{\beta\Delta\epsilon_{xd}}$ | $0.8\ s^{-1}$ |
| $k_{XPP}$ | unbinding of TF from RNAP-bound genome | $k_{PXP}N(1\ nM)e^{\beta(\Delta\epsilon_{xd}+\epsilon_{xp})}$ | $2 \times 10^{-2}\ s^{-1}$ |
| $k_{SX}$ | binding of TF to empty genome | $:= k_{on}$ | $0.1\ s^{-1}\ nM^{-1}$ |
| $k_{PXP}$ | binding of TF to RNAP-bound genome | $:= k_{on}$ | $0.1\ s^{-1}\ nM^{-1}$ |
| $k_{PS}$ | unbinding of RNAP from empty genome | $k_{SP}N(1\ nM)e^{\beta\Delta\epsilon_{pd}}$ | $2 \times 10^4\ s^{-1}$ |
| $k_{XPX}$ | unbinding of RNAP from TF-bound genome | $k_{XXP}N(1\ nM)e^{\beta(\Delta\epsilon_{pd}+\epsilon_{xp})}$ | $5 \times 10^2\ s^{-1}$ |
| $k_{SP}$ | binding of RNAP to empty genome | $:= k_{on}$ | $0.1\ s^{-1}\ nM^{-1}$ |
| $k_{XXP}$ | binding of RNAP to TF-bound genome | $:= k_{on}$ | $0.1\ s^{-1}\ nM^{-1}$ |

**Table S1. Summary of orders-of-magnitude estimates of rates at equilibrium that govern transcription.**

#### C. Biologically, timescales are plausibly separated enough that transcription is well represented by small Markov chain graphs.

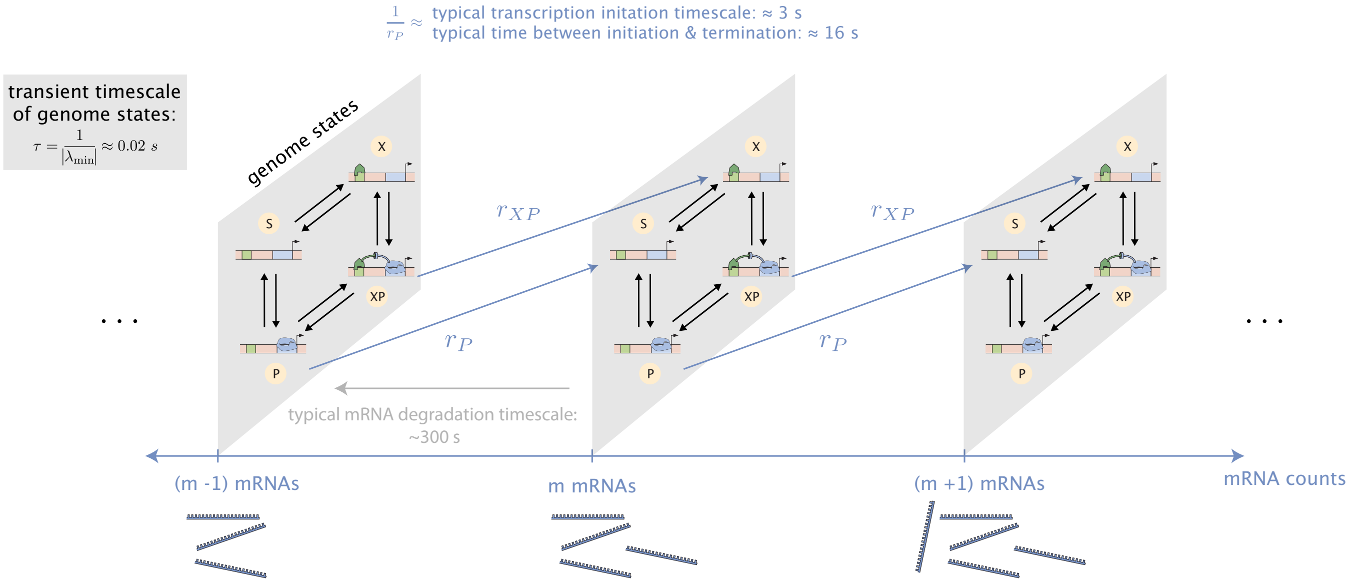

The magnitudes of eigenvalues **λ**<0 of the Laplacian **L** are decay rates that set how slowly p(t) transiently approaches steady state; this decay is dominated by the slowest rate $\lambda_{\min}$.

Assuming that the abundance of the polymerase is about [X] ~ 200 copies/cell = 200 nM; and the polymerase concentration is [P] ~ $10^3$ copies/cell = $10^3$ nM, the Laplacian is

$$\rightarrow \mathbf{L}_{\text{genome}} = \begin{pmatrix} -120 & 0.8 & 0 & 20000 \\ 20 & -100.8 & 500 & 0 \\ 0 & 100 & -500.02 & 20 \\ 100 & 0 & 0.02 & -20020 \end{pmatrix} s^{-1}$$

(columns: source state S, X, XP, P; rows: destination state S, X, XP, P)

and computing the eigenvalues, we find that the smallest decay rate $\lambda_{\min} \approx -20\ s^{-1}$ is fast relative to transcription or degradation.

**Fig. S2.** A separation of timescales between transcription and binding or unbinding is well justified, for the order-of-magnitude rate constant estimates we adopt to model transcription.

Technically, gene expression is governed by a fuller chemical master equation than that defined by merely the states of the genome. In principle, the current number of mRNA transcripts could affect the allowed transitions, and *a priori* one might worry that an additional mechanism to transition from a state where the polymerase is bound to the genome ($P$ or $XP$) to a state where it is unbound ($S$ or $X$) is when the polymerase has transcribed a transcript successfully enough to vacate the polymerase binding site. These technicalities would in fact imply a larger, fuller ladder of states that define the discrete state Markov chain, as visualized in Figure S2. However, here we argue that the time to transcribe is typically much longer than the equilibration timescale of the four states of the genome alone. This separation of timescales formally justifies the assumption that the net accumulation of mRNA transcripts is proportional to the probability of being in the polymerase-bound states.

First, we estimate the rate at which the count of mRNA transcripts accumulate once the polymerase is bound. RNAP elongates nascent transcripts at a rate of about 3.72 kb/min in *E. coli* (BNID 103021; (11)); this is $v = 62$ nucleotides/second. The average protein is $L_p \approx 340$ peptides long (BNID 10895; (11)), implying that protein-coding mRNAs are about $3L_P \approx 10^3$ nucleotides long, consistent with reports elsewhere of mean mRNA lengths of 924 nt across prokaryoates (21). Hence, once transcribing, it takes approximately $\tau_{\text{transcribe}} \approx \frac{3L_p}{v} \approx 1000/62 \approx 16$ seconds to serially transcribe a typical gene. This is a lower bound on the accumulation rate, however, since the RNAP can leave the promoter faster than a transcript is complete, permitting a larger transcription initiation rate. In *E. coli*, transcription iniatation has been reported to occur at a typical rate of 20 initiations/min/gene, or at a rate of $\sim$ 0.3initiations per second (BNID 111997; (11)). Therefore, in the fuller lattice of states of a Markov chain explicitly tracking mRNA counts (Fig. S2), the rates of transitions from states with count $m$ to states with count $m+1$ are plausibly between the lower bound of $r_{\text{transcribe}} = 1/\tau_{\text{transcribe}} \approx 1/16s \approx 0.06\ s^{-1}$ and an upper bound of $r_{\text{initiate}} \approx 0.3\ s^{-1}$, or in summary, we take $r \sim \boxed{\text{few} \times 10^{-1}\ s^{-1}}$. In addition, degradation is even slower: the typical half-life of an mRNA in *E. coli* is reported to be on the order of a few minutes (BNID 108598; (11)), implying the degradation rate (governing how quickly $m$ mRNAs could decrement to $m-1$ mRNAs) is on the order of $\gamma_d \sim \text{few} \times 10^{-3}\ s^{-1}$.

In contrast, the slowest timescale within which the four genome states converge towards their steady-state distribution—set by the smallest magnitude eigenvalue of the four state Laplacian matrix of transition rates for the genome—is approximately $1/20 \approx 0.05$ seconds (see Fig. S2 for the calculation). This is much faster than the transcriptional transition timescales. Therefore, the condensation of the larger ladder graph into the smaller graph of just four binding and unbinding reactions on the genome is justified, for this particular set of plausible rate constants.

**D. Deriving the universal form: The Matrix Tree Theorem on the square graph yields a ratio of quadratic polynomials.** Applying the Matrix Tree Theorem to derive steady-state probabilities $p_i$ of each state $i$, and hence any response observable $\langle r\rangle \equiv \sum_{\text{states } i} r_i p_i$, reveals that these responses follow the following universal form,

$$\boxed{\langle r\rangle = \frac{A + B[X] + C[X]^2}{D + E[X] + F[X]^2}}, \tag{24}$$

where the coefficients are given by weighted sums of spanning trees with different possible $[X]$-dependencies, namely,

$$\begin{cases} A = r_P T_P^0 + r_S T_S^0 \\ B = r_P T_P^1 + r_S T_S^1 + r_{XP} T_{XP}^1 + r_X T_X^1 \\ C = r_{XP} T_{XP}^2 + r_X T_X^2 \\ D = T_P^0 + T_S^0 \\ E = T_P^1 + T_S^1 + T_{XP}^1 + T_X^1 \\ F = T_{XP}^2 + T_X^2, \end{cases} . \tag{25}$$

Here, $T_Y^n[X]^n$ is the sum of spanning trees rooted in $Y$ where $n$ edges depend on $[X]$ participate. For example, $T_{XP}^1 = k_{SP}[P]k_{PXP}k_{XXP}[P] + k_{PS}k_{SX}k_{XXP}[P] + k_{XS}k_{SP}[P]k_{PXP}$ is the sum of all spanning trees rooted in state $XP$ that carry a linear $[X]$-dependence. The other explicit expressions of the coefficients $T_Y^n$ are visualized in Figure S3.

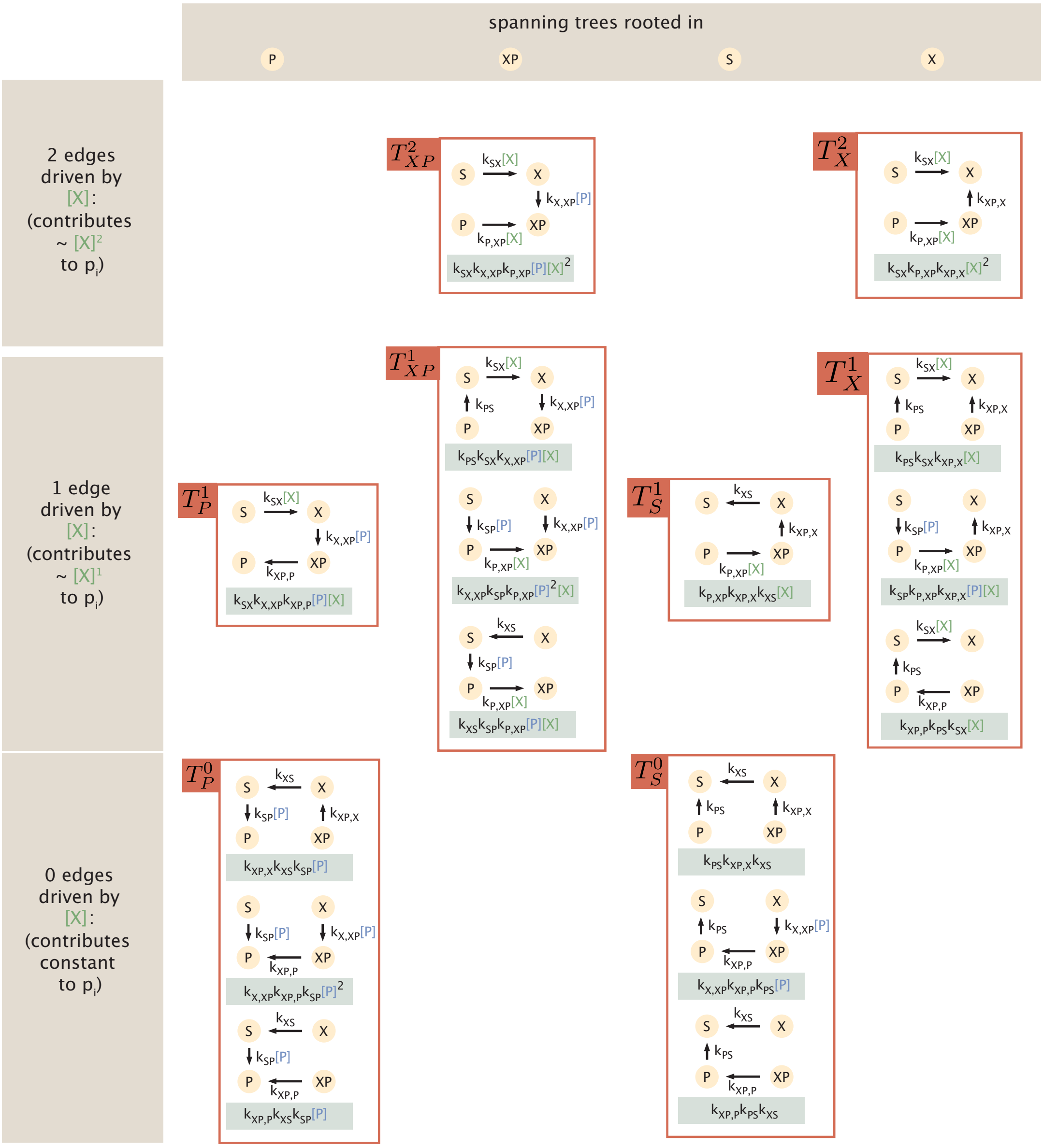

**Fig. S3.** All 16 rooted spanning trees of the four-state cycle can be classified by which node serves as the root (in columns) and the participating number of edges that contribute a power $n$ of the transcription factor concentration $[X]$ (in row $n$). The weighted spanning trees completely determine the universal form of the fold-change output, as specified by Eqs.24 and 25.

**E. Discussion on observable conventions: the logarithmic control variable.** Throughout our analysis and discussion in this paper, we monitor the shape, number of inflection points, and sensitivity of transcriptional outputs with respect to the control parameter of the concentration of transcription factor, on a *logarithmic* scale. We use this logarithmic convention in alignment with common practice in biochemical and transcriptional studies (1, 20, 22). Using log concentration is convenient in the common setting where environmental inputs or governing transcription factor concentrations can vary over orders of magnitude, or where biochemical control systems are conceptually implementing a sort of fold-change detection (23).

This logarithmic convention is largely benign, since it is grounded in a monotonic one-to-one transformation of the control variable measured on a linear scale; however, it has two small mathematical consequences we briefly appraise. First, counting the number of inflection points with respect to the *logarithmic* control variable can introduce an additional point of inflection compared to the linear control variable. This occurs for the discussions of the shape of detailed balance responses,

$$\langle r \rangle^{\mathrm{eq}} = \frac{A' + B'[X]}{C' + D'[X]}. \qquad [26]$$

This is famously just a Langmuir binding curve or Hill function of order one, which on a linear scale is a hyperbola (nonsigmoidal and without any inflection points). However, it is quite common to depict such curves on a logarithmic scale, where the curve gains sigmoidal character and a point of inflection; the inflection point's local slope defines an effective Hill coefficient. This canonical view, with respect to a logarithmic control variable, is the picture we invoke while counting inflection points or describing shapes.

Second, taking a logarithm invites a mathematical comment on units. Any logarithm of a concentration control variable must be understood as a logarithm of that concentration relative to some standard concentration scale, for instance 1 nanomolar. In plots where $\log[X]$ appears, the reference concentration merely denotes the horizontal offset/position of the curve. The particular choice of such a standard reference concentration scale $[X]_0$ has no effect on logarithmic derivatives, because of the simple fact that

$$\frac{df(x)}{d\log([X]/[X]_0)} = \frac{df(x)}{d(\log[X] - \log[X]_0)} = \frac{df(x)}{d\log[X]}. \tag{27}$$

**F. Collapse of eight parameters into two emergent fundamental shape parameters $(a, b)$.** Now, by neglecting scales and shifts, we show how we can reduce the ratio of quadratic polynomials Eq. [25]—possessing six coefficients that are functions of eight rate constants—to an emergent form of just two shape parameters, namely:

$$\boxed{\frac{\langle r\rangle - \langle r\rangle_0}{\langle r\rangle_\infty - \langle r\rangle_0} = \frac{ax + x^2}{1 + bx + x^2}}, \tag{28}$$

where $\langle r\rangle_0$ and $\langle r\rangle_\infty$ are the leakiness and saturation of the function, expressible in terms of ratios of coefficients: $\lim_{[X]\to 0} \langle r\rangle = \langle r\rangle_0 = \frac{A}{D}$ and $\lim_{[X]\to\infty} \langle r\rangle \equiv \langle r\rangle_\infty = \frac{C}{F}$.

To show this two-parameter form of Eq. 28, we preview our procedure as follows. We divide by one of the six original coefficients of Eq. 24 (here, the coefficient $D$); extract an additive factor of the leakiness $\langle r\rangle_0$; nondimensionalize the concentration $[X]$ by a convenient concentration scale that emerges; perceive that a multiplicative factor of the dynamic range $\langle r\rangle_\infty - \langle r\rangle_0$ can be demanded to appear; and summarize the resulting expression by defining just two emergent shape parameters. To wit,

$$\langle r\rangle = \frac{A + B[X] + C[X]^2}{D + E[X] + F[X]^2} \tag{29}$$

$$= \frac{\frac{A}{D} + \frac{B}{D}[X] + \frac{C}{D}[X]^2}{1 + \frac{E}{D}[X] + \frac{F}{D}[X]^2} \tag{30}$$

$$= \langle r\rangle_0 + \frac{\frac{A}{D} + \frac{B}{D}[X] + \frac{C}{D}[X]^2 - \langle r\rangle_0(1 + \frac{E}{D}[X] + \frac{F}{D}[X]^2)}{1 + \frac{E}{D}[X] + \frac{F}{D}[X]^2} \tag{31}$$

$$= \langle r\rangle_0 + \frac{(\frac{B}{D} - \langle r\rangle_0\frac{E}{D})[X] + (\frac{C}{D} - \langle r\rangle_0\frac{F}{D})[X]^2}{1 + \frac{E}{D}[X] + \frac{F}{D}[X]^2} \tag{32}$$

Now we nondimensionalize the control parameter by a convenient concentration scale, $[X]_0 = \sqrt{\frac{D}{F}}$, thus expressing the observable with respect to the rescaled concentration variable, $x \equiv \frac{[X]}{[X]_0}$:

$$\langle r\rangle = \langle r\rangle_0 + \frac{\frac{E}{\sqrt{DF}}(\frac{B}{E} - \langle r\rangle_0)x + (\frac{C}{F} - \langle r\rangle_0)x^2}{1 + \frac{E}{\sqrt{DF}}x + x^2} \tag{33}$$

As long as $\langle r\rangle_\infty \neq \langle r\rangle_0$, a condition we will consider shortly, we can rewrite this form of the observable as

$$\langle r\rangle = \langle r\rangle_0 + (\langle r\rangle_\infty - \langle r\rangle_0)\frac{\frac{E}{\sqrt{DF}}\frac{\frac{B}{E} - \langle r\rangle_0}{\langle r\rangle_\infty - \langle r\rangle_0}x + x^2}{1 + \frac{E}{\sqrt{DF}}x + x^2}. \tag{34}$$

Finally, this form invites us to define shape parameters $a, b$ as

$$\begin{cases} b = \frac{E}{\sqrt{DF}} \\ a = b\frac{\frac{B}{E} - \langle r\rangle_0}{\langle r\rangle_\infty - \langle r\rangle_0}, \end{cases} \tag{35}$$

and allows us to write

$$\langle r\rangle = \langle r\rangle_0 + (\langle r\rangle_\infty - \langle r\rangle_0)\frac{ax + x^2}{1 + bx + x^2}, \tag{36}$$

recovering the simplified expression Eq. [28].

Now we return to address the assumption that $\langle r\rangle_0 \neq \langle r\rangle_\infty$, i.e. that the uninduced response (leakiness) is different from the maximally induced response (saturation). If instead we are in the unusual special case that the response does not change with $[X]$ at all, we extend $a$ by continuity to $a = b$. In this constant case, $\frac{B}{E} - \langle r\rangle_0 = \langle r\rangle_\infty - \langle r\rangle_0$. In fact the function is constant when $\frac{B}{E} = \frac{A}{D} = \frac{C}{F}$ and the whole polynomial of order two factors out. Is this limit, $\frac{a}{b} \to 1$, and the form of equation Eq. [28] still holds.

Otherwise, if $\langle r \rangle_0 = \langle r \rangle_0$ but the function is *not* constant everywhere, $a$ is infinite and the proper simplified parameterization of the observable instead becomes $\langle r \rangle = \langle r \rangle_0 + \frac{cx}{1+bx+x^2}$, with $c = b(\frac{B}{E} - \langle r \rangle_0)$. In this case, the function is non-monotonic. Indeed, the function has to both increase and decrease to have the same limit at zero and infinity without being constant. We do not make an elaborate quantitative study of this class of function, because we propose that in biological systems that succeed at accomplishing regulation, it is usually the case that the uninduced and maximally induced responses are at least infinitesimally different, namely $|\langle r \rangle_\infty - \langle r \rangle_0| = \epsilon$ with $\epsilon$ finite. However, philosophically, this type of eccentric response is still accommodated by the parameterization of Eq. [28] in the limit that $a \to \infty$.

### G. Equilibrium responses of the square graph.

***G.1. Demotion of responses to a (monotonic) ratio of linear polynomials at equilibrium.*** Here, we derive Eq. 3 of the main text (also reproduced here as Eq. [26]), that any observable produced by the square graph is demoted to a ratio of *linear* polynomials in $[X]$ at detailed balance. Informally, our strategy will be to factor out a statistical weight of a particular reference state from every statistical weight that participates in defining the observable $\langle r \rangle$; this forces ratios of statistical weights to appear, which the detailed balance condition relates to ratios of rate constants. In the square graph, the ratios of rate constants can carry only a single power of $[X]$, motivating the appearance of linear terms only. (Along the quick mathematical journey, we will resolve the minor mathematical wrinkle that the detailed balance condition only comments immediately on the ratio of two statistical weights when those states are connected in the graph.)

We proceed. Choose the reference state to be state $P$, for concreteness though arbitrarily (as long as this reference state has nonzero steady-state probability). We can write,

$$\langle r \rangle = \sum_i r_i p_i \qquad [37]$$

$$= \frac{\sum_i r_i \rho_i}{\sum_i \rho_i} \qquad [38]$$

$$= \frac{\rho_P \sum_i r_i \frac{\rho_i}{\rho_P}}{\rho_P \sum_i \frac{\rho_i}{\rho_P}} \qquad [39]$$

$$= \frac{r_P + \sum_{\substack{\text{connected} \\ i \neq P}} r_i \frac{k_{Pi}}{k_{iP}} + \sum_{\substack{\textbf{dis}\text{connected} \\ j \neq P}} r_j \frac{\rho_j}{\rho_P}}{1 + \sum_{\substack{\text{connected} \\ i \neq P}} \frac{k_{Pi}}{k_{iP}} + \sum_{\substack{\textbf{dis}\text{connected} \\ j \neq P}} \frac{\rho_j}{\rho_P}} \qquad [40]$$

Why does the last line have separated sums? This is the mathematical wrinkle we alluded to. Detailed balance guarantees that $\rho_i k_{iP} = \rho_P k_{Pi}$ for any state $i$. Normally, if the rates are nonzero, this suggests we can replace a ratio of statistical weights by a ratio of rate constants (the first sum). However, if a state $j$ is *not* connected to $P$ (namely $k_{jP} = k_{Pj} = 0$), then we can no longer necessarily write $\frac{\rho_j}{\rho_P}$ as a pure ratio of just two rate constants.

To make further progress, we consider the second sum in the numerator, whose summands are those ratios $\rho_j/\rho_P$ for states $j$ that are not connected to $P$. By the strongly-connected structural assumption that empowers us to apply the Matrix Tree Theorem, there must be at least one path (built from some number $q$ of edges in the graph) that connects state $j$ to state $P$. Hence, the ratio of statistical weights can be written as a product of rate ratios along that path, giving

$$\frac{\rho_j}{\rho_P} = \frac{\rho_j}{\rho_a}\frac{\rho_a}{\rho_b}\frac{\rho_b}{\rho_c}\cdots\frac{\rho_r}{\rho_q}\frac{\rho_q}{\rho_P} \qquad [41]$$

$$= \underbrace{\frac{k_{aj}}{k_{ja}}\frac{k_{ba}}{k_{ab}}\frac{k_{cb}}{k_{bc}}\cdots\frac{k_{qr}}{k_{rq}}\frac{k_{Pq}}{k_{qP}}}_{q \text{ ratios}}. \qquad [42]$$

Since here, each directed edge carries at most a linear factor of $[X]$, any ratio of rate constants is either constant; proportional to $1/[X]$; or proportional to $[X]$.

Returning to the specifics of the four-state graph and our reference state $P$, we see that states $S$ and $XP$ are both connected to $P$, giving the first, connected-state sum as $\sum_{\substack{\text{connected} \\ i \neq P}} r_i \frac{k_{Pi}}{k_{iP}} = r_S \frac{k_{PS}}{k_{SP}[P]} + r_{XP} \frac{k_{PXP}[X]}{k_{XPP}}$.

The only state that is disconnected from state $P$, giving the disconnected sum, is state $X$. Without loss of generality, we now rewrite $\frac{\rho_X}{\rho_P}$ using the path of edges that goes through $S$. (We recover the same ultimate $[X]$-dependency if we had chosen the path through $XP$ instead.) This gives,

$$\frac{\rho_X}{\rho_P} = \frac{\rho_X}{\rho_S}\frac{\rho_S}{\rho_P} \tag{43}$$

$$= \frac{k_{SX}[X]}{k_{XS}}\frac{k_{PS}}{k_{SP}[P]}. \tag{44}$$

So the disconnected sum is just $\sum_{\substack{\text{disconnected}\\ j\neq P}} r_j \frac{\rho_j}{\rho_P} = r_X \frac{k_{SX}[X]}{k_{XS}}\frac{k_{PS}}{k_{SP}[P]}$. Altogether, we recover

$$\langle r\rangle^{\text{eq.}} = \frac{r_P + \left(r_S \frac{k_{PS}}{k_{SP}[P]} + r_{XP}\frac{k_{PXP}[X]}{k_{XPP}}\right) + \left(r_X \frac{k_{SX}[X]}{k_{XS}}\frac{k_{PS}}{k_{SP}[P]}\right)}{1 + \left(\frac{k_{PS}}{k_{SP}[P]} + \frac{k_{PXP}[X]}{k_{XPP}}\right) + \left(\frac{k_{SX}[X]}{k_{XS}}\frac{k_{PS}}{k_{SP}[P]}\right)} \tag{45}$$

$$:= \boxed{\frac{A' + B'[X]}{C' + D'[X]}}, \tag{46}$$

where we have highlighted how both the numerator and denominator admit only up to a linear dependence on $[X]$, and $A', B', C', D'$ are coefficients that depend only on weighted ratios of opposing rate constants (and are hence set fully thermodynamically by energy parameters).

The reasoning above suggests that the fact that every path connecting two states contains at most one power of $[X]$ was a crucial architectural ingredient for the collapse of the ratio of quadratic polynomials to a ratio of linear polynomials in the square graph. One interesting transparent consequence this reasoning highlights is that the same collapse (to a ratio of linear polynomials at detailed balance) must occur for the completely-connected graph.

***G.2. Leakiness, saturation, and EC50 are tunable at equilibrium.*** As mentioned in the main text, the response's leakiness (value when $[X]$ is completely absent) and saturation (value when $[X] \to \infty$) are set by the fact that the four state graph collapses into a different two-state linear graph for each limit. Specifically, the kinetics reduce to,

$$\begin{cases} [X] \to 0: & S \underset{k_{PS}}{\overset{k_{SP}[P]}{\rightleftharpoons}} P \\ [X] \to \infty: & X \underset{k_{XP,X}}{\overset{k_{X,XP}[P]}{\rightleftharpoons}} XP \end{cases}. \tag{47}$$

Since these two-state truncated graphs are linear, and so must be at equilibrium, we observe that the values of the leakiness and saturation must be thermodynamic statistical averages of the $r_i$. We conclude that

$$\begin{cases} \langle r\rangle_0 = r_P p_P + r_S\,(1 - p_P) \\ \langle r\rangle_\infty = r_{XP} p_{XP} + r_X\,(1 - p_{XP}), \end{cases} \tag{48}$$

where $p_P = \frac{k_{SP}[P]}{k_{SP}[P] + k_{PS}} \equiv \frac{1}{1+e^{-\beta\Delta\epsilon_{SP}}}$ and $p_{XP} = \frac{k_{X,XP}[P]}{k_{X,XP}[P] + k_{XP,X}} \equiv \frac{1}{1+e^{-\beta\Delta\epsilon_{XXP}}}$ are the simple stationary-solutions of each two-state system, and where we have defined the appropriate Boltzmann energy parameters via each ratio of rates. Hence leakiness and saturation are controllable by thermodynamic means.

Further assessing the form of the inflection point when the observable is at detailed balance reveals that it can be set by another ratio of rates, hence energy parameter. However, the raw sharpness at the inflection point remains equal to one fourth of the dynamic range. We demonstrate this obligatory proportionality between the maximum raw sharpness and dynamic range as follows. At equilibrium, taking one derivative of the detailed balance response described by Eq. [46] gives the raw sharpness as,

$$\frac{d\langle r\rangle^{\text{eq.}}}{d(\ln[X]/[X]_0)} = \frac{(B'C' - A'D')[X]}{(C' + D'[X])^2}. \tag{49}$$

Taking an additional derivative to solve for the inflection point where $\frac{d^2\langle r\rangle^{\text{eq.}}}{d(\ln[X]/[X]_0)^2} = 0$ gives,

$$\frac{d^2\langle r\rangle^{\text{eq.}}}{d(\ln[X]/[X]_0)^2} = \frac{(B'C' - A'D')(C' - D'[X])[X]}{C' + D'[X]}. \tag{50}$$

The inflection point, where this second derivative vanishes and the raw sharpness is maximized, occurs at $[X]_* = C/D$. Substituting this into the maximal sharpness expression, we find the maximum sharpness at equilibrium is merely

$$\max \frac{d\langle r\rangle^{\text{eq.}}}{d(\ln[X]/[X]_0)} = \frac{1}{4}\left(\frac{B'}{D'} - \frac{A'}{C'}\right). \tag{51}$$

Now, note that the equilibrium leakiness is given by

$$\langle r \rangle_0^{\text{eq}} \equiv \lim_{[X]\to 0} \langle r \rangle^{\text{eq}} = \frac{A'}{C'}, \qquad [52]$$

and the saturation is given by

$$\langle r \rangle_\infty^{\text{eq}} \equiv \lim_{[X]\to \infty} \langle r \rangle^{\text{eq}} = \frac{B'}{D'}, \qquad [53]$$

so the maximum sharpness is indeed one fourth the dynamic range,

$$\max \frac{d\langle r \rangle^{\text{eq.}}}{d(\ln[X]/[X]_0)} = \frac{1}{4}\left(\langle r \rangle_\infty^{\text{eq}} - \langle r \rangle_0^{\text{eq}}\right). \qquad [54]$$

These constrained behaviors of the equilibrium response are summarized in Figure S4.

A transcription factor is a global, overall repressor when the saturation is smaller than the leakiness, $\langle r \rangle_\infty < \langle r \rangle_0$. Conversely, a transcription factor is overall an activator when the saturation is larger than the leakiness, $\langle r \rangle_\infty > \langle r \rangle_0$. As we have just seen, since the leakiness and saturation are set thermodynamically, so too is the global nature of the transcription factor as an overall repressor or activator.

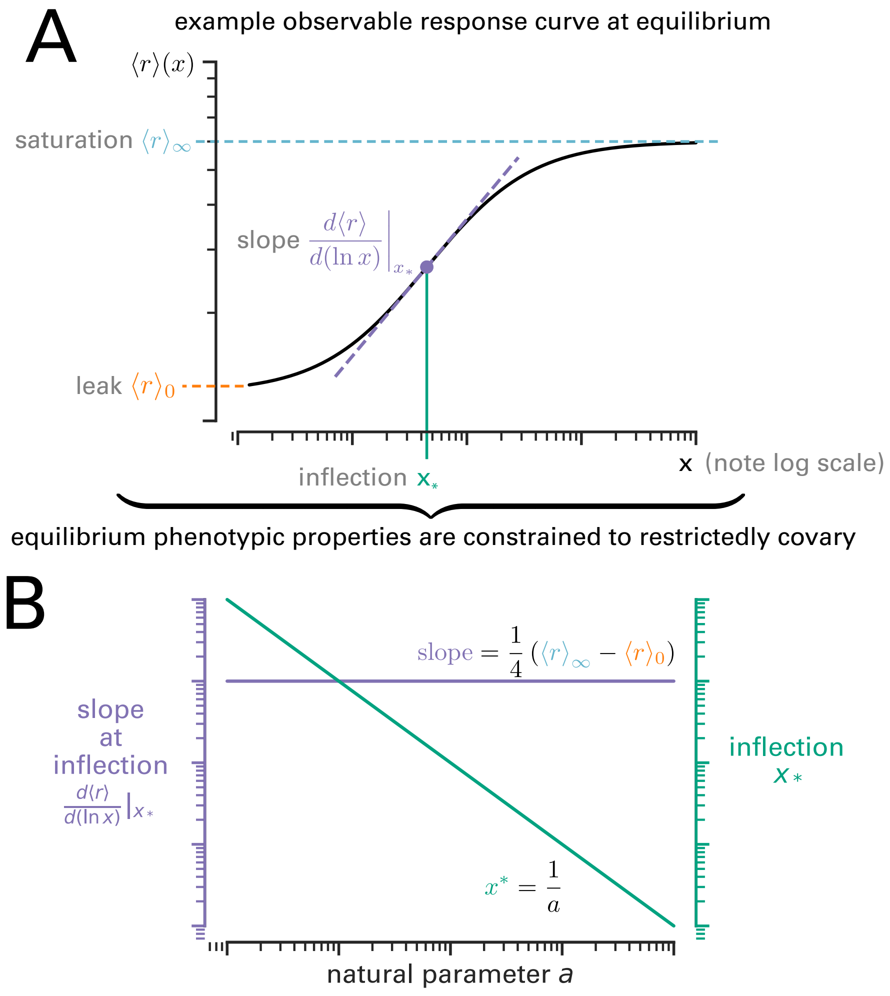

**Fig. S4.** At equilibrium, response curves (A) are always monotonic in the control variable $x$, with (at most) one inflection point in $\ln x$. The leak (observable at zero $x$, $\langle r \rangle_0$, in orange); location $x_*$ of the inflection point (in green); slope at the inflection (in purple); and saturation limit (in pale blue) capture the properties of the curve. Equilibrium imposes the constraint that these phenotypic properties vary in fixed relationships, as illustrated in (B).

***G.3. Validating consilience between kinetic and thermodynamic viewpoints.*** To be helpful to the reader interested in reconciling thermodynamic models; experimental parameters such as equilibrium dissociation constants that may parameterize them; and the more elaborate kinetic parameterization of continuous-time Markov chains and the Matrix Tree Theorem, below we endeavor a parameter-by-parameter correspondence between these viewpoints. This correspondence is valid when energy dissipation vanishes.

From a kinetic viewpoint, detailed balance implies that the ratio of two states is expressible as a ratio of rate constants. From a thermodynamic viewpoint, the same ratio of two states is expressible as a ratio of Boltzmann weights set by thermodynamic energy parameters. To link these perspectives, we define an effective equilibrium dissociation constant between a molecule $Y$ and a site $H$, where the site can either be completely empty or also occupied by another molecule in its vicinity. We denote these equilibrium constants $K_{HY,H}$ and largely following the conventions discussed in Ref. (19), define them as

$$K_{HY,H} = \frac{[Y]}{y^H}, \quad [55]$$

where $y^H = \frac{\rho_{HY}}{\rho_H}$ is a ratio of statistical weights; specifically, $\rho_{HY}$ is statistical weight of the molecule $Y$ bound to the site $H$, and $\rho_H$ is the statistical weight of the state where the molecule $Y$ is not bound to the site $H$. With this definition, the ratio of probabilities of two states is constant and the dissociation constant has units of a concentration.

For the square graph of four states—namely when the site is empty, $S$; when the transcription factor is bound to the DNA, $X$; when the polymerase is bound to the DNA, $P$; and when both the transcription factor and the polymerase are both bound to the DNA, $XP$—we can define the effective equilibrium dissociation constants explicitly, seeing,

$$\begin{cases} K_{SP,S} = \frac{[P]}{p^S} = \frac{[P]\rho_S}{\rho_P} \\ K_{XP,P} = \frac{[X]}{x^P} = \frac{[X]\rho_P}{\rho_{XP}} \\ K_{XP,X} = \frac{[P]}{p^X} = \frac{[P]\rho_X}{\rho_{XP}} \\ K_{SX,S} = \frac{[X]}{x^S} = \frac{[X]\rho_S}{\rho_X}. \end{cases} \quad [56]$$

In the 4-state-graph, detailed balance implies that,

$$\begin{cases} \frac{\rho_X}{\rho_S} = \frac{[X]k_{SX}}{k_{XS}} \\ \frac{\rho_P}{\rho_S} = \frac{[P]k_{SP}}{k_{PS}} \\ \frac{\rho_X}{\rho_{XP}} = \frac{k_{XP,X}}{[P]k_{XP,X}} \\ \frac{\rho_P}{\rho_{XP}} = \frac{k_{XP,P}}{[X]k_{XP,P}}. \end{cases} \quad [57]$$

So we can express the effective equilibrium dissociation constants as functions of rate constants, recovering,

$$\begin{cases} K_{SP,S} = \frac{k_{PS}}{k_{SP}} \\ K_{XP,P} = \frac{k_{XP,P}}{k_{XP,P}} \\ K_{XP,X} = \frac{k_{XP,X}}{k_{XP,X}} \\ K_{SX,S} = \frac{k_{XS}}{k_{SX}}. \end{cases} \quad [58]$$

Similarly, we can derive their expression with the thermodynamic formalism. Referring to Reference (19), we can define the partition function of the 4 states characterising the simple activation as follows: $Z(P,X) = \frac{N!}{P!X!(N-P-X)!}e^{-P\beta\epsilon^{ns}_{pd}/k_bT - X\beta\epsilon^{ns}_{xd}}$, where $\beta = 1/k_BT$, $k_B$ is the Boltzmann constant, and $T$ the temperature. For the transcription case we can define $\Delta\epsilon_{yd} = \epsilon^s_{yd} - \epsilon^{ns}_{yd}$, where $\epsilon^s_{yd}$ is the energy of the molecule $Y$ being on a specific site and $\epsilon^{ns}_{yd}$ the energy of the molecule being on a non specific site and $\epsilon_{xp}$ the interaction energy between the transcription factor and the polymerase. $X$ and $P$ are respectively the number of sites free on the DNA for the transcription factor and the polymerase to bind. $N$ is the number of nonspecific binding sites. We can define the weights of the different nodes at thermodynamic equilibrium (19):

$$\begin{cases} \rho_S = Z(P,X) \\ \rho_P = Z(P-1,X)e^{-\beta\epsilon^s_{pd}} \\ \rho_{XP} = Z(P-1,X-1)e^{-\beta(\epsilon^s_{pd}+\epsilon^s_{xd}+\epsilon_{px})} \\ \rho_X = Z(P,X-1)e^{-\beta\epsilon^s_{xd}} \end{cases} \quad [59]$$

Using the statistical mechanics approximation ($N \gg P, X$), we compute the effective equilibrium dissociation constants:

$$\begin{cases} K_{SP,S} = [P]\frac{N}{P}e^{\beta\Delta\epsilon_{pd}} \\ K_{XP,P} = [X]\frac{N}{X}e^{\beta(\Delta\epsilon_{xd}+\epsilon_{xp})} \\ K_{XP,X} = [P]\frac{N}{P}e^{\beta(\Delta\epsilon_{pd}+\epsilon_{xp})} \\ K_{SX,S} = [X]\frac{N}{X}e^{\beta\Delta\epsilon_{xd}} \end{cases} \quad [60]$$

We can note that $[X] = \frac{X}{N_A V_{cell}}$ and $[P] = \frac{P}{N_A V_{cell}}$. This then simplifies to equation Eq. [61], which give the expression of this dissociation constants in both the kinetic and thermodynamic viewpoints as,

$$\begin{cases} K_{SP,S} = \frac{k_{PS}}{k_{SP}} = C_N e^{\beta\Delta\epsilon_{pd}} \\ K_{XP,P} = \frac{k_{XP,P}}{k_{XP,P}} = C_N e^{\beta(\Delta\epsilon_{xd}+\epsilon_{xp})} \\ K_{XP,X} = \frac{k_{XP,X}}{k_{XP,X}} = C_N e^{\beta(\Delta\epsilon_{pd}+\epsilon_{xp})} \\ K_{SX,S} = \frac{k_{XS}}{k_{SX}} = C_N e^{\beta\Delta\epsilon_{xd}}, \end{cases} \tag{61}$$

where $C_N = \frac{N}{N_A V_{cell}}$ is the molar concentration of empty sites in the cell.

Let us express the probability of the polymerase being bound to the DNA. First, we may write $p_P = \frac{\rho_P}{\rho_P+\rho_X+\rho_{XP}+\rho_X}$ and $p_{XP} = \frac{\rho_{XP}}{\rho_P+\rho_X+\rho_{XP}+\rho_X}$. Then, we may write,

$$p_{bound} = p_P + p_{XP} = \frac{1+\frac{\rho_{XP}}{\rho_P}}{1+\frac{\rho_{XP}}{\rho_P}+\frac{\rho_X}{\rho_P}+\frac{\rho_S}{\rho_P}} = \frac{1+\frac{[X]}{K_{XP,P}}}{1+\frac{[X]}{K_{XP,P}}+\frac{[X]K_{SP,S}}{[P]K_{SX,S}}+\frac{K_{SP,S}}{[P]}}, \text{ and,}$$

$$p_{bound} = \frac{1+\frac{[X]}{K_{XP,P}}}{1+\frac{K_{SP,S}}{[P]}+[X](\frac{1}{K_{XP,P}}+\frac{K_{SP,S}}{[P]K_{SX,S}})}. \tag{62}$$

We can express this probability in terms of kinetic rate constants and concentrations as,

$$p_{bound} = \frac{1+\frac{[X]k_{XP,P}}{k_{XP,P}}}{1+\frac{k_{PS}}{k_{SP}[P]}+[X](\frac{k_{XP,P}}{k_{XP,P}}+\frac{k_{PS}k_{XS}}{[P]k_{SP}k_{SX}})}. \tag{63}$$

Alternatively, we can also write this probability in term of energies and number of sites as:

$$p_{bound} = \frac{1+Xe^{-\beta(\Delta\epsilon_{xd}+\epsilon_{xp})}}{1+\frac{e^{\beta\Delta\epsilon_{pd}}}{P}+Xe^{-\beta\Delta\epsilon_{xd}}(e^{\beta\epsilon_{xp}}+\frac{e^{\beta\Delta\epsilon_{pd}}}{P})}. \tag{64}$$

We note that $X = \frac{[X]}{C_N}$ and $P = \frac{[P]}{C_N}$. These two expressions are equivalent.

***G.4. Detailed balance is implied by $\gamma = 1$ and steady-state.*** To give concreteness to the general cycle condition we discussed in §C, we return to illustrate this result using the specific parameters of the square graph and a different, perhaps more transparent, algebraic tact.

Why is detailed balance—as expressed in Equation Eq. [57]—equivalent to having a graph at steady state (where the Matrix Tree Theorem applies) and enforcing the cycle condition that the ratio of products of rate constants $\gamma$ is unity? In the square graph, this cycle condition of unity is

$$\gamma \equiv \frac{\gamma_+}{\gamma_-} = \frac{k_{SX}k_{X,XP}k_{XP,P}k_{PS}[X][P]}{k_{XS}k_{XP,X}k_{P,XP}k_{SP}[X][P]} = \frac{k_{SX}k_{X,XP}k_{XP,P}k_{PS}}{k_{XS}k_{XP,X}k_{P,XP}k_{SP}} := 1. \tag{65}$$

First, define $\gamma_+ \equiv k_{SX}k_{X,XP}k_{XP,P}k_{PS}[X][P]$ and $\gamma_- \equiv k_{XS}k_{XP,X}k_{P,XP}k_{SP}[X][P]$, respectively, as the products of rate constant in the + (clockwise) and - (counterclockwise) directions.

We will now prove that at steady state, we can write:

$$\begin{cases} \rho_S k_{SX}[X] - \gamma_+ = \rho_X k_{XS} - \gamma_- \\ \rho_X k_{X,XP}[P] - \gamma_+ = \rho_{XP}k_{XP,P} - \gamma_- \\ \rho_{XP}k_{XP,P} - \gamma_+ = \rho_P k_{P,XP}[X] - \gamma_- \\ \rho_P k_{PS} - \gamma_+ = \rho_S k_{SP}[P] - \gamma_-. \end{cases} \tag{66}$$

This Eq. 66 suffices to show that when $\gamma_+ = \gamma_-$—which guarantees $\gamma = 1$, the cycle condition that ensures equilibrium—the gamma terms cancel, and we recover the equations Eq. [57] that define detailed balance.

To demonstrate the system of equations Eq. [66], we invoke the Matrix Tree Theorem. To illustrate the proof, we discuss just the first equation; the rest follow analogously. Specifically, we can write the statistical weights for the states $X$ and $S$ by applying the Matrix Tree Theorem, seeing that

$$\begin{cases} \rho_S = [X]k_{XS}k_{XP,X}k_{P,XP} + k_{XS}k_{XP,X}k_{PS} + k_{XS}k_{XP,P}k_{PS} + k_{X,XP}k_{XP,P}k_{PS}[P] \\ \rho_X = [X]^2 k_{XP,X}k_{SX}k_{P,XP} + [X]k_{XP,X}k_{SX}k_{PS} + [X]k_{XP,P}k_{SX}k_{PS} + [X]k_{XP,X}k_{SP}k_{P,XP}[P] \end{cases} \tag{67}$$

Then, we multiply by the appropriate rate constants:

$$\begin{cases} \rho_S k_{SX}[X] = k_{XS}k_{XP,X}[X]k_{P,XP}k_{SX}[X] + k_{XS}k_{XP,X}k_{PS}k_{SX}[X] + k_{XS}k_{XP,P}k_{PS}k_{SX}[X] + k_{X,XP}k_{XP,P}k_{PS}[P]k_{SX}[X] \\ \rho_X k_{XS} = k_{XS}k_{XP,X}[X]k_{P,XP}k_{SX}[X] + k_{XS}k_{XP,X}k_{PS}k_{SX}[X] + k_{XS}k_{XP,P}k_{PS}k_{SX}[X] + [X]k_{XP,X}k_{SP}k_{P,XP}[P]k_{XS}. \end{cases} \tag{68}$$

In red, we recognize $\gamma_+$ and in orange $\gamma_-$; the rest of the expressions in blue are equal; and we recover the first equation of Eq. [66], as desired.

***G.5. The cycle condition implies that changing transcription factor or polymerase concentrations does not affect the extent of disequilibrium in the square graph.*** Note that Eq. [65] demonstrates that because $[X]$ and $[P]$ appear in both the products of rates in the clockwise and counterclockwise directions, their influence on the value of $\gamma$ cancels out. This means that adjusting the concentration of transcription factor or polymerase maintains the extent of disequilibrium or equilibrium exhibited by the system.

**H. Driving different arrows in the square graph can still yield a ratio of quadratic polynomials.** Throughout this article, we study the response observable relative to the concentration of transcription factor $[X]$, tuning the edges in green in our square graph as visualized in Figure 1 of the main text. However, depending on the observable and the graph's architecture, the parameter controlling the observable could be different than this transcription factor. For instance, in different biological settings, two rate constants could be adjusted simultaneously by the same scalar control parameter if they are driven by the concentration of a different external (like $ATP$) or internal (like the polymerase $P$) molecule governing the system. Therefore, we can ask: for what classes of control parameter will the observable $\langle r \rangle$ exhibit the same functional form of a ratio of quadratic polynomials?

The Matrix Tree Theorem gives a precise structural answer to this question: when the graph has at least one rooted spanning tree with each of zero, one, and two edges that depend on the control parameter, the observable will inherit such a familiar quadratic dependence. This is a broad class of graphs. We now show some of the diversity of these graphs, whose response shapes and sensitivity bounds are necessarily mathematically identical to those we establish in the first half of the paper, by giving a few concrete examples of related graphs.

Figure S5A illustrates various graphs whose responses are mappable to that of our original square graph (itself illustrated in S5A(i)). The response's form is unchanged when we create a new graph by vertically reflecting the original graph (as in Fig. S5A(ii)), or merely rotating it (not displayed).

Another structurally-distinct but mathematically-equivalent type of graph is shown in Fig. S5A(iii) (also representing any other graph with two controlled edges that may be mapped by reflection or rotation onto the indicated red edges in Fig. S5A(iii)). To understand why this graph has the same quadratic dependence, we can refer to the spanning trees of the square graph using our original rate labels; these spanning trees include $k_{SX}[X]k_{XP,P}k_{X,XP}$ and $k_{SX}[X]k_{XP,P}k_{PS}$, which are both proportional to $k_{SX}k_{XP,P}$, namely both transitions in red imagined to be controlled by the common control variable in Fig. S5A(iii).

Figure S5A(iv) gives another graph where the red indicated arrows both participate in a common spanning tree, assuring the same quadratic dependence of interest. To see this fact, take the two indicated edges and add either the edge $k_{XP,P}$ or the edge $k_{XP,X}$; the results are both valid spanning trees rooted in $S$. Rotating this set of edges also generates three other equivalent graphs with the same behavior (not shown). (One minor difference between the observable produced by this type of graph is that when $[X] \to \infty$, the limit of this graph's observable is now constrained to 1, since the leading order spanning trees in the control parameter are rooted in the same node.)

Last, Fig. S5A(v) acknowledges that many other graphs with a larger set of nodes than four can exhibit the same quadratic form. As just one example, when there are only two controlled (red) transitions localized among some states in a suitable subgraph, all spanning trees of the larger graph can inherit the structural requirements imposed by the subgraph.

Of course, many graphs will not necessarily exhibit this quadratic dependence. Fig. S5B depicts examples of graphs whose outputs will instead display a response behavior mathematically evocative of detailed balance, a ratio of linear polynomials. We can see this contrasting behavior by recalling that a valid spanning tree cannot have more than one outgoing edge per node, nor can it form a complete cycle, meaning that the illustrated graphs will give spanning trees with at most one edge dependent on the control parameter.

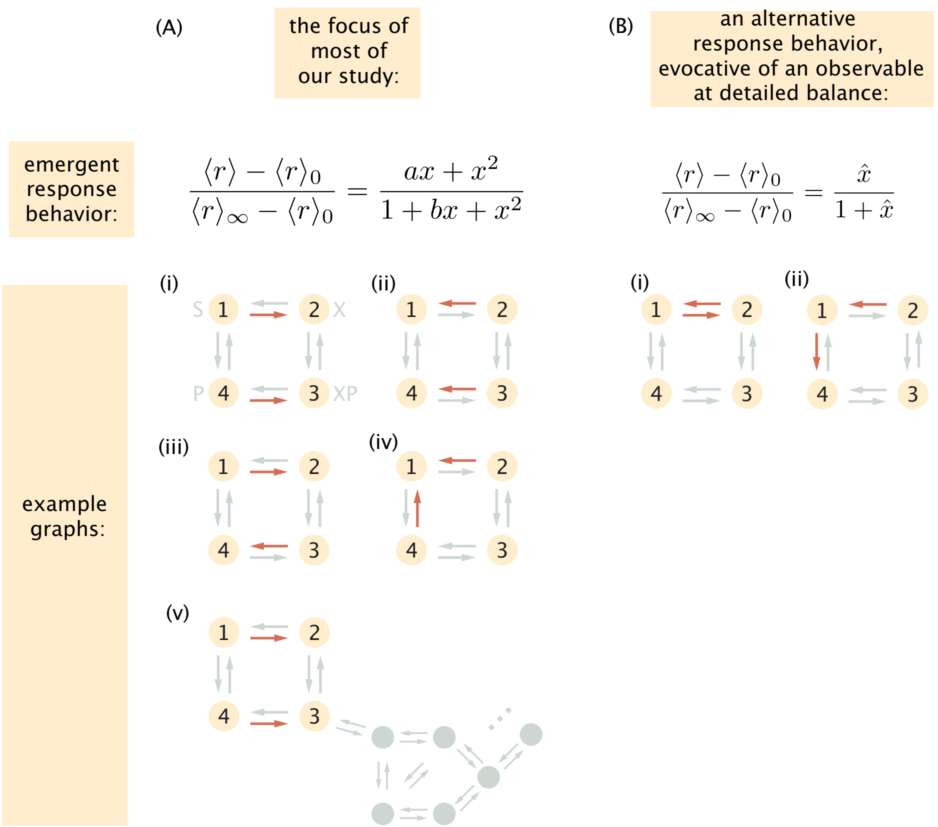

**Fig. S5.** Examples of alternative graph architectures that display (A) the same ratio-of-quadratic-polynomial dependence of the observable (and hence simplified two-parameter emergent shape behavior) in the control parameter, or (B) an observable behavior that evokes a detailed-balance response instead. The red arrows represent transitions whose rates are simultaneously scaled by the control parameter (such as a given transcription factor's concentration).

### I. Any averaged observable $\langle r\rangle$ has zero, one, two, or three inflection points, with varying monotonicity.

***I.1. Descartes' rule of signs on second-derivative-polynomial with $(a, b)$ reveals precise restrictions on numbers of inflections.*** Descartes' rule of signs states that a polynomial $a_0 + a_1 x + a_2 x^2 + \cdots + a_n x^n$ with real coefficients $\{a_i\}$ has at most as many positive roots $P$ as the number of changes in sign $S$ in the sequence $a_0, a_1, \ldots, a_n$ (ignoring coefficients that are zero). Further, this count of the coefficients' sign changes $S$ and the number of positive roots $P$ differ by an even number (24).

Combined with the convenience of the reduced $(a, b)$ shape parameterization, this rule gives transparent and straightforward information about how many inflection points the observable $\langle r\rangle$ may exhibit with respect to the (log) control variable. These inflection points satisfy $\frac{d^2\langle r\rangle}{d(\ln x)^2} = 0$. Since the (changes in) concavity are unchanged by scaling or shifting the function, we can evaluate this equation with respect to the normalized response in terms of the two $(a, b)$ parameters—as in Eq. [28]—instead of the six parameters of the raw quadratic response. Computing the derivative gives

$$\frac{d^2\langle\tilde{r}\rangle}{d^2\ln x} = \frac{x\big(a\big(-b\big(x^3+x\big)\big)+x^4-6x^2+1\big)+x(x(b(x(b-x)+3)-4x)+4)\big)}{(x(b+x)+1)^3}, \qquad [69]$$

where $\langle\tilde{r}\rangle \equiv \frac{\langle r\rangle - \langle r\rangle_0}{\langle r\rangle_\infty - \langle r\rangle_0}$.

This vanishes when the polynomial in the numerator vanishes; so we focus on

$$q(x) \equiv (a-b)x^4 + \left(-ab + b^2 - 4\right)x^3 + (3b - 6a)x^2 + (4 - ab)x + a. \qquad [70]$$

Recalling that $b$ is strictly positive, consider the possible changes in sign in this sequence of coefficients, rewritten suggestively as

$$\{a, 4 - ab, 3(-(a-b) - a), -b(a-b) - 4, a - b\}.$$

These coefficients' signs are constrained differently depending on when $a$ is respectively positive, negative, or zero:

- $a < 0$**:** When all coefficients are nonzero, the signs are $\{\ominus, \oplus, \oplus, \ominus \text{ OR } \oplus, \ominus\}$. This means the sign sequence is either $\{\ominus, \oplus, \oplus, \ominus, \ominus\}$ (giving $S = 2$ sign changes) or $\{\ominus, \oplus, \oplus, \oplus, \ominus\}$ (still giving $S = 2$ sign changes). (While some of these coefficients can go to zero at certain $(a, b)$, shortening the sign sequence, these happen to leave the number of sign changes unchanged from $S = 2$.) Hence when $a < 0$ there are exactly $\boxed{\text{zero or two}}$ (positive) inflection points: in other words, every nontrivial input-output curve with $a < 0$ has two inflection points.

- $a = 0$**:** Now the signs (of nonzero coefficients) are $\{\oplus, \oplus, \oplus \text{ OR } \ominus, \ominus\}$. Observe that there is exactly $S = 1$ sign change. (This is unchanged even if the third coefficient vanishes). So input-output curves with $a = 0$ must have exactly $\boxed{\text{one}}$ inflection point (they are "equilibrium-like").
- $a > 0$**:** Here the sign of $a - b$ critically affects how many positive roots exist:
  - If $a > b$, the signs are $\{\oplus, \oplus \text{ OR } \ominus, \ominus, \ominus, \oplus\}$; hence $S = 2$ sign changes permit exactly $\boxed{\text{zero or two}}$ positive inflection points.
  - If $a < b$, the signs are $\{\oplus, \ominus \text{ OR } \oplus, \ominus \text{ OR } \oplus, \ominus \text{ OR } \oplus, \ominus\}$. Hence there are up to $S = 3$ sign changes, permitting $\boxed{\text{one or three}}$ positive inflection points.

In general, this analysis has often benefited from the fact that if the signs of two or more coefficients are fixed at key positions in the coefficient sequence, then ambiguity in the signs of the coefficients in between has no effect on the number of possible changes of sign. For instance, the fact that the zeroth and fifth coefficients are respectively positive $\oplus$ and negative $\ominus$ in the last $0 < a < b$ case just examined immediately ensures that $S < 4$, so there are not four inflection points possible here (despite initial impressions from the fact that the underlying polynomial is a quartic).

The general conclusions we have reached from this elementary application of Descartes' rules are wholly consistent with a more precise, and algebraically-elaborate, inspection of the inflection points in the $(a, b)$ space, as now follows. (We give both analyses because the former may add some transparency.)

***I.2. Monotonicity of response via $(a, b)$ parameterization.*** Here, we find the conditions on the emergent shape parameters $(a, b)$ participating in the normalized response of Eq. [28] that assure nonmonotonicity. Since the logarithm is itself a monotonic transformation, the (non)monotonicity of responses remains unchanged whether we regard them with respect to the input variable on a linear scale or logarithmic scale. So for algebraic convenience, we inspect the first derivative of the response Eq. [28] with respect to the input on a linear scale, finding

$$\frac{d\langle r\rangle}{dx} = (\langle r\rangle_\infty - \langle r\rangle_0)\,\frac{(b-a)x^2 + 2x + a}{(x(b+x)+1)^2}. \qquad [71]$$

The response $\langle r\rangle(x)$ is nonmonotonic if this derivative changes sign. Since $x$ must be positive on physical grounds (as when it represents a concentration), we further demand that the derivative change sign for some $x > 0$. The polynomial in the derivative's numerator, $p(x) \equiv (b-a)x^2 + 2x + a$, behaves according to its discriminant

$$\Delta \equiv 4(1 - a(b-a)), \qquad [72]$$

and the roots

$$x_\pm = \pm\sqrt{\frac{a^2 - ab + 1}{(a-b)^2}} + \frac{1}{a-b} = \frac{1}{a-b}\left(1 \pm \sqrt{1 + a(a-b)}\right). \qquad [73]$$

This polynomial has real solutions when the discriminant is nonnegative, $\Delta \geq 0$, namely, $1 - a(b-a) \geq 0$. Recalling that $b > 0$ by construction, one way for this to happen is when $a < 0$. Another way for the discriminant to be positive is when $a > 0$ while still ensuring that $a(b-a) < 1$, or equivalently $0 < b < a + \frac{1}{a}$.

The requirement that at least one root be positive further refines these conditions on $(a, b)$. We proceed by inspecting the positivity of roots under each possible condition that ensures they are real:

- $a < 0$**:** Only the root $x_- = \frac{1}{a-b} - \frac{1}{a-b}\sqrt{1 + a(a-b)}$ could be positive, since sign$\left(\frac{1}{a-b}\right) = \ominus$. In this case, we still need to verify that this root $x_- > 0$; this is true when $1 - \sqrt{1 + a(a-b)} < 0$. Happily this must be true, since $a(a-b)$ is a positive number, meaning the term in the square root is greater than one and so the square root is also greater than one. Hence, the case of $a < 0$ automatically ensures there is a real and positive solution to the inflection point changing sign (and thus nonmonotonicity).
- $0 < b < a + \frac{1}{a}$**, but** $b > a > 0$: Since $a$ is now positive but still smaller than $b$, we still have sign$\left(\frac{1}{a-b}\right) = \ominus$, still suggesting $x_+$ cannot be positive. However, in this case, we further see that $1 + a(a-b) < 1$, so the other root $x_-$ is also negative. Therefore, this condition does not guarantee nonmonotonicity.
- $a > b > 0$: Now, sign$\left(\frac{1}{a-b}\right) = \oplus$, and the term under the square root in the discriminant is greater than one. This means that only the root $x_+$ can be positive, which is automatically the case. Hence $a > b$ suffices to ensure nonmonotonicity.

(We also note that the discriminant cannot vanish and also produce a positive $x > 0$, ensuring these are the only conditions enabling nonmonotonicity.) Altogether, we summarize the necessary and sufficient conditions for nonmonotonicity, where $a, b$ are defined, as

$$\boxed{\text{nonmonotonicity} \equiv \begin{cases} a > 0 \text{ and } a > b, \text{or} \\ a < 0 \text{ and } b > 0 \end{cases}}. \qquad [74]$$

When we return shortly to consider the number of inflection points possible for a response curve, we will see that these conditions for nonmonotonicity only intersect the conditions for having two inflection points, establishing that singly or triply inflected responses must be monotonic.

***I.3. Bounds on the absolute magnitudes of response extrema.*** If a response is monotonic, then for any $[X]$, it must always be bounded above and below by the leakiness and saturation values $\langle r\rangle_0$ or $\langle r\rangle_\infty$. So finding an upper or lower bound on the response only becomes more subtle and interesting in the case of nonmonotonic responses.

To make progress, we translate the nonmonotonicity conditions Eq. [74] more concretely in term of the values $\frac{B}{E}$, $\langle r\rangle_0$ and $\langle r\rangle_\infty$. This process shows that a response is nonmonotonic if any of the following conditions are true:

$$\begin{cases} \text{condition 1:} & \langle r\rangle_\infty > \langle r\rangle_0 > \frac{B}{E}, \text{ or} \\ \text{condition 2:} & \frac{B}{E} > \langle r\rangle_\infty > \langle r\rangle_0, \text{ or} \\ \text{condition 3:} & \frac{B}{E} > \langle r\rangle_0 > \langle r\rangle_\infty, \text{ or} \\ \text{condition 4:} & \langle r\rangle_0 > \langle r\rangle_\infty > \frac{B}{E}. \end{cases} \quad [75]$$

In general, this reasoning establishes that for any type of response (nonmonotonic or monotonic),

$$\min\left\{\langle r\rangle_0, \langle r\rangle_\infty, \frac{B}{E}\right\} \le \langle r\rangle \le \max\left\{\langle r\rangle_0, \langle r\rangle_\infty, \frac{B}{E}\right\}. \quad [76]$$

Returning to the individual conditions for nonomonotonicity, we see they each give separate bounds for the extremal values of the observable:

$$\begin{cases} \text{condition 1:} & \frac{B}{E} \le \langle r\rangle \le \langle r\rangle_\infty \\ \text{condition 2:} & \langle r\rangle_0 \le \langle r\rangle \le \frac{B}{E} \\ \text{condition 3:} & \langle r\rangle_\infty \le \langle r\rangle \le \frac{B}{E} \\ \text{condition 4:} & \frac{B}{E} \le \langle r\rangle \le \langle r\rangle_0. \end{cases} \quad [77]$$

Therefore the quantity $\frac{B}{E}$ bounds the extremum of any nonmonotonic response function.

The upper and lower bounds on any observable, Eq. [76], follow from a simple elementary result bounding ratios of sums. We quickly digress to establish the elementary result:

*Simple bound on ratios of non-negative sums.* For nonnegative $a_i, b_i$,

$$\min_i\left(\frac{a_i}{b_i}\right) \le \frac{\sum_{i=1}^{N} a_i}{\sum_{i=1}^{N} b_i} \le \max_i\left(\frac{a_i}{b_i}\right). \quad [78]$$

---

Consider the lower bound/left inequality. By definition, we know

$$\min_i\left(\frac{a_i}{b_i}\right) \le \frac{a_j}{b_j}, \text{ for all } j \in [1, N] \quad [79]$$

Multiplying by $b_j$ on both sides,

$$\min_i\left(\frac{a_i}{b_i}\right) b_j \le a_j, \text{ for all } j \in [1, N] \quad [80]$$

and summing over all $j$ gives

$$\min_i\left(\frac{a_i}{b_i}\right) \times \sum_{j=1}^{N} b_j \le \sum_{j=1}^{N} a_j. \quad [81]$$

Hence indeed, $\min_i\left(\frac{a_i}{b_i}\right) \le \frac{\sum_{j=1}^{N} a_j}{\sum_{j=1}^{N} b_j}$ as desired. The right (upper bound) inequality follows identically.

Returning to the ratio of polynomials form $\langle r\rangle = \frac{A+B[X]+C[X]^2}{D+E[X]+F[X]^2}$, this means that

$$\min\left\{\frac{A}{D} = \langle r\rangle_0, \frac{B}{E}, \frac{C}{F} = \langle r\rangle_\infty\right\} \le \langle r\rangle \le \max\left\{\frac{A}{D} = \langle r\rangle_0, \frac{B}{E}, \frac{C}{F} = \langle r\rangle_\infty\right\}, \quad [82]$$

which supports the claim of Eq. [76] and Eq. [77].

***I.4. Number of inflection points via the $(a, b)$ parameterization.*** Now we study the number of inflection points of the observable with respect to the control parameter on a logarithmic scale. To do this, we study the polynomial that appears in the numerator of the second derivative with respect to log control variable, Eq. [69],

$$q(x) \equiv x^4(a-b) + x^3\left(-ab + b^2 - 4\right) + x^2(3b - 6a) + x(4 - ab) + a. \quad [83]$$

In what follows, we examine how many roots of this polynomial can simultaneously be real and positive. As a preview of this logic, we do this by solving for each of the roots of the quartic; finding independent conditions on the parameters $a, b$ that ensures each of these roots would be positive and real; then consider all the possible logical unions of these conditions, testing whether zero up to four inflections are simultaneously defined. We largely perform this tedious procedure using the symbolic capabilities of *Mathematica*—see our Github code repository for more details—and do not suggest that the intermediate conditions on individual roots are themselves enlightening or transparent. Yet their collective implications are meaningful and so we summarize them below.

The polynomial Eq. [83] can have up to four roots; denote them $(x_1, x_2, x_3, x_4)$. These roots have a closed-form solution given by the famously grotesque quartic formula or returnable by *Mathematica*. Asking each of them to be positive and real gives individual conditions on $(a, b)$; denote these conditions $C_1, C_2, C_3, C_4$, where $C_i$ is the set of conditions where root $x_i$ is real and positive. Then the condition of finding zero inflection points is the setting where none of $C_1, C_2, C_3$, or $C_4$ are true; the condition of finding one inflection point is where exactly one of them is true; and so on.

This analysis reveals two trivial cases. First, when there are no inflection points, the response transpires to be constant everywhere for all positive $x$, namely $\langle r\rangle = \langle r\rangle_0 = \langle r\rangle_\infty$. Second, we find that since not all of $C_1, C_2, C_3, C_4$ can be simultaneously true, it is impossible for the function to have four inflection points.

In contrast, it is readily possible to reach one, two, or three inflection points under specific parametric conditions. The borders between these conditions have somewhat complicated structure, particularly between the one and three inflection point cases. To assist us in expressing them as concisely as feasible, define the polynomial

$$H_a(b) \equiv -1024 - 1024a^2 + 1024ab + (-64 - 64a^2)b^2 + 64ab^3 + (-28 - a^2)b^4 + ab^5, \quad [84]$$

and in particular define its three real and positive roots when solving it with respect to the shape parameter $b$ given $a$: denote them $b_1(a), b_2(a), b_3(a)$. (These roots turn out to form independent branches of an implicit representation of the border between one and three inflection point regimes, each valid for different restricted values of $a$.) The final ingredient needed to define the borders between logical conditions turns out to be a numerical constant cutoff value of $a$, approximately $a_{\text{lim}} \approx 2.35$ (see Mathematica code on Github and figure S6). Armed with these ingredients, the conditions to reach one, two, and three inflection point curves are expressed as follows, and plotted explicitly in Figure S6.

Output curves are "equilibrium-like," presenting only one inflection point, when

$$\boxed{\textbf{one inflection, monotonic} \equiv (b \le b_1(a) \text{ or } (b_3(a) \ge b \ge b_2(a), a \in [2, a_{\text{lim}}])) \text{ and } a \ge b \text{ or } a = 0}. \quad [85]$$

It transpires that output curves have two inflection points exactly under the same conditions on $a, b$ as we found assured nonmonotonicity in Eq. [74]: namely,

$$\boxed{\textbf{two inflections, nonmonotonic} \equiv \begin{cases} a > 0 \text{ and } a > b, \text{or} \\ a < 0 \text{ and } b > 0 \end{cases}}. \quad [86]$$

(Note that this condition also subsumes the case $a = \pm\infty$, where the observable is also nonmonotonic.)

Output curves show three inflection points if,

$$\boxed{\textbf{three inflections, monotonic} \equiv b > b_1(a) \text{ and } (b > b_3(a) \text{ or } b_2(a) > b, a \in [2, a_{lim}])}. \quad [87]$$

We can summarize the border between one and three inflection point responses by considering the shape of this overall implicit function, $b_{\text{cutoff}}(a)$, defined as

$$b_{\text{cutoff}}(a) = \begin{cases} \max(b_1(a), b_2(a), b_3(a)) \; if \; 2 \le a < a_{lim} \\ b_1(a) \; else \end{cases} \quad [88]$$

We visualize this cutoff function in Fig. S7.

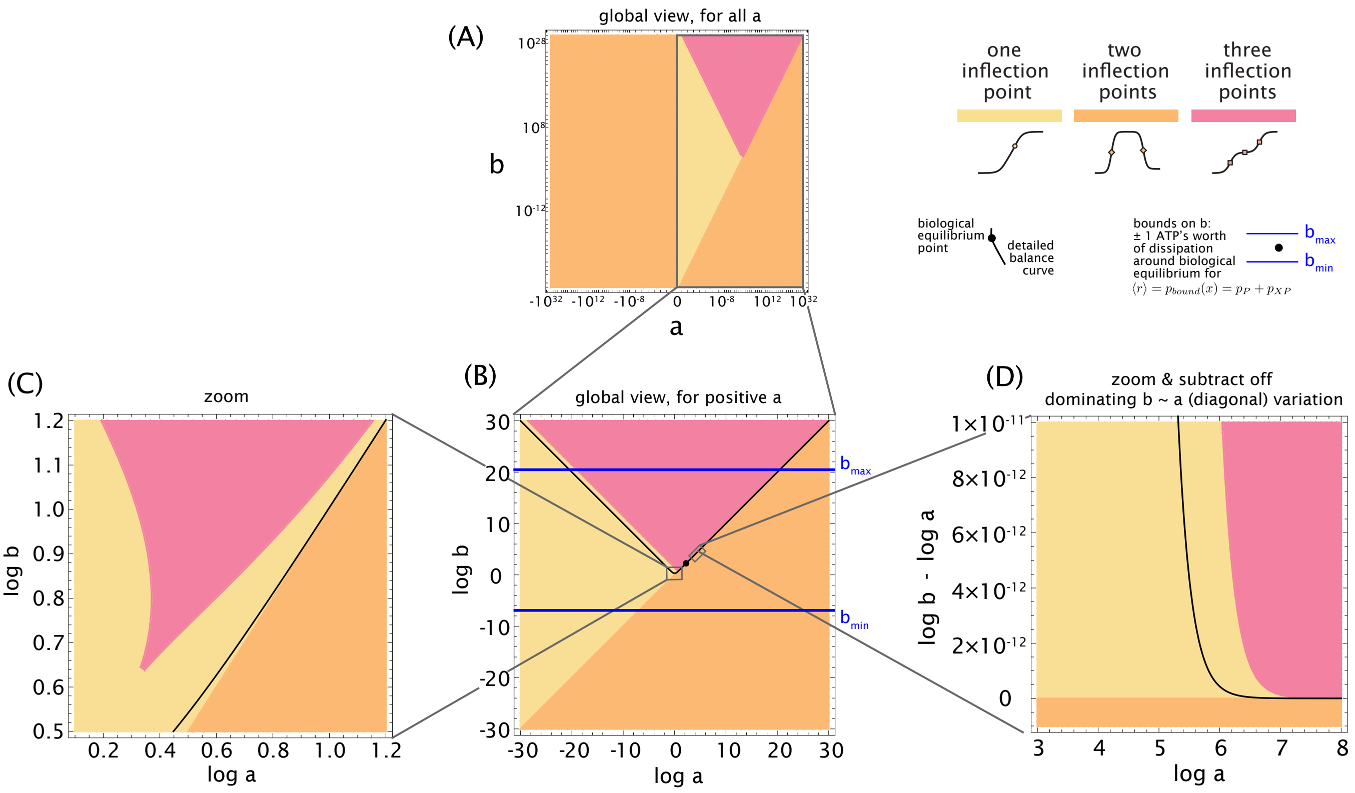

**Fig. S6.** The values of natural parameters $(a, b)$ completely determine the shape of each response curve. Quantitative criteria partition the space into regions with either one inflection (pale yellow), two inflections (orange), or three inflections (pink). The central panels (A) and (B) give global views of $(a, b)$ phase space centered around biological equilibrium, either for both positive and negative $a$ (panel A) or for the subset $a > 0$ (panel B). Blue lines indicate the minimum and maximum values of $b$ reachable by driving any single edge at a time by $\Delta\mu \leq 20k_BT$. When $a < 0$, response curves are always nonmonotonic (with two inflection points). Overall, the two-inflection-point phenotype is the most common in this space (for all $a$; the subspace where $a$ is positive; or in the region where $a > 0; b \in [b_{min}, b_{max}]$). Systems satisfy detailed balance on the black line. The black dot denotes the default equilibrium starting rates reported in Fig. 1A of the main text, or Fig. K.1. At left in (C) is a zoom of the same space near biological equilibrium, validating that the detailed balance curve always lies within the one-inflection thinly-shaped region that bridges the two-inflection point and three-inflection point regions. At right is another zoom of the ribbon region, but where the major diagonal covariation of $b$ with $a$ has been subtracted away (by plotting $\log b - \log a$ versus $a$ instead of $\log b$ versus $\log a$). This visualizes how the detailed balance curve becomes asymptoptically closer to the border with the two-inflection-point regime (lower boundary/orange) versus the (upper boundary/pink) three-inflection-point regime as $a$ grows larger.

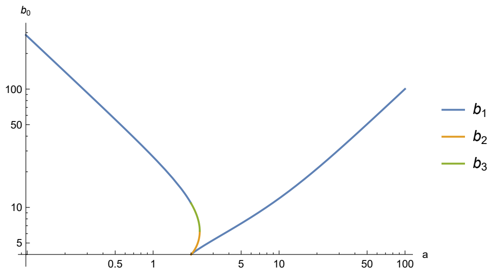

**Fig. S7.** The value of the cutoff $b_{\text{cutoff}}(a)$, defined in Equation Eq. [88] with respect to $a$, delimits the first and third inflection points regimes.

At equilibrium, the collapse of an observable to a ratio of linear polynomials (Eq. [26]) allows us to rewrite the normalized response as

$$\langle r \rangle_{eq} = \langle r \rangle_0 + (\langle r \rangle_\infty - \langle r \rangle_0)\frac{dx}{1 + dx}.$$

The constant $d$ is the same in the numerator and denominator, so that the limit at infinity of the observable is $\langle r \rangle_\infty$. For the detailed balance case, we can identify $\langle r \rangle(x) = \langle r \rangle_{eq}(x) \forall x \in \mathbb{R}^{*+}$. This is equivalent to seeing the polynomial $R(X) = X(d(b - a) - 1) + d - a$ have each of its coefficients vanish. This situation implies that the coefficients are related to one another according to,

$$\begin{cases} d = a \\ b = a + \frac{1}{a}. \end{cases} \quad [89]$$

Note that the detailed balance curve always lies within the one-inflection (pale yellow) region: this region forms is a thin ribbon between the three and two inflection points region along the diagonal $a = b$. The detailed balance curve becomes asymptoptically closer to the border with the two-inflection-point regime (lower boundary/orange) versus the (upper boundary/pink) three-inflection-point regime as $a$ grows larger (see Figure S6).

### J. New bounds on nonequilibrium sensitivity.

***J.1. Motivation of the the definition of the normalized sensitivity.*** Sensitivity—how steeply output changes with input—is one of the most fundamental quantitative traits that energy expenditure can modulate in biological systems, as celebrated by a plethora of famous biological models (e.g. the Goldbeter-Koshland ultrasensitivity mechanism (25), *inter alia*). Nonetheless, network architecture imposes strong constraints on the maximal sensitivities systems can achieve (1), even under arbitrarily large drive. We investigate sensitivity (and bounds thereof) for our setting in this spirit, but strive to use mathematical quantities that align closely with experimental conventions.

One common measure of sensitivity in conversation with experimental measurements and existing performance bounds is simply the (raw) *sharpness* (with respect to an input $x$),

$$\text{sharpness} \equiv \frac{d\langle r\rangle}{d\ln x} \tag{90}$$

$$= x\frac{d\langle r\rangle}{dx}. \tag{91}$$

Reference (9) is an example of a recent study which assesses sensitivity using this sharpness. The convention of considering changes in the raw response output with respect to a logarithmic input is also natural and coherent with the plotting convention of a logarithmic input, as discussed in §E. (If the response were exactly a Hill function with a Hill coefficient $H$, itself a common measure of sensitivity, then this sharpness would reach a maximal value of $H/4$ at the vertical midpoint of the response curve (1).) (When $x$ is viewed as a concentration, we should recall that we render it unitless before taking the logarithm by viewing it as a normalized concentration relative to some reference $[X]_0$, say $[X]_0 \equiv 1$ nanomolar, just as discussed in §E.)

To establish bounds on the sensitivity agnostic to specific parameter values or energetic dissipations, we normalize the raw sharpness, defining as our principal measure of *normalized sensitivity*,

$$\text{normalized sensitivity } s([X]) \equiv \left|\frac{d\langle r\rangle}{d\ln\left({}^{[X]}/_{[X]_0}\right)}\frac{1}{\langle r\rangle_{\max}-\langle r\rangle_{\min}}\right|. \tag{92}$$

where we defined $\langle r\rangle_{\min} \equiv \min\limits_{[X]} \langle r\rangle$ and $\langle r\rangle_{\max} \equiv \max\limits_{[X]} \langle r\rangle$.

This definition of normalized sensitivity is related to the separately-normalized output $\tilde{r} \equiv \frac{\langle r\rangle-\langle r\rangle_0}{\langle r\rangle_\infty-\langle r\rangle_0}$ in ways that vary depending on the curve's shape. We review these relationships in each possible curve shape now. When the response remains monotonic (namely when it has one or three inflection points), the normalized sensitivity is equal to

$$\text{monotonic:}\quad s([X]) = \frac{d\langle r\rangle}{d\ln[X]}\frac{1}{\langle r\rangle_\infty-\langle r\rangle_0} = \frac{d\tilde{r}}{d\ln x}, \tag{93}$$

since $\langle r\rangle_\infty - \langle r\rangle_0$ is the range of variation of the output curve.

When the output is nonmonotonic, if $a < 0$, then the output is first decreasing up to $\langle r\rangle_*$ and then increasing, since $a$ is the value of the slope at zero concentration of the normalized rate. In this regime the maximum of slope is reached at second inflection. Hence, the corresponding range of variation of the rate is $\langle r\rangle_\infty - \langle r\rangle_*$, and the normalized sensitivity assumes the meaning

$$\text{nonmonotonic, } a<0\text{: } s([x]) = \frac{d\langle r\rangle}{d\ln[X]}\frac{1}{\langle r\rangle_\infty-\langle r\rangle_*} = \frac{d\tilde{r}}{d\ln x}\frac{\langle r\rangle_\infty-\langle r\rangle_0}{\langle r\rangle_\infty-\langle r\rangle_*} = \frac{d\tilde{r}}{dx}\frac{1}{1-\tilde{r_*}} = \frac{d\tilde{r}}{dx}\frac{1}{\tilde{r_\infty}-\tilde{r_*}} \tag{94}$$

When the output is nonmonotonic but $\frac{a}{b} < 1$, the response is first increasing up to $\langle r\rangle_*$ and then decreasing to the value $\langle r\rangle_\infty$. The maximum of slope is reached at first inflection and the range of variation of the output values is $\langle r\rangle_* - \langle r\rangle_0$. Therefore the normalized slope becomes:

$$\text{nonmonotonic, } a/b<1\text{: } s([x]) = \frac{d\langle r\rangle}{d\ln[X]}\frac{1}{\langle r\rangle_*-\langle r\rangle_0} = \frac{d\tilde{r}}{d\ln x}\frac{\langle r\rangle_\infty-\langle r\rangle_0}{\langle r\rangle_*-\langle r\rangle_0} = \frac{d\tilde{r}}{d\ln x}\frac{1}{\tilde{r_*}} = \frac{d\tilde{r}}{d\ln x}\frac{1}{\tilde{r_*}-\tilde{r_0}}. \tag{95}$$

***J.2. Connection to other measures of sensitivity and the effective Hill coefficient.*** Here we clarify a few distinct but related notions of sensitivity. First, the *logarithmic sensitivity* of a response, measuring how inputs change a fold-change in response, is the response's logarithmic derivative with respect to its input,

$$\text{log. sensitivity} \equiv \frac{d\ln\langle r\rangle}{d\ln x} \tag{96}$$

$$= \frac{1}{\langle r\rangle}\underbrace{\frac{d\langle r\rangle}{d\ln x}} \tag{97}$$

$$= \frac{x}{\langle r\rangle}\frac{d\langle r\rangle}{dx}. \tag{98}$$

The derivative of the raw response with respect to the log control variable, $\frac{d\langle r\rangle}{d\ln x}$ as emphasized with an underbracket in Eq. [98], is the raw sharpness we focus on throughout our analysis. It differs from logarithmic sensitivity only by a factor $\frac{1}{\langle r\rangle}$, whose own magnitude is bounded.

As discussed superbly and pedagogically by Owen and Horowitz (1), the logarithmic sensitivity is directly related to various notions of effective Hill coefficients. One definition of an effective Hill coefficient $H_{\text{eff}}$ is explicitly proportional to the logarithmic sensitivity at a midpoint of the response (1), as used for example by references (26, 27):

$$H_{\text{eff}} \equiv 2\, \frac{d\ln\langle r\rangle}{d\ln x}\bigg|_{x=x*} = 2\frac{1}{\langle r\rangle(x^{*})}\, \frac{d\langle r\rangle}{d\ln x}\bigg|_{x=x*} \qquad [99]$$

Hence the sharpness or normalized sensitivity we consider thus enjoys a close, though not identical, connection with these other measures of sensitivity such as effective Hill coefficients.

***J.3. Summary of our results; contrast with existing bounds.*** As we report and illustrate in Figure 2 of the main text, we find that the normalized sensitivity is bounded by finite values,

$$\text{1 inflection:} \qquad 0.158045 \leq s\left([X]\right) \leq \frac{1}{2}, \qquad [100]$$

$$\text{2 inflections:} \qquad \frac{1}{4} \leq s\left([X]\right) \leq \frac{1}{2}, \qquad [101]$$

$$\text{3 inflections:} \qquad \frac{1}{8} \leq s\left([X]\right) \leq \frac{1}{4}. \qquad [102]$$

Our main foundation for bounding response sensitivity is a dense numerical sampling of response curves facilitated by our two-dimensional representation of all responses: see Fig. S8. Specifically, we compute the normalized sensitivity on a fine grid of $(a, b)$ values, observing the bounds above; we also symbolically simplify analogous logical conditions using *Mathematica*, finding concordance with these numbers. For instance, the curious number 0.158045 as a lower-bound on singly-inflected responses is reported with six decimals of precision because this was verified by explicit symbolic simplifications in *Mathematica*.

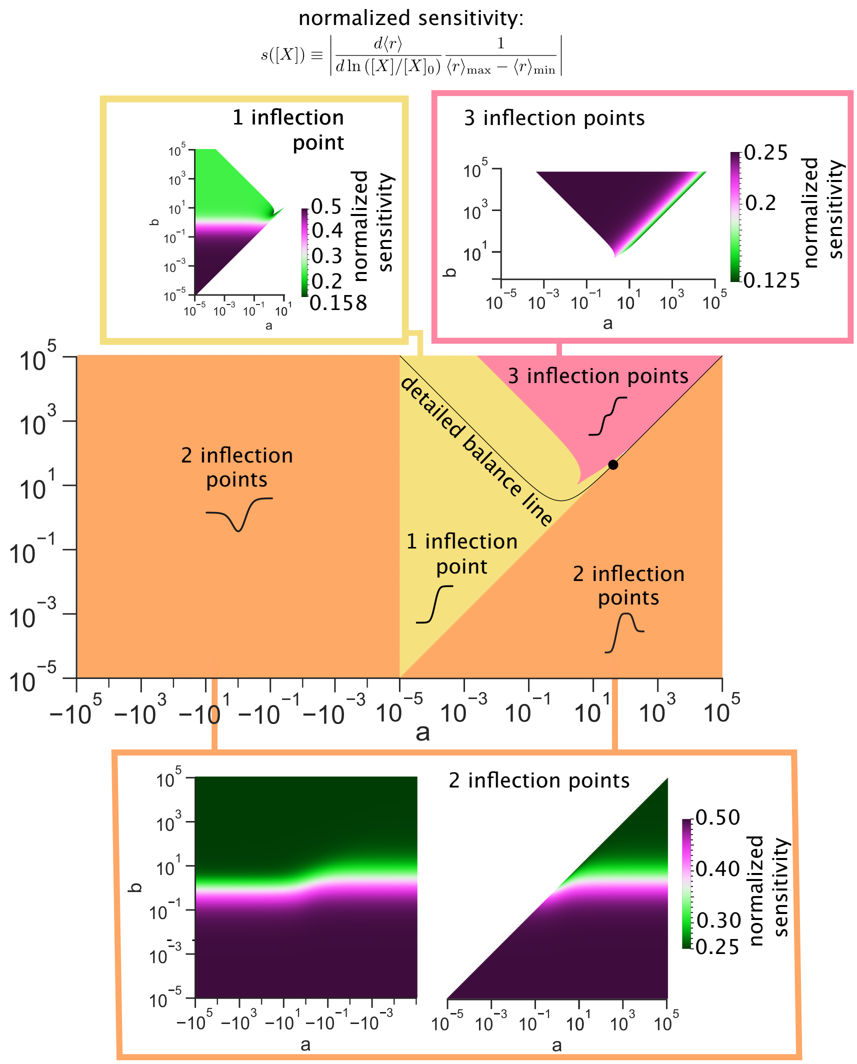

**Fig. S8.** Numerical validation of bounds on normalized maximal sensitivities over all curve phenotypes. Outset heatmaps depict the normalized sensitivities for curves of each region in $(a, b)$ curve shape parameter space. Bounds are visible as the minimum and maximum sensitivities observed in each shape category.

To augment these numerical results, we provide some—albeit incomplete—analytical results; these follow in the next three subsections. First, we establish a looser global analytic upper bound on sensitivity, using a straightforward extension of recently-established upper bound arguments (1) on a related, differently-normalized slope. Second, we establish symbolically a slightly tighter global upper bound for monotonic outputs, that $s([X]) \le \frac{1}{2}$. Last, for triply-inflected curves, we demonstrate symbolically both of our lower and upper bounds, $\frac{1}{8} \le s([X]) \le \frac{1}{4}$.

In conclusion, however, we continue to lack elegant or insightful analytical justifications for all of the lower bounds across regulatory shape phenotypes, or the upper bound on nonmonotonic responses, that we discover in numeric sampling. Interpretably demonstrating these behaviors will be a natural, fruitful subject of analytical work in the future.

***J.4. General upper bound on a related, differently-normalized slope.*** Here we prove a (weaker) upper bound on a different sensitivity, closely connected with the fertile results of Owen & Horowitz (1). We will show that

$$\left| \frac{d\langle r \rangle}{d\ln x} \frac{1}{r_{\max} - r_{\min}} \right| \le \frac{1}{2}, \qquad [103]$$

where we define the (unbracketed) quantities $r_{\min} \equiv \min_{\text{states } i} r_i$ and $r_{\max} \equiv \max_{\text{states } i} r_i$. We will call these quantities "theoretical" extrema because they are the ultimate extrema of observable weights over all microscopic states. Importantly these theoretical extrema are **not** the same as the (bracketed) quantities $\langle r \rangle_{\min} \equiv \min_{[X]} \langle r \rangle$ and $\langle r \rangle_{\max} \equiv \max_{[X]} \langle r \rangle$, the "observed extrema," that our actual normalized sensitivity transacts in. (We will return to contrast the implications of these extrema shortly, after we have established this weaker result.)

To proceed, we invoke a useful result from Owen & Horowitz (1), who establish that

$$\left| \frac{d \ln \langle O_1 \rangle / \langle O_2 \rangle}{d \ln x} \right| \leq m, \tag{104}$$

where $\langle O_1 \rangle \equiv \sum_{\text{states } i} O_{1i} p_i$ and $\langle O_2 \rangle \equiv \sum_{\text{states } i} O_{2i} p_i$ are observables defined by (positive) coefficients $O_{1i}, O_{2i}$; and $m$ is the "size of the support," namely the number of states possessing at least one outgoing transition that is scaled by the control variable. Here in our square graph, $m = 2$.

Next, to invoke the normalization by extrema we desire, we choose the observable weights $O_{1i} \equiv r_i - r_{\text{min}}$ and $O_{2i} \equiv r_{\text{max}} - r_i$. These weights are clearly nonnegative, and so Eq. [104] applies. As a consequence, observe that $\langle O_1 \rangle = \sum_i (r_i - r_{\text{min}}) p_i = \sum_i r_i p_i - r_{\text{min}} \sum_i p_i = \langle r \rangle - r_{\text{min}}$, and similarly $\langle O_2 \rangle = r_{\text{max}} - \langle r \rangle$. The bound Eq. 104 then becomes,

$$\frac{d \ln(\langle r \rangle - r_{\text{min}})}{d \ln x} - \frac{d \ln(r_{\text{max}} - \langle r \rangle)}{d \ln x} \leq m \tag{105}$$

$$\rightarrow \frac{1}{\langle r \rangle - r_{\text{min}}} \frac{d\langle r \rangle}{d \ln x} - \frac{1}{r_{\text{max}} - \langle r \rangle} \frac{-d\langle r \rangle}{d \ln x} \leq m \tag{106}$$

$$\rightarrow \frac{d\langle r \rangle}{d \ln x} \left( \frac{1}{\langle r \rangle - r_{\text{min}}} + \frac{1}{r_{\text{max}} - \langle r \rangle} \right) \leq m \tag{107}$$

$$\rightarrow \frac{d\langle r \rangle}{d \ln x} (r_{\text{max}} - r_{\text{min}}) \leq m(\langle r \rangle - r_{\text{min}})(r_{\text{max}} - \langle r \rangle). \tag{108}$$

On the right side, note that $\langle r \rangle - r_{\text{min}}$ can be at most halfway between the minimum and maximum values of $r$, namely $(\langle r \rangle - r_{\text{min}}) \leq \frac{r_{\text{max}} - r_{\text{min}}}{2}$. The same is true for $r_{\text{max}} - \langle r \rangle$, e.g. $(r_{\text{max}} - \langle r \rangle) \leq \frac{r_{\text{max}} - r_{\text{min}}}{2}$. So their product in the right-hand side is at most $\frac{(r_{\text{max}} - r_{\text{min}})^2}{4}$. This gives

$$\rightarrow \frac{d\langle r \rangle}{d \ln x} (r_{\text{max}} - r_{\text{min}}) \leq m \frac{(r_{\text{max}} - r_{\text{min}})^2}{4}, \tag{109}$$

or

$$\boxed{\frac{d\langle r \rangle}{d \ln x} \leq \frac{m}{4} (r_{\text{max}} - r_{\text{min}})}. \tag{110}$$

Substituting $m = 2$, as appropriate for the square graph, yields the desired result Eq. [103]. □

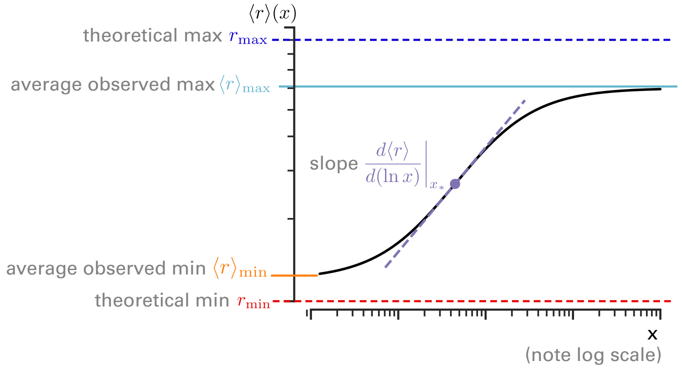

**Fig. S9.** Comparison of response extrema entering different bounds. In general, the observed minima of responses give tighter bounds on a particular response curve than theoretical minima of responses over microstates.

Now we contrast this result Eq. [103], defined in terms of the theoretical extrema $r_{\text{min}}, r_{\text{max}}$ over microstates, with our observed bounds on sensitivity defined in terms of the average *observed* extrema, $\langle r \rangle_{\text{min}}, \langle r \rangle_{\text{max}}$. In general, the theoretical resposne extrema themselves more conservatively bound the response than the observed response extrema. That is, in general the extrema of the *average* observable response curve over all $[X]$ are usually more restricted than the most extreme potencies over microstates (namely, $r_{\text{max}} \equiv \max_i \{r_i\} \geq \langle r \rangle_{\text{max}}$ and $r_{\text{min}} \equiv \min_i \{r_i\} \leq \langle r \rangle_{\text{min}}$. This property is visualized in Fig. S9.

Hence, for a *generic* response curve, the bounds Eq. [102] we discover and focus on in the main text of the paper are in fact tighter than that reported by Eq. 103.

One reason we study that normalized sensitivity $s([X]) \equiv \left| \frac{d\langle r \rangle}{d \ln x} \frac{1}{\langle r \rangle_{\text{max}} - \langle r \rangle_{\text{min}}} \right|$ is to try to connect more directly with measurements of biological curves that do not necessarily represent architectural optima. Indeed, for instance, the observable weights (e.g. here, microscopic transcription rates) $r_i$ of every microstate $i$ are sometimes less easily known or convenient to measure (and so too their extremal values $r_{\max} \equiv \max_i\{r_i\}$ and $r_{\min} \equiv \min_i\{r_i\}$) than the average observable itself. Conversely, the observed extrema $\langle r\rangle_{\max}, \langle r\rangle_{\min}$ can often be directly "read off" from an averaged observable curve $\langle r\rangle([X])$.

We remark that when one is instead asking questions about optimal sensitivities realizable over all architectures, it is is plausible that these two styles of bound become equivalently informative. Specifically, as Jordon Horowitz suggests in personal communication, it is plausible that the response architectures which in fact saturate the bounds are also exactly those where $\langle r\rangle_{\min} \to r_{\min}$ and $\langle r\rangle_{\max} \to r_{\max}$.

***J.5. General upper bound on our normalized sensitivity.*** Now, returning to our normalized slope $s([X]) = \left|\dfrac{d\langle r\rangle}{d\ln x}\dfrac{1}{\langle r\rangle_{max} - \langle r\rangle_{min}}\right|$ that is defined in terms of the observed (not theoretical) extrema, we show $s([X]) \leq \frac{1}{2}$ for all outputs.

For monotonic cases, we use the main result stated earlier from Reference (1), Eq. [104]. For simplicity, we note

$$\hat{r} = \frac{\langle r\rangle - \langle r\rangle_{min}}{\langle r\rangle_{max} - \langle r\rangle_{min}}, \qquad [111]$$

where $\langle r\rangle_{min/max}$ is the minimum (maximum) value of the average observable $\langle r\rangle$ over all positive values of concentration $[X]$. Both $\langle O_1\rangle = \hat{r}$ and $\langle O_2\rangle = 1 - \hat{r}$ are rational functions with positive coefficients. Now, using the general expression of the output rate Eq. [24], we re-express the form of $\hat{r}$ as,

$$\hat{r} = \frac{(A - \langle r\rangle_{min}D) + (B - \langle r\rangle_{min}E)[X] + (C - \langle r\rangle_{min}F)[X]^2}{(D + E[X] + F[X]^2)(\langle r\rangle_{max} - \langle r\rangle_{min})}, \qquad [112]$$

We note that $D$, $E$, $F$ are by definition positive, because they are sums of positive weighted spanning trees. We recall that $\langle r\rangle_0 = \frac{A}{D}$, $\langle r\rangle_\infty = \frac{C}{F}$ so by definition of $\langle r\rangle_{min}$, $(A - \langle r\rangle_{min}D)$ and $(C - \langle r\rangle_{min}F)$ are positive coefficients. Furthermore, $(B - \langle r\rangle_{min}E)$ is positive for monotonic outputs, using the negation of non monotonicty condition Eq. [77]. Indeed the conditions for monotonicity can be expressed as,

$$\begin{cases} \text{condition 1:} \quad \langle r\rangle_\infty > \frac{B}{E} \text{ and } \langle r\rangle_0 < \frac{B}{E}, \text{ or} \\ \text{condition 2:} \quad \langle r\rangle_\infty < \frac{B}{E} \text{ and } \langle r\rangle_0 > \frac{B}{E}. \end{cases} \qquad [113]$$

This conditions enforce the fact that $(B - \langle r\rangle_{min}E) > 0$, because since the function is monotonic $\langle r\rangle_{min} = \min(\langle r\rangle_\infty, \langle r\rangle_0)$.

Similarly, the observable $1 - \hat{r}$ is also a rational function with positive coefficients, with the following expression:

$$1 - \hat{r} = \frac{(\langle r\rangle_{max}D - A) + (\langle r\rangle_{max}E - B)[X] + (\langle r\rangle_{max}F - C)[X]^2}{(D + E[X] + F[X]^2)(\langle r\rangle_{max} - \langle r\rangle_{min})}. \qquad [114]$$

With the same arguments than for the previous case, we show that all the coefficients of this rational function in $[X]$ are positive. Last, since $|s(x)| = \left|\dfrac{d\hat{r}}{d\ln x}\right|$, we recover $|s(x)| \leq \frac{1}{2}$ for monotonic outputs.□

Next, we consider nonmotonic responses. Here, we do *not* use the equality Eq. [104] because we can't define observables, which have the form of a positive rational function. Instead, we use the formalism of the coefficients $a$ and $b$. Let us first settle to the case where $a > b > 0$. The extremum of the normalized function $\frac{\langle r\rangle - \langle r\rangle_0}{\langle r\rangle_\infty - \langle r\rangle_0}$ is then a maximum because $a = \frac{dr}{dx}|_{x=0}\frac{1}{\langle r\rangle_\infty - \langle r\rangle_0} > 0$, which implies that the output function first increases and then decreases and therefore reaches a maximum. The minimum of the normalized output is 0 because any increase or decrease of the concentration departing from the value that maximizes the output reduces the output value, by definition. So the minimum is reached at vanishing or infinite concentration. As these values for the normalized output are 0 or 1, we conclude that the minimum is 0. We call $\hat{r} = \frac{\langle r\rangle - \langle r\rangle_0}{\langle r\rangle_\infty - \langle r\rangle_0}$ and show that $\frac{d\hat{r}}{d\ln x} < \frac{1}{2\hat{r}_{max}}$, in order to prove that $s([X]) < \frac{1}{2}$. This is equivalent to showing that $\frac{\hat{r}_{max}}{2}x^4 + (a - b + b\hat{r}_{max})x^3 + (-2 + \hat{r}_{max} + \frac{b^2\hat{r}_{max}}{2})x^2 + (b\hat{r}_{max} - a)x + \frac{\hat{r}_{max}}{2} > 0$, with $\hat{r}_{max} = \frac{ab - 2(1 + \sqrt{1 + a^2 - ab})}{-4 + b^2}$. This is demonstrable by a direct appeal to Mathematica FullSimplify. The case $a < 0$ can be derived similarly.

***J.6. Symbolic derivation of bounds for triply-inflected outputs.*** When the curve has three inflections, the normalized slope has 1/8 for its lower bound and 1/4 for its upper bound. We now demonstrate this behavior analytically.

For the upper bound, we aim to show that $s([X]) < \frac{1}{4}$ for all concentration $[X]$. First we notice that sensitivity with respect to the raw concentration is the same as the sensivity with respect to a renormalized concentration, $s([X]) = s(x)$. This is clear since sensitivity $s$ is a derivative with respect to a logarithmic variable. Substituting our normalized response function in terms of $(a, b)$, the desired upper sensitivity bound is equivalent to the following condition:

$$f(x) = 1 + 2(b - 2a)x + (b^2 - 6)x^2 - 2(b - 2a)x^3 + x^4 > 0. \qquad [115]$$

We note that $f(0) = 1 > 0$ and that $\lim_{x\to\infty} f(x) = +\infty$, so if the function $f$ remains positive on positive values of $x$ the condition Eq. [115] is satisfied. The algebraic conditions assuring three inflection points, as discussed in §I.4, implies $1 + a^2 > ab$, which implies that the function $f$ has no roots.

Indeed, we can prove this quick lemma. Specialize to the case where $b < 2a$. In this case, we study the sign of the polynomial $x(2(b-2a)+(b^2-6)x-2(b-2a)x^2)$. This polynomial vanishes at $x=0$ and at $x_+ = \frac{b^2-6-\sqrt{(b^2-6)^2+16(b-2a)^2}}{4(b-2a)}$. Therefore, this polynomial takes negative values between 0 and $x_+$ and positive for $x > x_+$. The minimal value is taken at $x_{min} = \frac{6-b^2+\sqrt{36+48a^2-48ab+b^4}}{6(2a-b)}$ and lies between 0 and $x_+$. The value at $x_{min}$ of the function $f(x_{min})$ is positive if $1+a^2 > ab$. So in this case $f(x) > 0$.

For the case where $b > 2a$, we study the sign of the polynomial $x^2(b^2-6-2(b-2a)x+x^2)$, which is strictly positive because the associated discriminant of $b^2-6-2(b-2a)x+x^2$ is $\Delta = 16(a^2+\frac{3}{2}-ab)$ is negative if $1+a^2 > ab$.

Now we focus on the lower bound. We note that the maximum of slope is reached either at the 2nd of the 4th inflection, that we called $x_2$ and $x_4$. For we need to prove that it is impossible to have $s(x_2) < \frac{1}{8}$ and $s(x_4) < \frac{1}{8}$ for the same couple $(a,b)$, while satisfying the algebraic condition for three inflection points. Indeed, this condition cannot be satisfied. Therefore, we recover that a lower bound for the maximum of slope of the output over the whole $(a,b)$ space is $\frac{1}{8}$. This is demonstrable by a direct appeal to Mathematica FullSimplify.

### K. Systematic census of effects of pushing on one and two edges.

***K.1. Scaling a single rate constant at a time is identified with a proportional drive.*** The cycle condition relating the ratio of rate constants to the net nonequilibrium driving force $\Delta\mu$ affords us concise expressions for how modifying individual rate parameters induces a net drive. In the main text (or more extensively shortly here in §K), we investigate breaking detailed balance edge-by-edge (while keeping seven rate constants fixed at their default equilibrium values). Say that we are modifying a rate constant $k_{ij}$ away from its default equilibrium value $k_{ij}^{\text{eq.}}$. The cycle condition Eq. [21] implies that

$$\Delta\mu/k_BT = \ln\gamma = \ln\left(\frac{\prod_{i=1}^{N} k_{i,i+1}}{\prod_{i=1}^{N} k_{i+1,i}}\right) \quad [116]$$

$$= \ln\frac{k_{ij}}{k_{ij}^{\text{eq.}}}, \quad [117]$$

since $\gamma = 1$ at equilibrium.

By similar logic, when we adjust two rate constants at once, if they are oriented in the same clockwise or counterclockwise direction in the cycle, then the product of their multiplicative adjustments sets $\gamma$ and therefore $\Delta\mu$. If the rates are instead oriented in opposite directions around the cycle, the ratio of their multiplicative adjustments sets $\gamma$.

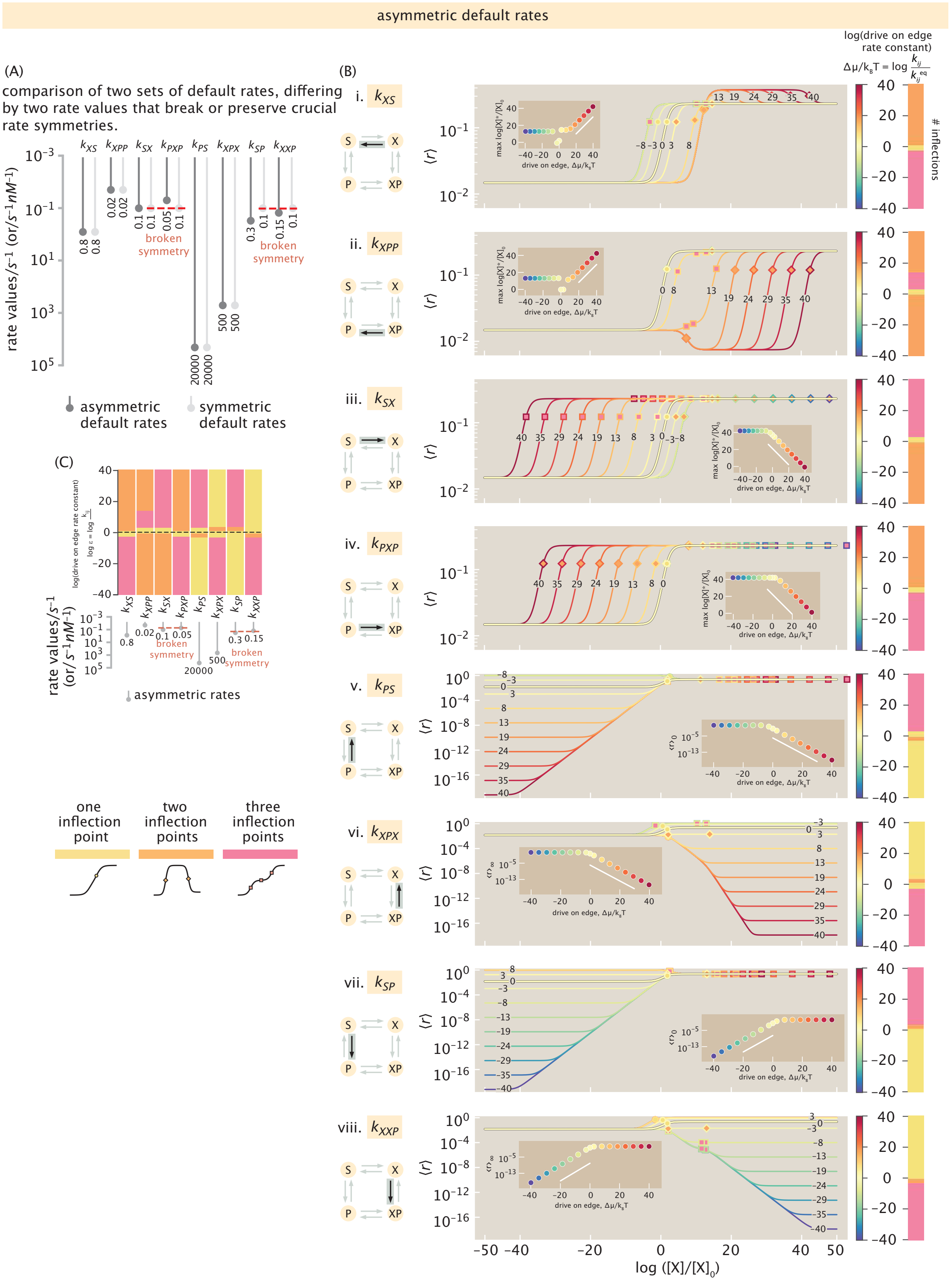

**Fig. S10.** Systematic census of breaking detailed balance, one edge at a time, departing from (slightly) asymmetric default values. These are the parameters used for Figures 3 & 4 of the main text; main text Figure 3 contains two panels of this set. Contrast, panel-by-panel, with the effects of pushing on the same rates, but at different starting values where some symmetries are preserved among the rates, shown in Fig. K.1. In particular, notice that nonmonotonic responses (orange in phase space plots) are significantly less common than in Fig. K.1. (A) Comparison of two sets of starting rates; the sets are the same for four rates, but vary by a factor of less than a few in the other rates, differing in whether critical symmetries are preserved or broken among the rates. (B) The effect of increasing or decreasing each individual rate on the input-output curve, while keeping seven other rates constant. Responses from rate values larger than (or smaller than) at equilibrium are shown in increasingly red (or blue) colors, respectively; curves are also labeled with the numerical values of the net drive that generated them in $k_B T$ units (positive for an increase; negative for a decrease). Each curve's resulting inflection points are marked by yellow, orange, or pink markers, denoting one to three inflection points (respectively), and summarized in the associated one-dimensional (shape phenotypic) phase-diagram with the same colors on the right. (C) Summary of how all eight rates respond to energy expenditure to realize different regulatory shape phenotypes.

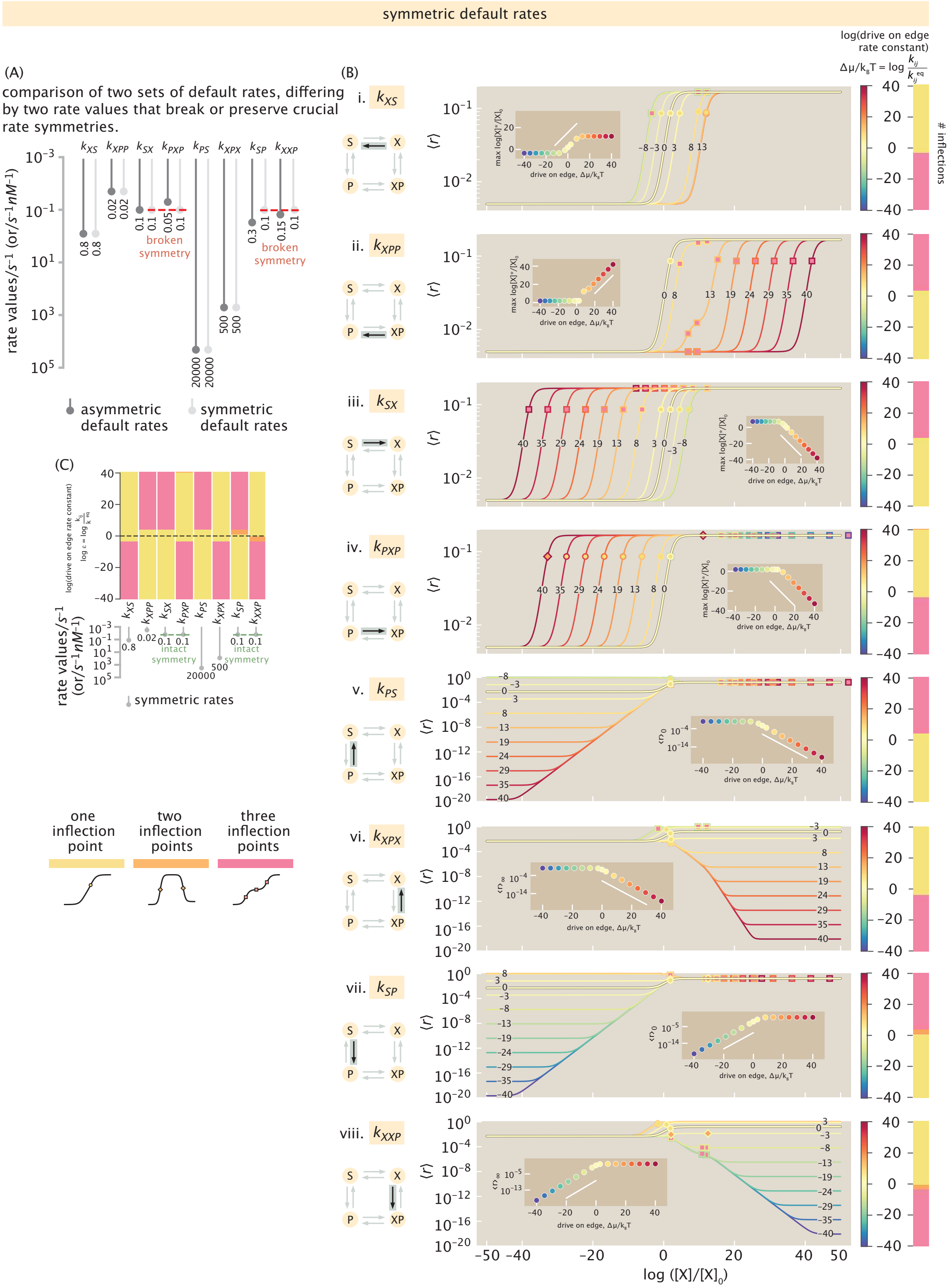

**Fig. S11.** Systematic census of breaking detailed balance, one edge at a time, departing from *symmetric* default values. These are only slightly different than the default parameters used for Figures 3 & 4 of the main text, yet yield richly different behaviors in accessing nonmonotonicity and other phenotypes and illustrate different effects of control. Contrast, panel-by-panel, with Fig. S10. (A) Comparison of two sets of starting rates; the sets are the same for four rates, but vary by a factor of less than a few in the other rates, differing in whether critical symmetries are preserved or broken among the rates. (B) The effect of increasing or decreasing each individual rate on the input-output curve, while keeping seven other rates constant. Responses from rate values larger than (or smaller than) at equilibrium are shown in increasingly red (or blue) colors, respectively; curves are also labeled with the numerical values of the net drive that generated them in $k_BT$ units (positive for an increase; negative for a decrease). Each curve's resulting inflection points are marked by yellow, orange, or pink markers, denoting one to three inflection points (respectively), and summarized in the associated one-dimensional (shape phenotypic) phase-diagram with the same colors on the right. (C) Summary of how all eight rates respond to energy expenditure to realize different regulatory shape phenotypes.

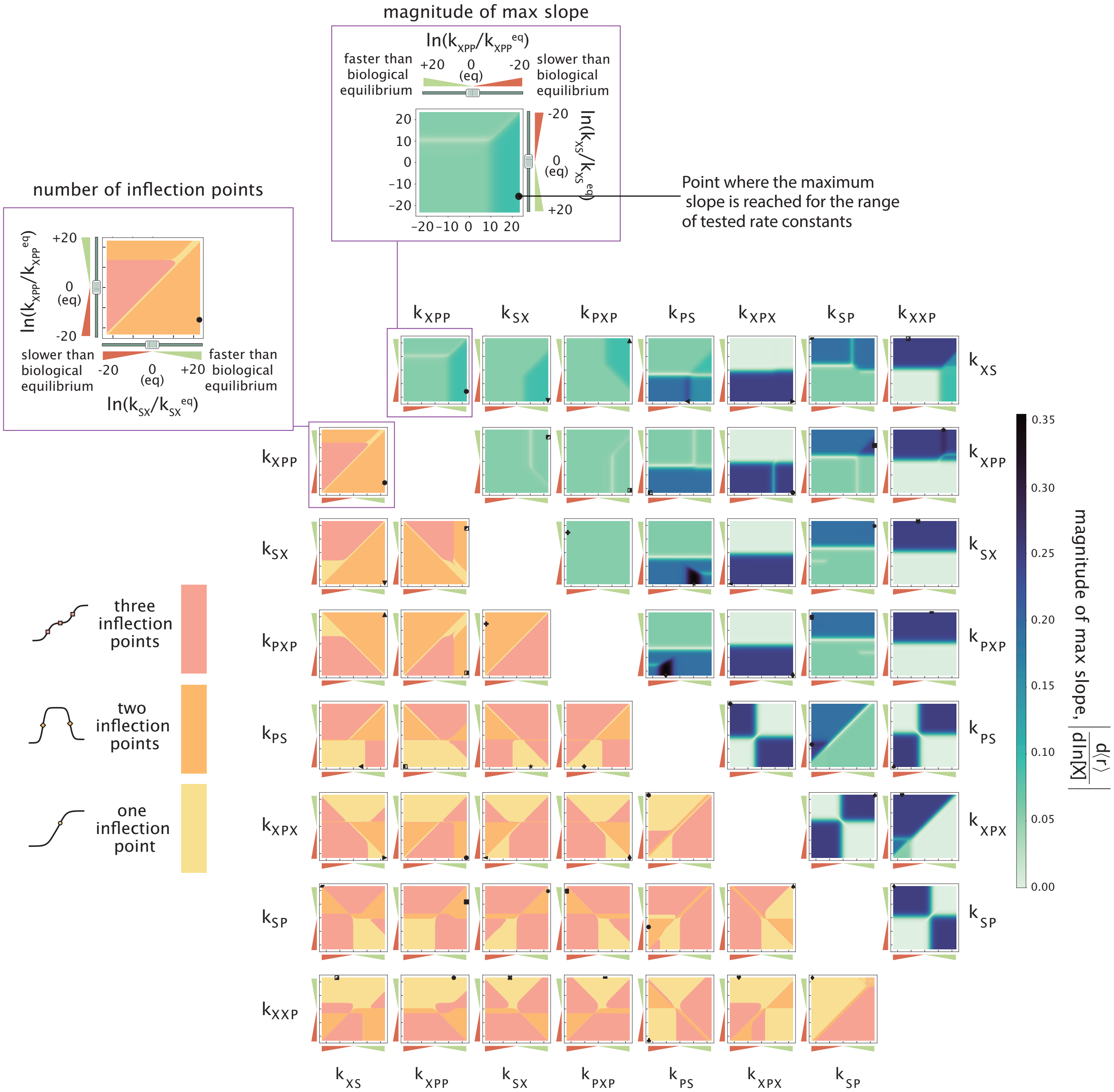

**Fig. S12.** Breaking detailed balance two-edges-at-a-time unlocks wide regions of rate-space where nonmonotonic and triply-inflected phenotypes are observed (lower left triangular matrix). The quantitative properties of the resulting output-curves, such as the slope at first inflection, are also modulated flexibly by these drives (upper right triangular matrix).

**L. Crucial imbalances in rate-constants are required for nonmonotonic responses.** In this section, we derive conditions on the values of rate constant that enable or forbid access to nonmonotonicity. In additional, we find the minimal (nonzero) net drive needed to access nonmonotonicity when kinetic conditions permit. We preview our strategy as follows. First, we translate each of the two conditions guaranteeing nonmonotonicity we found in Eq. [74] from the space of shape parameters $(a, b)$ back into expressions purely in terms of the eight rate constants governing the system. Next, we compel $\gamma$—the product of rate constants in one direction around the cycle divided by the product taken in the opposite direction, whose logarithm gives the net drive, as discussed in §C and §D—to appear in these conditions, by substituting out one of the eight rate constants. We simplify the resulting expressions to surprisingly concise forms that yield minimal drives required to access nonmonotonicity. However, these critical drive values are only defined when precise imbalances among the rates are satisfied, thus establishing sufficient conditions to forbid nonmonotonicity.

1. **We start with the first way to reach nonmonotonicity according to Eq. [74], namely $0 < b < a$:** For algebraic convenience, since this condition specifies the relative value of $a$ and $b$, define $\alpha \equiv 1 - \frac{a}{b}$; this first nonmonotonicity condition is then expressed as $\alpha < 0$. Substituting the definitions of the shape parameters $a, b$ (Eq. [35]) and the definitions of the coefficients $A, B, C, D, E, F$ appropriate for the square graph (Eq. [24]) casts this condition back into the language of rate constants: nonmonotonicity is guaranteed when,

$$\alpha \equiv \frac{([P]k_{SP}+k_{PS})(k_{XXP}(-k_{XS}k_{XPX}k_{PXP}+k_{XPX}k_{XPP}k_{SX}-k_{XPP}k_{SX}k_{PS})+k_{XS}k_{XPX}k_{SP}k_{PXP})}{(k_{XPX}k_{SP}-k_{XXP}k_{PS})([P](k_{SP}k_{PXP}([P]k_{XXP}+k_{XS}+k_{XPX})+k_{XXP}k_{XPP}k_{SX})+k_{SX}k_{PS}([P]k_{XXP}+k_{XPX}+k_{XPP})+k_{XS}k_{XPX}k_{PXP})} < 0. \quad [118]$$

Next, we simplify by positive factors, and use fact that $\frac{1}{k_{XPX}k_{SP}-k_{XXP}k_{PS}}$ and $k_{XPX}k_{SP} - k_{XXP}k_{PS}$ have the same sign. Since we want to force $\gamma \equiv \frac{k_{XXP}k_{XPP}k_{PS}}{k_{SX}k_{XS}k_{XPX}k_{SP}k_{PXP}}$ to appear to comment on energetic drive, we choose a rate constant to express in terms of $\gamma$ and the other seven rates. Without loss of generality, we choose to replace $k_{SX}$ by $k_{SX} = \gamma \frac{k_{XS}k_{XPX}k_{SP}k_{PXP}}{k_{XXP}k_{XPP}k_{PS}}$. These manipulations convert Eq. 118 into the much more succinct and revealing form,

$$\left(\frac{k_{SP}}{k_{PS}} - \frac{k_{XXP}}{k_{XPX}}\right)\left(1 - \frac{k_{XXP}}{k_{SP}} - \gamma\left(1 - \frac{k_{XPX}}{k_{PS}}\right)\right) < 0. \quad [119]$$

Now, we solve for possible values of $\gamma$, under the mathematical constraints that $\gamma$ must itself remain positive (that is, nonnegative because it is a ratio of positive rate constants, and greater than zero because we know nonmonotonic outputs cannot occur at detailed balance). We could solve this condition Eq. [119] by hand, case-by-case; but for ease we use a call to *Mathematica*'s `Reduce` command over $\gamma$ on the `PositiveReals`, while enforcing assumptions that all rates are positive. This analysis generates all the specific possible conditions where $\gamma$ is defined and satisfies this nonmonotonicity criterion; these transpire to be,

$$\begin{cases} 0 < \gamma < \frac{k_{PS}(k_{SP}-k_{XXP})}{k_{SP}(k_{PS}-k_{XPX})} & \text{and} \begin{cases} k_{SP} < k_{XXP} & \text{and } k_{PS}k_{XXP} < k_{SP}k_{XPX} \text{ or,} \\ k_{SP} > k_{XXP} & \text{and } k_{PS}k_{XXP} < k_{SP}k_{XPX}. \end{cases} \quad \text{or ,} \\ \gamma > \frac{k_{PS}(k_{SP}-k_{XXP})}{k_{SP}(k_{PS}-k_{XPX})} & \text{and} \begin{cases} k_{XPX} < k_{PS} < \frac{k_{SP}k_{XPX}}{k_{XXP}} \text{ or,} \\ \frac{k_{SP}k_{XPX}}{k_{XXP}} < k_{PS} < k_{XPX} \text{ and } k_{SP} < k_{XXP}. \end{cases} \end{cases} . \quad [120]$$

Clearly this panoply of logical conditions is intricate. To interpret and summarize these conditions, we define some notation for the constituent kinetic conditions, which often have physical interpretations:

- First, recall that the conditions for the transcription factor to be an overall repressor or activator are simply given by,

$$\begin{cases} \text{activation, } A \equiv \frac{k_{SP}}{k_{PS}} < \frac{k_{XXP}}{k_{XPX}} \\ \text{repression, } R \equiv \frac{k_{SP}}{k_{PS}} > \frac{k_{XXP}}{k_{XPX}} \end{cases} . \quad [121]$$

- Next, for concision, denote the following pairwise conditions among rates as,

$$\begin{cases} c_1 \equiv \frac{k_{XPX}}{k_{PS}} > 1 \\ c_2 \equiv \frac{k_{XPX}}{k_{PS}} < 1 \\ c_3 \equiv \frac{k_{XXP}}{k_{SP}} > 1 \\ c_4 \equiv \frac{k_{XXP}}{k_{SP}} < 1 \end{cases} . \quad [122]$$

  (Note that $c_1$ and $A$ imply $c_3$; $c_2$ and $R$ imply $c_4$; $c_4$ and $A$ imply $c_2$; and last, $c_3$ and $R$ imply $c_1$.)

- Recalling that the net drive present in the cycle is given by $\Delta\mu = k_B T \ln \gamma$ (see §D), we now identify two constituent requirements for nonmonotonicity from those of Eq. [120], expressed in terms of $\Delta\mu$. We denote them $c_+$ and $c_-$, because satisfying them respectively reflects a clockwise stationary flux and a counterclockwise flux while allowing nonmonotonicity; denote their logical union the condition $c$. These are defined as,

$$c \equiv \begin{cases} c_+(k_{XXP}, k_{SP}, k_{XPX}, k_{PS}) \equiv (\Delta\mu > 0) \text{ and } ((c_1 \text{ and } A) \text{ or } (c_2 \text{ and } R)), \text{ or,} \\ c_-(k_{XXP}, k_{SP}, k_{XPX}, k_{PS}) \equiv (\Delta\mu < 0) \text{ and } ((c_4 \text{ and } A) \text{ or } (c_3 \text{ and } R)) \end{cases} . \quad [123]$$

Finally, we use all this notation to interpret Eq. [120] as saying that when rate constants satisfy the necessary conditions $c(k_{XXP}, k_{SP}, k_{XPX}, k_{PS})$ (Eq. [123]), a minimum critical drive $\Delta\mu_1$ exists, defined by

$$\boxed{\Delta\mu_1 = k_B T \left| \ln \frac{\frac{k_{XXP}}{k_{SP}} - 1}{\frac{k_{XPX}}{k_{PS}} - 1} \right|}. \tag{124}$$

That is, when the drive $\Delta\mu$ exceeds this $\Delta\mu_1$ in magnitude under the right preexisting rate conditions,

$$|\Delta\mu| > \Delta\mu_1, \tag{125}$$

responses are nonmonotonic.

2. **Next, we turn to the second way to reach nonmonotonicity according to Eq. [74], namely $a < 0$:** Analogously to how we treated the first nonmonotonicity condition, we translate $a < 0$ to $1 - \alpha < 0$ and substitute rate constants into the definitions, expressing the present nonmonotonicity condition as

$$\frac{([P]k_{XXP}+k_{XPX})(-k_{XPX}k_{SP}(k_{XS}+[P]k_{SP})k_{PXP}+(k_{XXP}k_{XPP}k_{SX}-(k_{XPX}+k_{XPP})k_{SX}k_{SP}+([P]k_{XXP}+k_{XS})k_{SP}k_{PXP})k_{PS}+k_{XXP}k_{SX}k_{PS}^2)}{(k_{XPX}k_{SP}-k_{XXP}k_{PS})(k_{XS}k_{XPX}k_{PXP}+[P](k_{XXP}k_{XPP}k_{SX}+([P]k_{XXP}+k_{XS}+k_{XPX})k_{SP}k_{PXP})+([P]k_{XXP}+k_{XPX}+k_{XPP})k_{SX}k_{PS})}<0 \tag{126}$$

Simplifying by the positive terms; noticing that $\frac{1}{k_{XPX}k_{SP}-k_{XXP}k_{PS}}$ and $k_{XPX}k_{SP} - k_{XXP}k_{PS}$ have the same sign; and replacing $k_{SX}$ by $\gamma\frac{k_{XS}k_{XPX}k_{SP}k_{PXP}}{k_{XXP}k_{XPP}k_{PS}}$ recasts this condition as,

$$\left(\frac{k_{SP}}{k_{PS}} - \frac{k_{XXP}}{k_{XPX}}\right)\left(1 - \left(\frac{k_{PS}}{k_{XPX}} + \frac{[P]k_{XXP}k_{PS}}{k_{XS}k_{XPX}} - \frac{[P]k_{SP}}{k_{XS}}\right) - \gamma\left(1 - \left(\frac{k_{XPX}k_{SP}}{k_{XXP}k_{XPP}} + \frac{k_{SP}}{k_{XXP}} - \frac{k_{PS}}{k_{XPP}}\right)\right)\right) < 0\ . \tag{127}$$

Now, as before, we solve for the values of $\gamma$ that are positive, real, and compatible with this condition. Since the resulting specific conditions are most intrepretable when expressed directly in terms of $\Delta\mu = k_B T \ln\gamma$, we report them directly in this variable. To do so, we again define some notation for governing subconditions that materialize as follows.

- Denote the following logical conditions with the shorthand $d_i$,

$$\begin{cases} d_1 \equiv (k_{XPX} + k_{XPP})k_{SP} < k_{XXP}(k_{XPP} + k_{PS}) \\ d_2 \equiv (k_{XPX} + k_{XPP})k_{SP} > k_{XXP}(k_{XPP} + k_{PS}) \\ d_3 \equiv ([P]k_{XXP} + k_{XS})k_{PS} < k_{XPX}(k_{XS} + [P]k_{SP}) \\ d_4 \equiv ([P]k_{XXP} + k_{XS})k_{PS} > k_{XPX}(k_{XS} + [P]k_{SP}) \end{cases}. \tag{128}$$

- Then nonmonotonicity is possible, and rates induce clockwise (+) and counterclockwise (-) steady-state fluxes, respectively, when either of the conditions $d_+$ and $d_-$ are satisfied,

$$d \equiv \begin{cases} d_+(k_{XXP}, k_{SP}, k_{XPX}, k_{PS}, k_{XPP}, k_{XS}) \equiv (\Delta\mu > 0 \text{ and } k_{XPP} > k_{XPX}k_{PS}\left|\frac{\frac{k_{SP}}{k_{PS}} - \frac{k_{XXP}}{k_{XPX}}}{k_{XXP} - k_{SP}}\right| \text{ and } ((c_3 \text{ and } R) \text{ or } (c_4 \text{ and } A)) \\ d_-(k_{XXP}, k_{SP}, k_{XPX}, k_{PS}, k_{XPP}, k_{XS}) \equiv (\Delta\mu < 0 \text{ and } k_{XS} > k_{XPX}k_{PS}\left|\frac{\frac{k_{SP}}{k_{PS}} - \frac{k_{XXP}}{k_{XPX}}}{[P](k_{XPX} - k_{PS})}\right| \text{ and } ((c_1 \text{ and } A) \text{ or } (c_2 \text{ and } R)), \end{cases} \tag{129}$$

  where we have denoted their logical union $d$.
  We also remark that an alternative, equivalent way of expressing Eq. [129] is as follows,

$$d \equiv \begin{cases} d_+(k_{XXP}, k_{SP}, k_{XPX}, k_{PS}, k_{XPP}, k_{XS}) = (\Delta\mu > 0) \text{ and } ((d_1 \text{ and } c_3 \text{ and } R) \text{ or } (d_2 \text{ and } c_4 \text{ and } A)) \\ d_-(k_{XXP}, k_{SP}, k_{XPX}, k_{PS}, k_{XPP}, k_{XS}) = (\Delta\mu < 0) \text{ and } ((d_3 \text{ and } c_1 \text{ and } A) \text{ or } (d_4 \text{ and } c_2 \text{ and } R)). \end{cases} \tag{130}$$

This notation allows us to interpret Eq. [127] as saying that when rates satisfy the conditions $d(k_{XXP}, k_{SP}, k_{XPX}, k_{PS}, k_{XPP}, k_{XS})$, there is a minimal drive $\Delta\mu_2$ past which nonmonotonicity is activated,

$$|\Delta\mu| > \Delta\mu_2, \tag{131}$$

where

$$\boxed{\Delta\mu_2 = k_B T \left| \ln \frac{\frac{k_{XXP}}{k_{XPX}} - \frac{k_{SP}}{k_{PS}} + \frac{k_{XS}}{k_{XPX}P} - \frac{k_{XS}}{k_{PS}P}}{\frac{k_{XPX}}{k_{XXP}} + \frac{k_{XPP}}{k_{XXP}} - \frac{k_{XPP}}{k_{SP}} - \frac{k_{PS}}{k_{SP}}} \frac{k_{XPP}}{k_{XS}} \frac{k_{SP}}{k_{PS}} \right|}. \tag{132}$$

***L.1. Minimum drive to reach nonmonotonic phenotypes.*** In this section, we investigate analytical lessons from our preceding analysis that comment on the behaviors we encountered in our numerical analyses driving two edges in Fig. S12 and Fig. 4 of the main text.

When they are mathematically defined, the critical drive values $\Delta\mu_1$ and $\Delta\mu_2$ are the minimum inputs of drive required to convert a monotonic output to a nonmonotonic output. It is worth remarking that once those critical values are exceeded, nonmonotonicity can persist only for a finite range of drive, because the underlying kinetic conditions—namely, $c$ (Eq. [123]) or $d$ (Eq. [129])—that enable the critical drives to exist are not always satisfied. However, so long as at least one of $c$ or $d$ is always satisfied, $\Delta\mu_1$ and/or $\Delta\mu_2$ are rigorous values for the critical drive the system must maintain to create nonmonotonicity.

Now, we specialize to the case where we may control just one of the four rate constants $(k_{XXP}, k_{SP}, k_{XPX}, k_{PS})$, in addition to some other arbitrarily chosen one. To be concise, denote $x_1 = \frac{k_{XXP}}{k_{SP}}$ and $x_2 = \frac{k_{XPX}}{k_{PS}}$. The first way to access nonmonotonicity is when condition (Eq. [123]) is satisfied, allowing $\Delta\mu_1$ to exist. As long as $x_1 \neq x_2$, this condition $c$ may also be expressed as,

$$c(k_{XXP}, k_{SP}, k_{XPX}, k_{PS}) = \begin{cases} x_1 > 1 \text{ and } x_2 > 1 & \text{or,} \\ x_1 < 1 \text{ and } x_2 < 1. \end{cases} \qquad [133]$$

Under this condition, if $x_1 \to x_2$, $\Delta\mu_1 \to 0$ non-monotonicity is reached for any finite drive. When at detailed balance using our estimated biological starting rates, the default values of these governing ratios are $x_{1eq} < 1$ and $x_{2eq} < 1$. Accordingly, if we tune one of the four rate constants that define $x_1$ or $x_2$, we can approach the limit where $x_1 \to x_{2eq} < 1$ or $x_2 \to x_{1eq} < 1$, while preserving the necessary conditions for $\Delta\mu_1$ to exist and the response to be nonmonotonic. To compensate, the additional rate constant being tuned can then be adjusted to ensure that asymptotically-little energy is spent, $\gamma \to 1$. This protocol would ensure that an asymptotically-small adjustment of rate constants from such default values would unlock a nonmonotonic output at any nonzero drive. This special starting point is unique for a given pair of rate constants that satisfy this condition, because there are two unknowns (the two rate constants) and two asymptotic equations, namely,

$$\begin{cases} x_1 = \frac{k_{XXP}}{k_{SP}} \to \frac{k_{XPX}}{k_{PS}} = x_2 \\ \gamma \equiv \frac{k_{SX} k_{X,XP} k_{XP,P} k_{PS}}{k_{XS} k_{XP,X} k_{P,XP} k_{SP}} \to 1 \end{cases}. \qquad [134]$$

For the remaining six pairs of rate constants that do not include the four rates that define $x_1$ and $x_2$, the limit of the minimal drive needed to reach nonmonotonicity is a finite value. In fact, this value is same minimum drive needed when tuning only one of the two edges among a pair. We call this value $\Delta\mu_0$. Indeed, with the rates at equilibrium we chose, the minimal drive for the output to be non monotonic when energy is injected along one of the four rate constants $(k_{PXP}, k_{XPP}, k_{XS}, k_{SX})$ is the same (also valued at $\Delta\mu_0$).

***L.2. Conditions that suffice to forbid nonmonotonicity.*** Now, consider the cases where neither $\Delta\mu_1$ nor $\Delta\mu_2$ is defined. That is to say, when non-monotonicity cannot be achieved for any input of drive on the system. From the converse of the condition $c$ (Eq. [123]), we can deduce that as soon as one of the following conditions is not satisfied, $\Delta\mu_1$ is not defined,

$$\begin{cases} k_{XPX} = k_{PS} & \text{or,} \\ k_{XXP} = k_{SP} & \text{or,} \\ c_1 \text{ and } c_4 & \text{or,} \\ c_2 \text{ and } c_3. \end{cases} \qquad [135]$$

Substituting the meanings of the subconditions $c_1$ through $c_4$ expresses these conditions guaranteeing monotonicity as,

$$\boxed{\begin{cases} k_{XPX} = k_{PS}, \text{ or} \\ k_{XXP} = k_{SP}, \text{ or} \\ k_{XXP} > k_{SP} \text{ and } k_{XPX} < k_{PS}, \text{ or} \\ k_{XPX} > k_{PS} \text{ and } k_{XXP} < k_{SP} \end{cases}}. \qquad [136]$$

For instance, some of the conditions in Eq. [135] immediately suffice to forbid nonmonotonicity via $\Delta\mu_1$ because the argument of the logarithm in $\Delta\mu_1$'s definition becomes negative. Evaluating the second possible route to reach nonmonotonicity, via $\Delta\mu_2$ and its prerequisite condition $d$ (Eq. [129]), we see the same conditions above suffice to forbid its mathematical definition. In summary, if any of the conditions in Eq. [136] are satisfied, the response function must remain monotonic, even for any nonequilibrium driving on the system.

Notice that these conditions Eq. [136] depend only on four rate constants: the binding and and unbinding rates of the polymerase. These are the same four rate constants that fix both the leakiness and saturation. We illustrate these impacts of tuning ratios of these four rate constants in Fig. S13.

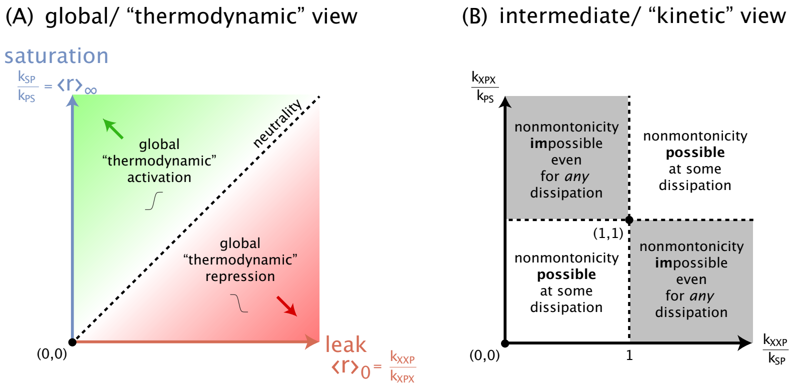

**Fig. S13.** Nonmonotonic input-output curves are impossible even under any dissipation when certain relationships are obeyed by rate constants. In particular, while $k_{SP}/k_{PS}$ and $k_{XXP}/k_{kXPX}$ set whether the transcription factor is globally an activator or repressor (showing a saturation larger (smaller) than the leak, respectively; panel (A)), it is the ratios $k_{XPX}/k_{PS}$ and $k_{XXP}/k_{SP}$ that set whether the curve can ever be nonmonotonic (panel (B)).

As discussed briefly in the main text, some biophysical contexts may, by default, satisfy some of the conditions Eq. [136] guaranteeing nonmonotonic responses. For instance, under the classical assumption that the binding rate of the polymerase is purely diffusion-limited, its on rate would not depend on whether the transcription factor is already bound to the genome or not, enforcing $k_{XXP} = k_{SP}$ and hence forbidding nonmonotonicity by default, even for any drive or modulation of the other rate constants. Manifesting nonmonotonicity departing from these default rates would then require energy investment to break this rate symmetry. This pivotal constraint is plausibly relievable by diverse modes of transcriptional regulation, but emphasizes the privileged roles that some ratios of rate constants have in determining the flexibility of output responses. We illustrate two such symmetries, with different default biological plausibility, in Fig. S14.

For biological reasons, other pairs of rate constants of the system could be equal. Indeed, if the binding of any molecules is only limited by diffusion, the on rates of the transcription factor should also be equal. We observe, and Eq. 5 of the main text reports, that the only equalities between pairs of rate constants that forbid non-monotonicity are the on- or off- rates of the polymerase. For instance, the equality between rates of the transcription factor does not forbid the access to non-monotonicity.

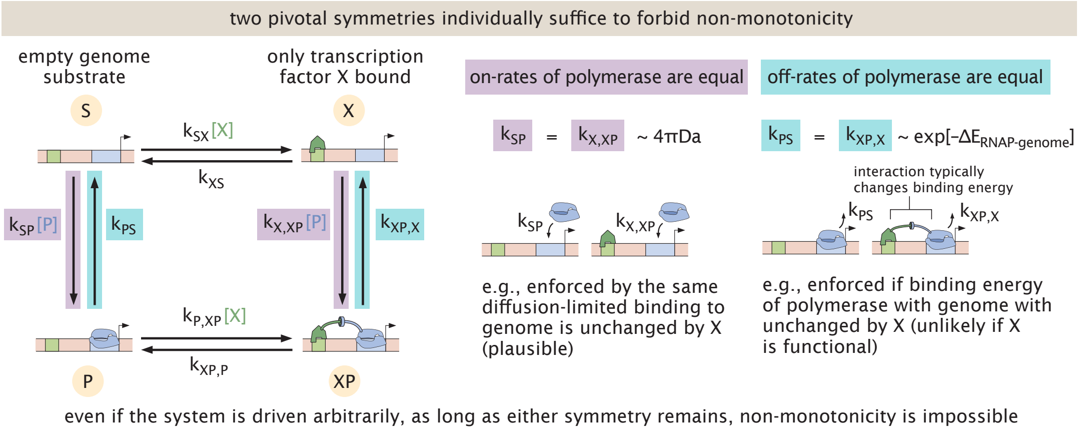

**Fig. S14.** Physical examples of critical symmetries among rate constants that suffice to forbid nonmontonic responses.

**M. Implications of critical symmetry conditions for widespread numerical screens.** A common, *a priori* reasonable, tactic to confront the explosion in the number of parameters of kinetic models that can accommodate nonequilibrium (relative to the fewer energetic paramaters of equilibrium) is to restrict parameters within certain ranges or under simplifying functional constraints. These constraints help grapple with the reality that each additional parameter implies an exponential increase in the number of, for example, combinatorially-investigated samples of a model. However, our analytical results highlight how imposing such constraints among parameters can unexpectedly collapse the complexity achievable by a kinetic model into a restricted set of output behaviors.

For example, Lammers, Flamholz, & Garcia (9) recently performed a study of how energetic and kinetic parameters affect the rate at which information is transferred from inputs to transcriptional outputs in a generic model of transcriptional activation inspired by the Monod-Wyman-Changeux model. This study imposed an apparently-benign constraint of parameters intuitively motivated by assuming that a transcription factor accomplishes activation. Specifically, Lammers *et al.* reasoned that if the transcription factor increases the rate at which the system switches between transcriptionally OFF and ON states (relative to this rate without the transcription factor), as encoded by an interaction term they call $\eta_{ab} > 1$, but also *decreases* the complementary switching rate from OFF to ON (encoded by another interaction term $\eta_{ib} < 1$), then the presence of the transcription factor activates transcription (namely, increases the probability of being in a transcriptionally ON state) ((9) and personal communication). In fact, however, this ($\eta_{ab} > 1$ and $\eta_{ib} < 1$) constraint is sufficient, but *not* necessary, for activation. Instead, a looser constraint—merely that the transcription factor makes the ON to OFF rate slower overall than the OFF to ON rate ($\eta_{ib} < \eta_{ab}$)—is the minimal condition adequate for activation. (Thus, a transcription factor can still ultimately activate transcription even when it increases or decreases both transcriptionally OFF-to-ON and ON-to-OFF rates, as long the former still exceeds the latter.) Further, surprisingly, our analytic reasoning establishes the stricter ($\eta_{ab} > 1$ and $\eta_{ib} < 1$) constraints previously assumed by Lammers & colleagues are precisely among those that suffice to *forbid* nonmonotonic output responses, even for any energy expenditure (see Eq. 5 of the main text, and also Fig. S15).

More specifically, a transcription factor is an activator when the "leak" transcriptional output $\langle r\rangle_0$ without any transcription factor is less than the "saturation" output $\langle r\rangle_\infty$ at a saturating (say infinite) concentration of transcription factor. As discussed earlier in §G.2, when the transcription factor is completely absent, the system cannot be found in any microstate that invokes it, collapsing four states into just the two states devoid of transcription factor. Similarly, when the transcription factor concentration is infinite, the system is never found in the two microstates without the transcription factor, again admitting an (orthogonal) two-state description. In the language of the model of Lammers, Flamholz, & Garcia, this implies that the leak $\langle r\rangle_0$ is set by a competition between an ON state with probability $p_{\textcircled{3}}$ and an OFF state with probability $p_{\textcircled{0}}$ (see Fig. S15A, right), where the former transitions to the latter at rate $k_i$ and the latter transitions to the former at rate $k_a$, just as in §G.2. Hence,

$$\langle r\rangle_0 = rp_{\textcircled{3}} = r\frac{k_a}{k_a + k_i} = r\frac{1}{1+\frac{k_i}{k_a}}. \quad [137]$$

Conversely, at saturating transcription factor, the output is set by a competition between an ON state with probability $p_{\textcircled{2}}$ and an OFF state with probability $p_{\textcircled{2}}$ that respectively transition between each other at rates $\eta_{ib}k_i$ and $\eta_{ab}k_a$. So the saturation is

$$\langle r\rangle_\infty = rp_{\textcircled{2}} = r\frac{\eta_{ab}k_a}{\eta_{ab}k_a + \eta_{ib}k_i} = r\frac{1}{1+\frac{\eta_{ib}}{\eta_{ab}}\frac{k_i}{k_a}}. \quad [138]$$

Overall, these expressions indicate that the transcription factor is a net activator, $\langle r\rangle_0 < \langle r\rangle_\infty$, exactly when $\frac{\eta_{ib}}{\eta_{ab}}\frac{k_i}{k_a} < \frac{k_i}{k_a}$, or namely

$$\text{activation: } \boxed{\frac{\eta_{ib}}{\eta_{ab}} < 1}. \quad [139]$$

Importantly, this is a *looser* condition than that simultaneously ($\eta_{ib} < 1$ and $\eta_{ab} > 1$), as assumed by Lammers, Flamholz, & Garcia (9).

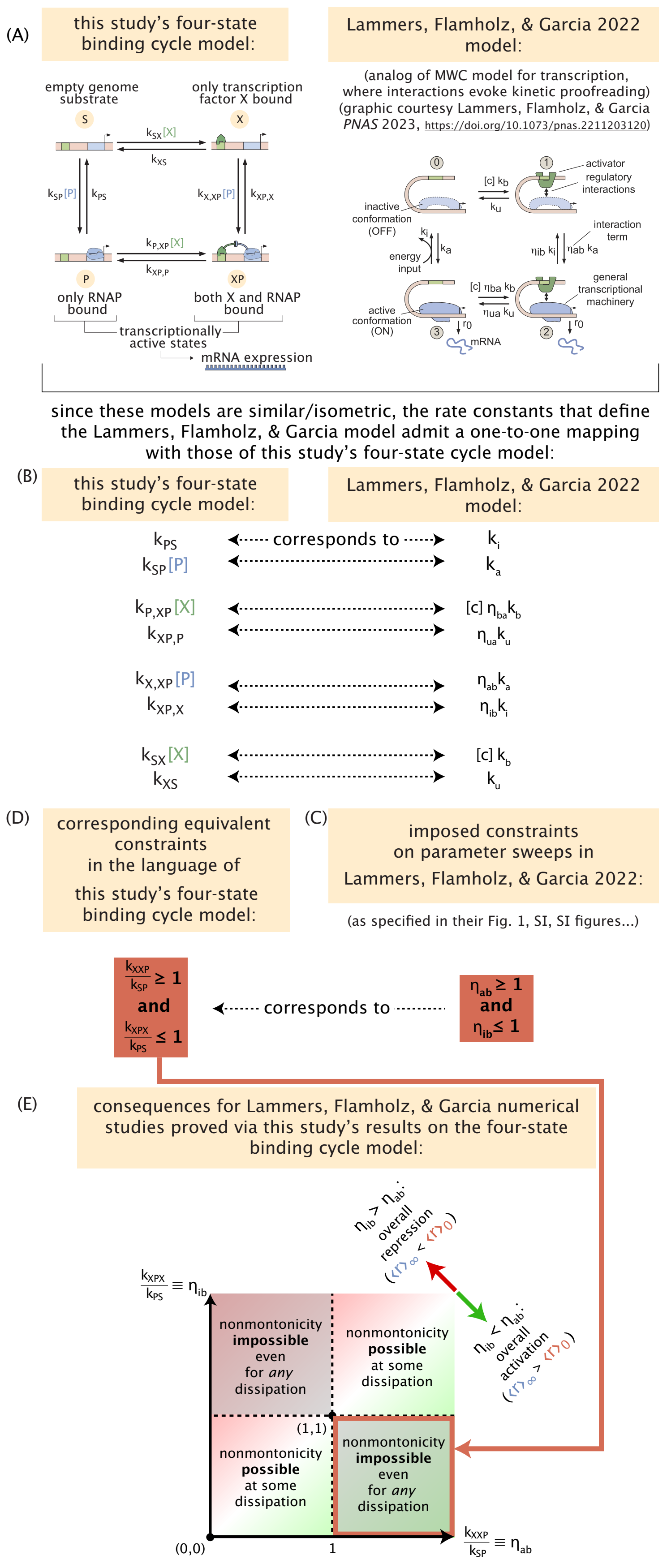

**Fig. S15.** The simple, analytic conditions we found among rate constants that permit or forbid nonmonotonicity in this four-state system imply strong consequences for the result of large numerical studies by other investigations. Specifically, our conditions imply that a recent study on the relationship between dissipative parameters and the accumulation of transcriptional information (9) admits a hidden/nonobvious restriction implying that all of their input-output curves must be monotonic, for any dissipation.

### References

1. JA Owen, JM Horowitz, Size limits the sensitivity of kinetic schemes. *Nat. Commun.* **14**, 1280 (2023).
2. R Grah, B Zoller, G Tkačik, Nonequilibrium models of optimal enhancer function. *Proc. Natl. Acad. Sci.* **117**, 31614–31622 (2020).
3. M Morrison, M Razo-Mejia, R Phillips, Reconciling kinetic and thermodynamic models of bacterial transcription. *PLoS computational biology* **17**, e1008572 (2021).
4. TL Forcier, et al., Measuring cis-regulatory energetics in living cells using allelic manifolds. *Elife* **7**, e40618 (2018).
5. H Qian, Phosphorylation energy hypothesis: open chemical systems and their biological functions. *Annu. Rev. Phys. Chem.* **58**, 113–142 (2007).
6. C Hueschen, R Phillips, *The Restless Cell: Continuum Theories of Living Matter.* (Princeton University Press), (2023).
7. WT Ireland, et al., Deciphering the regulatory genome of escherichia coli, one hundred promoters at a time. *Elife* **9**, e55308 (2020).
8. M Rydenfelt, HG Garcia, RS Cox III, R Phillips, The influence of promoter architectures and regulatory motifs on gene expression in escherichia coli. *PLoS one* **9**, e114347 (2014).
9. NC Lammers, AI Flamholz, HG Garcia, Competing constraints shape the nonequilibrium limits of cellular decision-making. *Proc. Natl. Acad. Sci.* **120**, e2211203120 (2023).
10. HC Nelson, RT Sauer, Lambda repressor mutations that increase the affinity and specificity of operator binding. *Cell* **42**, 549–558 (1985).
11. R Milo, P Jorgensen, U Moran, G Weber, M Springer, Bionumbers—the database of key numbers in molecular and cell biology. *Nucleic acids research* **38**, D750–D753 (2010).
12. AD Riggs, S Bourgeois, M Cohn, The lac represser-operator interaction: Iii. kinetic studies. *J. molecular biology* **53**, 401–417 (1970).
13. E Marklund, et al., Sequence specificity in dna binding is mainly governed by association. *Science* **375**, 442–445 (2022).
14. P Hammar, et al., Direct measurement of transcription factor dissociation excludes a simple operator occupancy model for gene regulation. *Nat. genetics* **46**, 405–408 (2014).
15. R Milo, R Phillips, *Cell biology by the numbers.* (Garland Science), (2015).
16. J Elf, GW Li, XS Xie, Probing transcription factor dynamics at the single-molecule level in a living cell. *Science* **316**, 1191–1194 (2007).
17. J Elf, GW Li, XS Xie, Probing transcription factor dynamics at the single-molecule level in a living cell. *Science* **316**, 1191–1194 (2007).
18. AP Singh, et al., 3d protein dynamics in the cell nucleus. *Biophys. journal* **112**, 133–142 (2017).
19. L Bintu, et al., Transcriptional regulation by the numbers: models. *Curr. opinion genetics & development* **15**, 116–124 (2005).
20. M Razo-Mejia, et al., Tuning transcriptional regulation through signaling: a predictive theory of allosteric induction. *Cell Syst.* **6**, 456–469 (2018).
21. L Xu, et al., Average gene length is highly conserved in prokaryotes and eukaryotes and diverges only between the two kingdoms. *Mol. biology evolution* **23**, 1107–1108 (2006).
22. AJ Meyer, TH Segall-Shapiro, E Glassey, J Zhang, CA Voigt, Escherichia coli "marionette" strains with 12 highly optimized small-molecule sensors. *Nat. chemical biology* **15**, 196–204 (2019).
23. O Shoval, et al., Fold-change detection and scalar symmetry of sensory input fields. *Proc. Natl. Acad. Sci.* **107**, 15995–16000 (2010).
24. D Curtiss, Recent extentions of descartes' rule of signs. *Annals Math.* pp. 251–278 (1918).
25. A Goldbeter, DE Koshland Jr, An amplified sensitivity arising from covalent modification in biological systems. *Proc. Natl. Acad. Sci.* **78**, 6840–6844 (1981).
26. Y Tu, The nonequilibrium mechanism for ultrasensitivity in a biological switch: Sensing by maxwell's demons. *Proc. Natl. Acad. Sci.* **105**, 11737–11741 (2008).
27. H Tran, et al., Precision in a rush: Trade-offs between reproducibility and steepness of the hunchback expression pattern. *PLoS computational biology* **14**, e1006513 (2018).
